# Leveraging fine-scale population structure reveals conservation in genetic effect sizes between human populations across a range of human phenotypes

**DOI:** 10.1101/2023.08.08.552281

**Authors:** Sile Hu, Lino A. F. Ferreira, Sinan Shi, Garrett Hellenthal, Jonathan Marchini, Daniel J. Lawson, Simon R. Myers

**Affiliations:** Department of Statistics, University of Oxford, Oxford, UK; Wellcome Centre for Human Genetics, University of Oxford, Oxford, UK; Human Genetics Centre of Excellence, Novo Nordisk Research Centre Oxford, Oxford, UK; Department of Genetics, Evolution and Environment, University College London, London, UK; UCL Genetics Institute, University College London, London, UK; Regeneron Genetic Center, Tarrytown, New York, USA; Department of Statistical Science, School of Mathematics, University of Bristol, Bristol, UK; MRC Integrative Epidemiology Unit, Population Health Sciences, Bristol Medical School, University of Bristol, Bristol, UK

## Abstract

An understanding of genetic differences between populations is essential for avoiding confounding in genome-wide association studies (GWAS) and understanding the evolution of human traits. Polygenic risk scores constructed in one group perform poorly in highly genetically-differentiated populations, for reasons which remain controversial. We developed a statistical ancestry inference pipeline able to decompose ancestry both within and between countries, and applied it to the UK Biobank data. This identifies fine-scale patterns of genetic relatedness not captured by standard and widely used principal components (PCs), and allows fine-scale population stratification correction that removes both false positive and false negative associations for traits with geographic correlations. We also develop and apply ANCHOR, an approach leveraging segments of distinct ancestries within individuals to estimate similarity in underlying causal effect sizes between groups, using an existing PGS. Applying ANCHOR to >8000 people of mixed African and European ancestry, we demonstrate that estimated causal effect sizes are highly similar across these ancestries for 26 of 29 quantitative molecular and non-molecular phenotypes (mean correlation 0.98 +/-0.08), providing evidence that gene-environment and gene-gene interactions do not play major roles in the poor prediction of European-ancestry PRS scores in African populations for these traits, contradicting previous findings. Instead our results provide optimism that shared causal mutations operate similarly in different groups, focussing the challenge of improving GWAS “portability” between groups on joint fine-mapping.

## Introduction

Genome-wide association studies (GWAS) have illuminated many genetic influences on a huge range of complex human phenotypes, revealed to be regulated by hundreds, or thousands, of distinct loci with small effect sizes. For some traits, such as height, sample sizes have become sufficiently large that within the most studied groups, typically European populations, a substantial fraction of trait heritability is now explained via polygenic scores (PGS), which add up effects across many SNPs genome-wide^1^. Because the true causal variants are unknown, SNPs contributing to these PGS are often expected not to influence a trait directly, but rather correlate with (“tag”) a true causal mutation, at least in the population in which the GWAS producing the PGS was conducted. The most serious confounder in GWAS studies is genetic stratification between groups. Firstly, where phenotypes and such stratification correlate, false positive associations can arise at SNPs whose frequencies vary between groups. Methods such as the use of mixed models^2,3^, and correction using principle component analysis (PCA)^4,5^ have proved powerful in solving major issues. However subtle population structure, such as within Europe, has been shown to remain a potential cause of bias in effect size estimates^6,7^, and this is a challenging issue in studying, for example, the evolution of traits through time^6^ or between human groups. Secondly, PGS typically have considerably reduced accuracy in other human populations in groups other than those sampled by the respective GWAS^8,9^, particularly those with stronger genetic differentiation to these groups. One explanation for this lack of portability is “local” around causal variants: genetic drift can leave some causal variants group-specific, while different linkage disequilibrium (LD) patterns between groups mean tagging variants may correlate with causal variants well in one group and much less well in another.

As well as such “local” causes, a variety of recent studies^10-14^ have suggested that causal variants have different average effect sizes on traits in different groups, due to “global” (i.e. non-local)” factors operating *outside* the region in LD with a causal marker. These Genome-wide or “global” changes in the effect sizes of causal SNPs, defined as their mean effects on a trait of interest, can arise by either gene-gene interactions not captured by the additive model, or gene-environment interactions, differing across populations. Such interactions have been documented to occur within populations^9^, so it seems likely *a priori* that they may contribute to differences between populations also. However, other authors have suggested similarities across groups in underlying effect sizes^15-19^, and studies show that at least the direction of effect of stronger GWAS hits appears conserved among groups^20^.

One recent study^14^ leverages individuals of mixed African and European ancestry, decomposing local ancestry to test whether within such individuals, causal variants occurring on African vs. European chromosomes show different effect sizes. By focussing on within-individual effects in African-ancestry populations, this approach largely removes the effects of factors that might result in different effect sizes across populations of different continental ancestries, including gene-environment interactions^14,21^ as well as long-range gene-gene interactions^21^. It also tests whether, for example, local gene-gene interactions might sometimes operate to alter effect sizes on different ancestral backgrounds. They infer strong sharing of underlying effect sizes at this within-individual level, suggesting such local interactions are *not* strong factors for most human traits, but also infer (using a different approach) strong differences in underlying causal effects across individuals possessing different continental ancestries. In Hou et al^14^, it is suggested that these might be explained by gene-environment interactions operating across individuals and groups. Resolving the existing controversy and uncertainty of the role of such interactions is vital in order to develop approaches to successfully apply genetic findings from one group in another – and even understand if this is generally possible for different traits – as well as for efficient study design, and for understanding whether evolution, or environmental differences, drives differences in trait observations between human populations. In this work we develop an approach to do this, using models consistent with the findings of Hou et al^14^. Specifically, we consider (in simulations and inference) models where genetic effect sizes are shared across ancestries within individuals, but can vary between individuals, for example those possessing mainly European ancestry and mainly African ancestry.

To provide a comprehensive analysis of the impacts of population structure we introduce a fine-scale ancestry pipeline. For convenience, we use the term “ancestry” throughout to refer to sharing an inferred most recent ancestor with a sampled individual from a particular self-reported ethnicity or geographic region. Building on our previous work^22-24^ decomposing the UK into 13 geographically meaningful populations, we design a new pipeline that achieves 23 regions including Ireland, and identifies 127 regions worldwide. These regions are geographically meaningful and genetically recoverable (see Methods), allowing us to describe ancestry information for all ∼500*k* (still participating) diverse UK Biobank^25^ (UKB) individuals. We show that using PCA for GWAS is unlikely to be the explanation for lack of portability, but detailed ancestry information provides improved correction for population stratification versus other state-of-the-art methods^2-4,26^: reduced false positives, and novel associations, across traits. We also develop a new statistical inference approach, ANCHOR, for settings where individuals are admixed, which complements approaches such as POPCORN^10^ and XPASS^27^ which can (only) be applied to non-admixed individuals (see Discussion). Importantly, ANCHOR does not require prior knowledge of the underlying disease etiology, for example the distribution of underlying effect sizes. As in Hou et al^14^ and other recent studies^11-13^, ANCHOR leverages decomposition of ancestry at fine scales *along* the genome, and assignment of mutations to particular population backgrounds. However, building upon recent results (e.g. Hou et al^14^), we aim to quantify the contributions of interactions involving genes and the environment on observed differences in predictive power for individuals drawn from different populations. Application of ANCHOR to 8,003 UKB individuals of mixed ancestry results in very different findings to other recent studies, and we discuss possible reasons and implications of our findings.

## Results

### A statistical pipeline to infer precise individual ancestry

We developed an approach able to break down the fine-scale ancestry of a single genome into a mixture of 127 regions genome-wide, including 23 regions within the UK and Republic of Ireland (Table S1). The UK Biobank dataset contains a majority of individuals of mainly British ancestry, but with substantial numbers of people from other worldwide populations: application of our approach identified 105 regions inferred as present in at least 5 individuals for at least 10% of their ancestry.

Briefly, our pipeline leverages data from previous studies^23,28-31^of human genetic variation, typed at distinct sets of variants, and methods^25,32^ to impute such inputs on a common set of variants designed to provide high-quality imputation across all data sources from a fully sequenced imputation panel, generating an initial unified “reference panel” of haplotypes (Methods; Figure 1a). We jointly phase this panel using SHAPEIT2^32^ to avoid differences in data quality by source. Reference haplotypes are given population labels via semi-supervised clustering combining ChromoPainter^33^ and fineSTRUCTURE^33^ clusters with geographic labels, to produce a painting reference panel.

**Figure 1.**
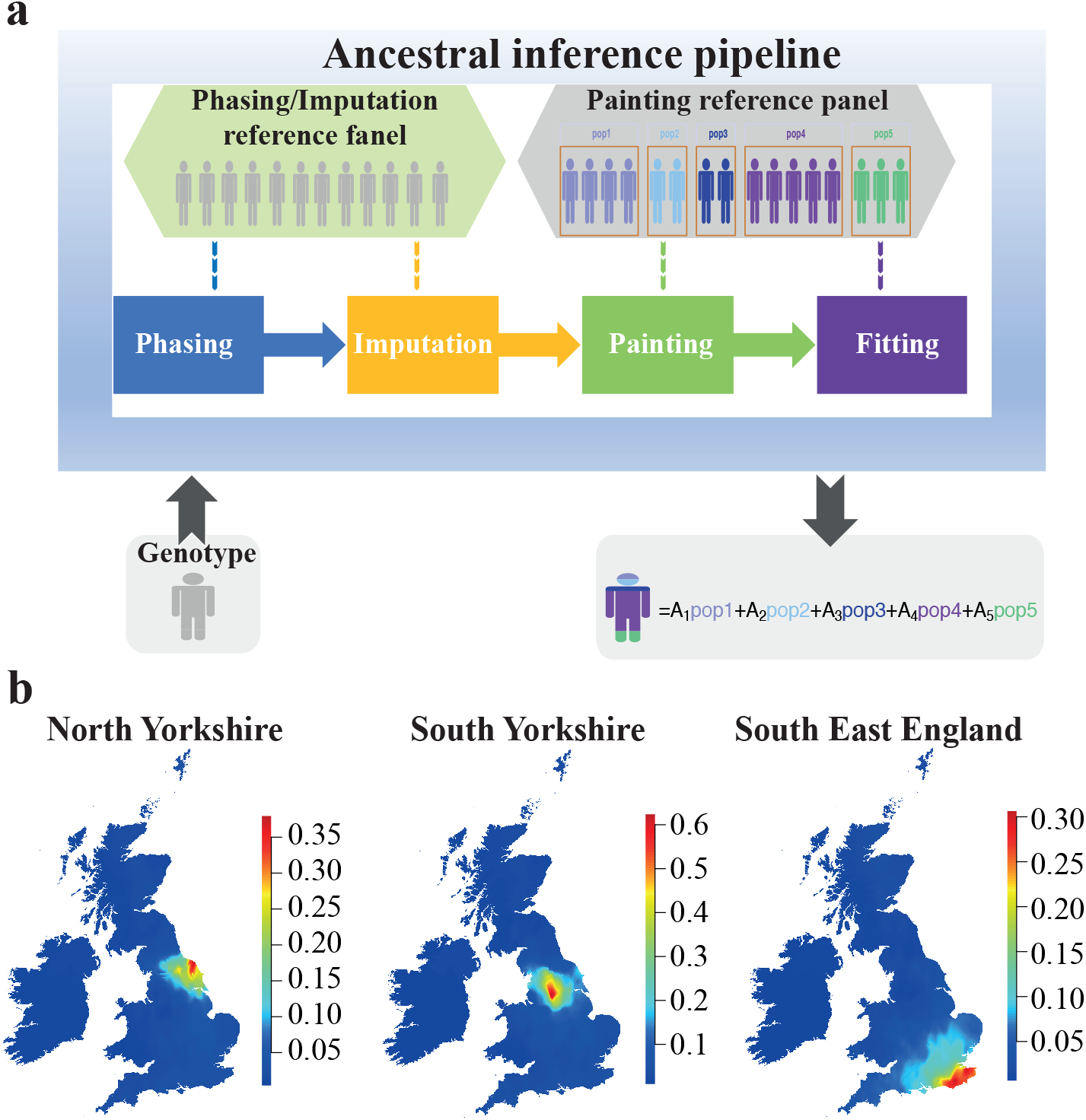
a)The main steps in our ancestral inference pipeline. The pipeline accepts individual genotype data (microarray or sequencing data) as input. The genotype data is phased and imputed against a phasing/imputation reference panel in first “Phasing” and “Imputation” stages, and painted against the painting reference panel including pre-clustered populations (5 populations in this example; 127 in our actual analysis). In a final mixture fitting stage, non-negative coefficients summing to one and representing the proportions of ancestries from the labelled populations in the reference panel are inferred. b) Average proportion of DNA in British-Irish individuals, placed on the map according to their birthplaces (see Methods), inferred to match to three UK groups: North Yorkshire, South Yorkshire and South East England.

Our pipeline leverages this panel to generate an ancestry decomposition for a new “target” sample (Figure 1a), extending our previously published approaches^33,34^, by closely mirroring the steps used to generate the reference panel itself. Unified panel markers are imputed in the target, using the imputation panel; if necessary the sample is phased (for UKB, haplotype data is pre-phased). Imputed haplotypes are then ChromoPainter-painted against the labelled painting reference panel, quantifying their level of ancestry sharing with each reference group by an overall painting vector. Finally, we run a non-negative least square (NNLS) based method (Methods^34^) to infer a set of ancestry coefficients, using the ChromoPainter output: loosely, this fits the observed painting vector as a mixture of the average painting vectors expected for members of each labelled reference panel. This approach benefits from utilising haplotype information to infer population structure^33^ alongside HMM approximations to the coalescent^33^, allowing us to analyse each sample in minutes (Methods).

We applied our pipeline to all 487,409 UKB participants. Because our pipeline uses only “out-of-sample” comparisons based on population structure, analysing single individuals independently, we note that it is designed to capture information about population structure, including within the UK and Ireland, and does not capture genetic relatedness between samples. Ancestry is defined in terms of partially subjective reference populations (in our case, those identifiable via fineSTRUCTURE), and the true *genetic* ancestry of UKB individuals is not known. Therefore, we assessed the usefulness of the pipeline in terms of its ability to effectively capture variation in ancestry across individuals in several ways: first, the relationship between inferred ancestry and individuals’ birth-place and self-reported ethnicity, which is also of direct interest; second, the ability of our ancestry components (ACs) to predict PCs, a well-established tool to identify genetic population structure, and thirdly, and importantly, the ability of our ACs to correct for ancestry in GWAS, compared to PCs and also Bolt-LMM, two widely used approaches^4,35^.

Alongside estimating ancestry differences among individuals, we also developed and implemented an EM-algorithm (Methods, Supplementary note 1) providing estimates of allele frequencies across SNPs, in the form of an estimate of the underlying frequency of each of 25,485,700 mutations imputed into the UKB^25^ in each of the 127 reference populations. These estimates are available from UKBiobank (https://www.ukbiobank.ac.uk) and also from the link shown in Data Availability and may aid researchers studying, for example, differences in regional allele frequencies within the UK.

### Fine-scale population structure across the UK

As a first test, the mean British-Irish (BI) ancestry inferred using our pipeline for the 434,781 UKB participants born in the UK/Republic of Ireland with self-reported white British/Irish (in short WBI) ethnicity was very high: 94.9%, with remaining contributions from Dutch (1.35%), Swiss (0.79%), Norwegian (0.49%), Polish (0.29%) and Danish (0.19%) inferred sources. The BI proportion falls dramatically, to 25.5%, for participants born elsewhere with a self-reported ethnicity of “other white background”. We note that even individuals born outside the UK/Ireland may possess some BI ancestry. At a finer scale, for individuals born in the Republic of Ireland, mean inferred Irish ancestry is 74.2%, while total inferred BI ancestry is 98.4%. Thus at this level, our pipeline is able to capture geographic and ethnicity information well (Figure 1b, Figure S1). Next, we plot the average proportion of ancestry inferred to be from each of 22 regions within the British Isles (see Methods for region definitions), by birthplace of UKB individuals (Figure 2a). These ancestry proportions are defined by an out-of-sample dataset of UK individuals whose grandparents were all born within a ∼80km radius and within each particular region^23^. For all BI regions, mean ancestry associated with that region decreases with geographic distance from that region, indicating DNA information is informative for birthplace (Figure 1b, Figure S1). 41.5% of UK-born individuals show the majority (>50%) of their ancestry from a single region, matching their birthplace 59.2% of the time, and this or a neighbouring region 82.7% of the time(Table S2). We also saw a strong correspondence between self-identified ethnicity and birthplace for non-UK ancestries (Figure 2b, Figure S1). However, within this overall pattern there is considerable regional variation: ancestry localisation is weaker in the south and east of England^23^, and stronger in Scotland, Wales, Northern England and South-west England. An entropy-based statistic (Methods) quantifies regional variation in mixing, in terms of smaller contributions of multiple ancestries to many people. The highest entropy in the UK is for individuals born in London, alongside the highest average fraction of ancestry from elsewhere in the world, while other major cities and South-east England in general also show strong mixing (Figure S2). Inferred non-British ancestries across the UK also vary geographically. For example higher inferred Irish ancestry (>10%) is seen in several large cities: Liverpool, Birmingham, Manchester and London, while Dutch ancestry is mainly found in the South of England, and around Bristol (3.5%); Polish ancestry has a peak near Wrexham in Wales, the site of the second highest concentration of Polish burials in the UK^36^.

**Figure 2.**
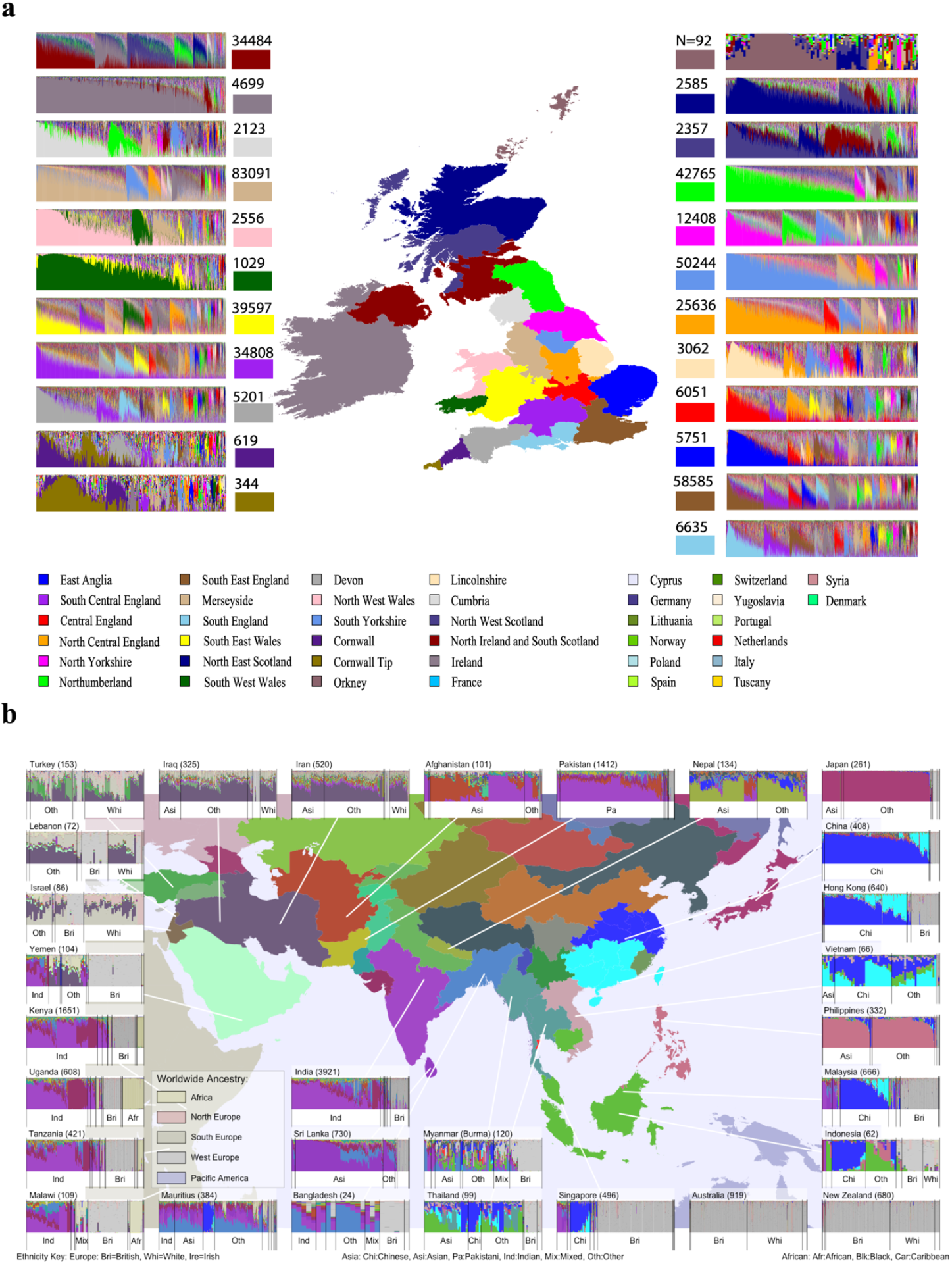
a) Ancestry inference stratified by birthplace region for UKB WBI individuals; for each regionally labelled barplot, each column shows ancestry decomposition for a single individual, with colours representing regions shown on the map; numbers represent counts of individuals from each area. b) As a, but showing decomposition for Asian, Oceanian and selected east African countries. Colours are as shown on the map, with colours for ancestry from additional regions given in the legend.

For individuals born outside the UK (Figure 2b, Figure S3, Table S2) we are likewise able to obtain fine-scale ancestry resolution. A minority of countries (e.g. Philippines, Japan) are inferred to be relatively genetically homogeneous, with most inferred as complex mixtures. Because UKB individuals reflect people who made their home in the UK, rather than unbiased samples from birthplace countries of non-UK-born individuals, we observe widespread BI ancestry in such individuals, and other events, such as the post-colonial exodus of South-Asians from many East-African countries, likely explain features such as the over-representation of Bangladesh ancestry for UKB individuals born in Uganda and Kenya.

### Ancestry components improve stratification control in GWAS for traits with geographic correlation

It is essential to correct for population stratification, to avoid false-positive GWAS associations. Principle components (PCs) are widely used for this purpose^4,5,37^, either alone or alongside other approaches such as mixed-model analysis^3,35^. One complication can be choice of number of PCs to include; using too few can result in false positive associations, while later PCs can be strongly impacted by extensive regions of LD within the human genome (e.g. beyond the first 40 publicly released UKB PCs^38^), resulting in potential overcorrection in GWAS. We explored the potential for utilising the 127 natural “ancestry components” (ACs) corresponding to the proportions of ancestry from each of our fitted groups. and contrast this with PC-based approaches in GWAS. As an initial test, we predicted the first 16 UKB PCs from these 127 ACs, and conversely predict 16 common BI ACs from the first 140 UKB PCs (Methods; Figure 3a-b, Figure S4, Figure S5). This reveals a good prediction of most PCs from ACs, but not the converse, suggesting that indeed ACs might capture additional information (Figure S4). PC-based prediction was also poor for non-UK regions (Figure S5).

**Figure 3.**
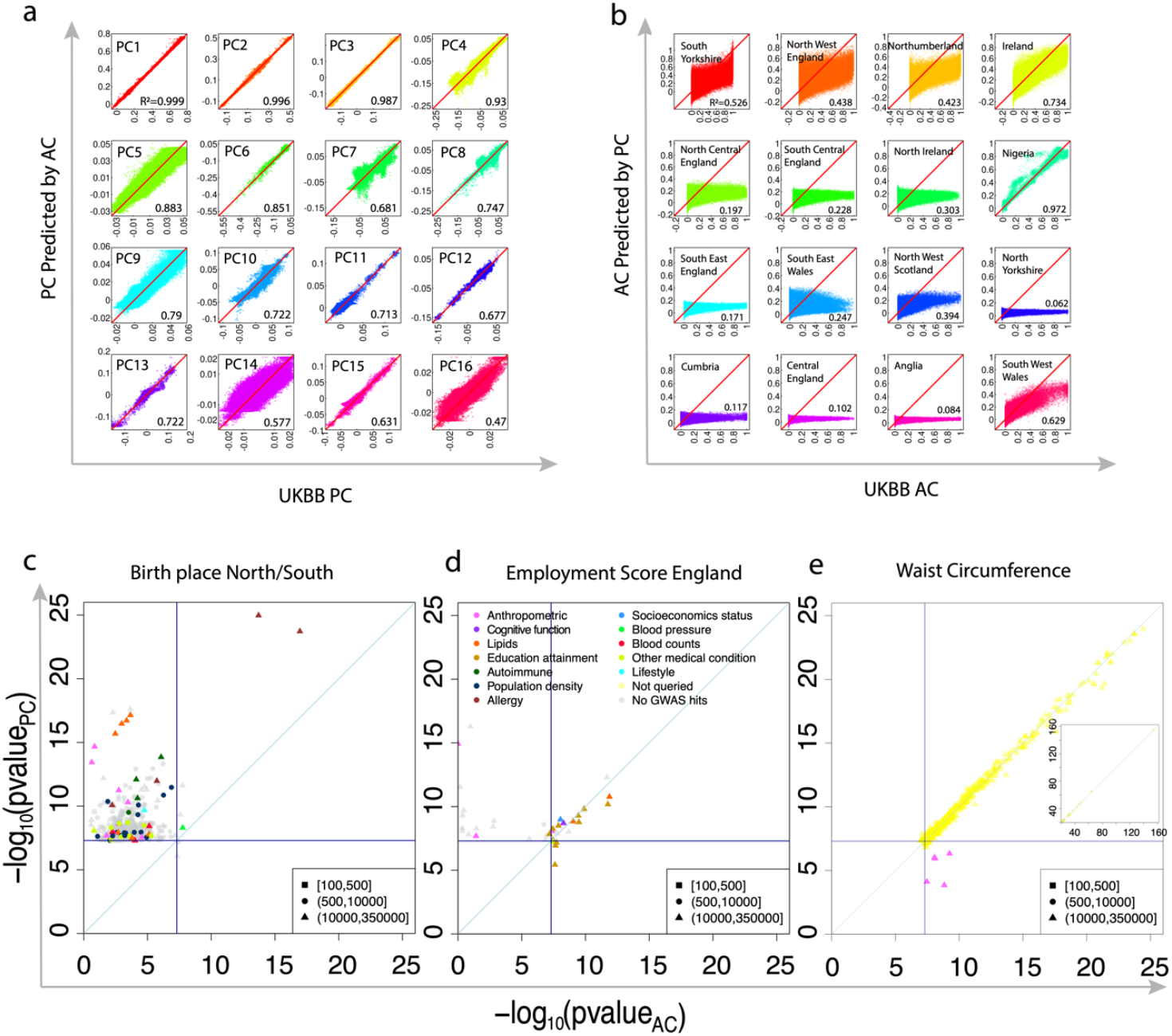
Comparison between AC-corrected and PC-corrected GWAS. a) Prediction of first 16 UKB PCs (x-axes) using a linear model-based prediction from the 127 ACs (y-axes) shows strong correlations (numbers). As a), but now predicting 16 UK and worldwide ACs from 140 PCs, often showing poor prediction. c)-e) Comparison of AC-corrected (x-axis) and PC-corrected (y-axis) -log10(p-values) in three exemplar GWAS for labelled traits; all plots are coloured according to the top legend in panel d, indicating prior evidence from GWAS for each SNP in particular phenotypic categories (grey SNPs show no prior evidence). In each plot, the points show only independent SNPs showing p<5×10^−8^ for one or both approaches; lower legend indicates the minor allele counts for the variants.

The most direct test of AC utility is correcting for stratification within GWAS, compared to PCs. We ran GWAS on each of 99 UKB quantitative traits with sample size of unrelated white British individuals larger than 10,000 (Methods). To quantify evidence of potential systematic inflation due to uncorrected stratification by either approach, we used LD-score regression (LDSC)^39^. LDSC provides an intercept estimate, generally expected to be 1 in the absence of confounding, although for strongly heritable traits such as height, or large sample sizes, intercepts slightly above 1 are often seen in practice^39,40^.

Birth location is often used as a control phenotype to test for stratification^41^ as there are few causal mechanisms for some SNPs to become more associated with it than the average. For latitude (Figure 3c), PC-based correction or analysis by Bolt-LMM (Figure 3c, Figure S6) yielded many apparent independent hits, and substantial genome-wide inflation (LDSC intercept of 1.6608), which remained even when 100 PCs were used (Figure S6). Encouragingly, almost all genome-wide significant signals are removed by the use of AC’s, implying uncorrected impacts of stratification from standard GWAS approaches, removed by the use of fine-scale ancestry information. Interestingly, easily the strongest remaining signal (p<10^−15^) is for a region of chromosome 4 around rs5743618, containing several toll-like-receptor genes including TLR1 and TLR10 which play a role in innate immunity^42,43^. This is one of the strongest GWAS hits in the genome for hay-fever in European populations^44,45^, with weaker effects on asthma risk. Because the protective allele against hayfever is more common in south England where hayfever is most prevalent (Figure S7), the geographic differentiation^46^ of this variant might reflect past natural selection. The attendant signal suggests that this selection (of migration or varying reproductive success depending on allele carried) has continued until recent times.

As another test of the use of AC’s and potential impacts of stratification in humans, we performed a GWAS for “employment score”, defined for England only. This is a trait not defined at an individual basis, but based on the region in which a person lives. Previously, GWAS associations with such traits have been observed^6^. However their use generates particular concerns due to the potential for regional stratification to impact regionally defined phenotypes, resulting in confounding. Indeed, for this trait results from ACs and PCs differ, with a number of shared hits, but some regions showing significant signals only for the PC-based analysis. Moreover, the AC associations were mainly significant in individual-based GWAS for cognitive traits and particularly education attainment (Figure 3d, Table S3), while the PC-only hits do not show such evidence, suggesting these are likely false positives (Figure 3d). These results alongside the poor PC-based prediction of ACs provide strong evidence that previous GWAS for “regionally defined” traits can be susceptible to false positives, and ACs correct for this at least in part.

Among the other 97 traits, PC-based and AC-based correction provided highly similar p-values and LDSC intercepts with some subtle improvements for AC (Figure S8). However, we still sometimes observed differences: 5 hits seen *only* in the AC-corrected analysis (Method, Figure 3e, Table S3), including some in strong LD with SNPs previously reported as GWAS hits for the same traits^45,47-49^, and thus unlikely to be false positives. In cases we examined, such as waist circumference (Figure 3e, Table S3), AC-specific hits occurred at SNPs falling within a region of strong LD with high regional SNP loadings for particular PCs (Table S4; Figure S9). Therefore applying ACs likely removed false negatives caused by PC overcorrection from PCs strongly correlated with genotypes in particular genomic regions.

### Biological effect sizes are highly similar across ancestries for a range of human traits

To study the extent and causes of (non-)portability of PGS across groups within UKB, we created PGS using 340K white British individuals for 29 heritable (estimated heritability >10%) UKB-measured traits, correcting for ancestry using either ACs or PCs, and tested their performance in independent samples representing different ancestries: with majority (>50% by ACs) inferred ancestry from seven different labelled regions: South Central England, Northumberland, Republic of Ireland, Poland, India, China and West Africa. Correction by ACs vs. PCs yielded different mean PGS for each group, but strong correlations (92.7-99.3%, Figure S10, Table S5): we focus on AC-corrected PGS for remaining analyses. By regressing the PGS against actual trait observations in each group, we quantified the increase β in trait per unit increase in the PGS (i.e. the regression slope), and the variance of the trait explained by the PGS after removal of variation due to other regressors including age, sex and AC’s; Methods), ΔR^2^. As expected, we observe a strong decline in both ΔR^2^ and β with increasing genetic distance from the discovery population of British-ancestry individuals (Figure. S11 and S12) for other continental ancestries, particularly those individuals with sub-Saharan African ancestry, with a >2.2-fold reduction in ΔR^2^ implying the PGS loses almost all predictive power. Our fine-scale ancestry information reveals this occurs even between BI regional ancestries for some traits: for standing height, FEV1 and Apolipoprotein B, ΔR^2^ for Northumberland differs from that for Ireland (p<0.02, p<0.005 and p<0.02 respectively).

To partition the drop-off in performance of PGS in different ancestries into local effects (such as causal variant tagging) and non-local effects (including gene-gene or gene-environment interactions), we developed the approach ANCHOR. As in previous approaches to either improve PGS performance^50-52^, examine effect sizes at individual markers^12^ or to compare effect sizes between chromosomes *within* individuals^14^, ANCHOR leverages variation in local ancestry along the genome and between admixed individuals (Figure 4a). Briefly, ANCHOR takes as input PGS coefficients from an independent sample, and quantitative phenotype and genotype data for a group of admixed individuals. Here we use estimated PGS coefficients from 340K UKB white British samples, and analyse 8003 “African Ancestry” individuals with varying (mean 83.6%) inferred sub-Saharan African ancestry and BI+Europe (mean 11.5%) ancestry. First, local ancestry is inferred along the phased genome, e.g. using HAPMIX^53^ here. Second, separate PGS’s are calculated for the African-like and European-like segments, yielding “APGS” and “EPGS” scores respectively for each individual, which must be centered by the ancestry-dependent mean SNP frequencies (Supplementary Note 2). Third, we regress the phenotype against APGS and EPGS, including other covariates, to estimate coefficients β_Af_ and β_Eu_, which quantifies the predictive power of the PGS on each background. Finally, we compare β_Eu_ estimated from admixed individuals to a β_Obs.Eu_ estimate obtained through analysing genetically similar non-admixed individuals (here BI individuals).

**Figure 4.**
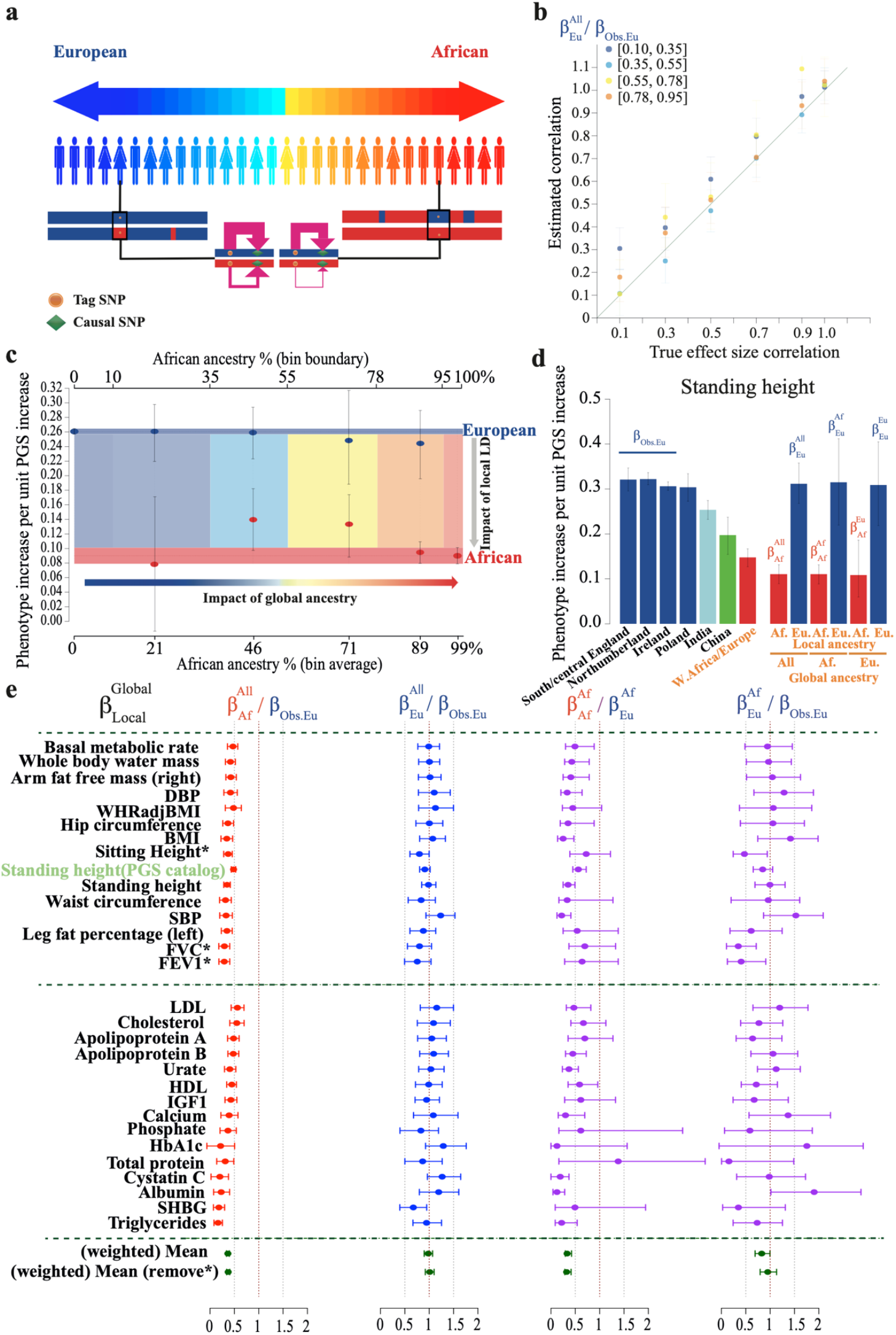
Separation of local and non-local factors influencing polygenic score portability. a) Test principles: in UKB samples containing segments of European (blue) and African (red) ancestries, a single causal variant contributing to a trait is captured by a tag SNP within the PGS. The predictive power of this tag SNP (pink arrow thickness) can vary according to local factors varying with “local” ancestry (top vs bottom chromosomes), or non-local factors impacting the effect of the causal SNP itself, quantified by Genome-wide “global” ancestry (left vs. right individuals); ANCHOR separates the contribution of these to PGS portability. b) Performance of ANCHOR in simulations of 24 traits (Methods), with varying true correlation between underlying effect sizes in African-ancestry individuals (x-axis) and European-ancestry individuals shows accurate estimation (y-axis) across five distinct levels of African ancestry (colours denote bins defined as in c) c) Application of ANCHOR for 29 UKB traits, for individuals of varying African ancestry binned as shown (x-axis; coloured regions). For each bin, mean estimates of coefficients β_Eu_ (blue) and β_Af_ (red) across traits are shown with 95% bootstrapped confidence intervals, representing the increase in phenotype per unit increase in PGS within European and African genomic regions respectively. Also shown are these estimates from individuals of ∼100% European (β_Obs.Eu_; blue horizontal bar) or ∼100% African (red horizontal bar) ancestry. The case ρ=1 where true effect sizes are identical across individuals of varying ancestry corresponds to blue points lying along the blue line, and similarly for red points, as observed. d) Mean increase in standing height per unit increase in its PGS (columns, with 95% bootstrapped CIs) for 7 ancestral groups, alongside local-ancestry-specific effect size estimates of from ANCHOR model fitting (final 6 columns; Methods), for global and local ancestry combinations as labelled on the x-axis and above the bars: 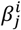 refers to local ancestry *j* and global ancestry *i*. e) ANCHOR results by trait. Columns show ratios of estimates defined as in d), rows show phenotypes: the second row for standing height uses a previously published PGS^16^, the final two rows show combined estimates. The second and fourth columns estimate the underlying correlation in causal effect sizes between European-ancestry and either all 8003 African-ancestry individuals, or individuals of 100% African-ancestry, respectively, while the first and third columns show the impact of local effects on predictive power for African-ancestry segments in the same two settings.

Consider a true (unknown) underlying PGS operating in European-ancestry UKB individuals, such that each unit increase in the PGS yields a unit increase in the trait in European ancestry segments. We define ρ as the average trait increase observed instead in European segments of admixed individuals. Under the “null” model with only local effects, the underlying causal effect sizes are identical in all European-ancestry regions, and ρ=1. If non-local (gene-gene or gene-environment) interaction effects do occur, then ρ<1 (Supplementary Note 2). Under reasonable modelling assumptions, the ratio β_Eu_/β_Obs.Eu_ estimates ρ, which is also equal to the correlation in effect sizes between the two groups if the variance of underlying effect sizes is the same in each (Supplementary Note 2). Assuming effect sizes scale linearly with genome-wide ancestry, ANCHOR can predict ρ in individuals of 100% European or African ancestry without sampling them (Supplementary Note 2; Methods). Because ρ is dimensionless, we can also combine estimated ρ values among groups of traits; this pooling helps reduce overall uncertainty. We obtain confidence intervals for all coefficients by bootstrapping individuals (Methods).

We verified that HAPMIX^53^ is able to accurately infer ancestry and allow ancestry-specific PGS construction via ancestry-aware phasing, by comparing to a group of individuals within trios where phase is mainly directly inferred (Figure S13, S14). We next tested the full ANCHOR procedure by simulating 24 quantitative traits with a range of heritabilities (0.3-0.6), numbers of underlying causal markers (100-10,000), clustering of causal mutations, and causal marker frequency spectra, using genetic data from the 8003 African-ancestry individuals (Methods). We analysed these, including performing GWAS, exactly as the real phenotypes. We simulated both under a null model of identical effect sizes for all individuals in UKB (i.e. no non-local interactions altering these effect sizes), and also under alternative models with varying underlying effect size correlations ρ. Firstly, to test the model we applied ANCHOR separately to individuals with similar genome-wide ancestry. Averaging over traits shows that if ρ=1, on average β_Eu_=β_Obs.Eu_ (Figure S15) across ancestry bins, and for 96% of individual traits, despite increased uncertainty the resulting 95% CIs contained the value ρ=1 (Figure S16). When we simulate non-local influences via different underlying causal effect sizes, so that ρ declines, β_Eu_ estimates decline correspondingly and ANCHOR estimates of ρ remain well calibrated (Figure 4b; Figure S15). In all cases β_Af_ < β_Eu_ as expected, due to local effects, but β_Af_ values otherwise rise and fall similarly to β_Eu_. These results across a wide range of settings provide confidence in the approach.

We applied ANCHOR to the actual UKB data of 29 quantitative phenotypes, again first binning by overall African ancestry to test robustness of our modelling assumptions, and average results across phenotypes. Examining (Figure 4c) the resulting β_Eu_ and β_Af_ estimates to the corresponding β_Obs.Eu_ values for individuals of 100% European (or β_Obs.Af_ for near-100% African ancestry) revealed strikingly constant β_Eu_ values across genome-wide ancestry bins. These β_Eu_ estimates overlap the corresponding estimate of β_Obs.Eu_ (narrow blue strip), which we obtained from a large number of 100% European individuals, and in no bin does ANCHOR reject the null ρ=1. Thus underlying effect sizes appear to be highly conserved between the African-ancestry and European-ancestry groups, despite the poor performance of the PGS in the former, across this broad range of human traits, including both molecular and quantitative phenotypes. Within the genomes of African-ancestry people in UKB, the European-ancestry segments retain predictive power as strong as that for European-ancestry individuals. Similarly, β_Af_ estimates vary little and match the corresponding β value estimated from individuals with near-100% African ancestry (Figure 4c), suggesting conserved effect sizes across the entire ancestry spectrum.

For individual traits, as in the simulations we combine all individuals and, as well as estimating an overall ρ, performed extrapolation to estimate implied effect sizes for European segments in individuals with almost 100% European ancestry genome-wide, or almost 100% African ancestry genome-wide (Figure 4d-e). For an exemplar trait, standing height (Figure 4d), we show the resulting β_Af_ and β_Eu_ estimates alongside β_Obs.Eu_, and β estimates for a variety of other AC-defined UKB populations, including African-ancestry individuals treated as a single group. β estimates decline with increasing genetic distance as underlying LD changes cause the PGS to have decreasing predictive power. However, the ANCHOR PGS decomposition by local ancestry identifies that the European-ancestry segments within the genomes of African-ancestry individuals have strong predictive power, within the range of β values for wholly European-ancestry individuals (blue bars), and the extrapolation does not identify any significant changes in effect sizes across the range of genome-wide ancestry. African ancestry segments (red) show a much lower ability to predict the trait, due to local LD and allele frequency differences, implying local, not genome-wide, drivers of non-portability.

Across all 29 quantitative and molecular phenotypes (Figure 4e), we saw very similar results to height: strongly reduced overall PGS power compared to non-admixed European individuals (first column), ρ estimates (second column) indistinguishable from 1, even after extrapolation to the individuals with the highest African ancestry (fourth column), indicating embedded European segments with similar predictive power to those in non-admixed European individuals. Results were also similar using a previously published alternative PGS^16^ for Standing height (Figure 4e). However, for two lung-function traits: forced expiration volume and forced vital capacity, and the correlated trait^54,55^ of sitting height, we see ρ estimates nominally significantly below 1, suggesting a role for gene-environment or gene-gene interactions. Our results provide evidence that there are some SNPs with reduced effect sizes of the same mutation on these traits in these UK-based African-ancestry individuals, compared to European-ancestry individuals. In future, increased sample sizes should offer greater power to detect such effects for these and other traits.

## Discussion

Here, we introduce new approaches to understand human ancestry, and its connections with the analysis of human traits, both for GWAS and subsequent trait prediction using PGS. We show that decomposing individuals’ ancestry into ancestry components is possible at the level of individual counties. Utilising these ACs improves correction for confounding, by capturing information missing from PCs, both in the form of reducing false positive (associations associated with ancestry) and false negatives (associations missed due to correlation between PCs and LD structure). When considering latitude (or longitude) as a phenotype, present approaches insufficiently control for geographic effects, while ACs offer unprecedented power to do so, within UKB at least. Many traits performed similarly in terms of p-values using ACs vs. other popular approaches, but we identified issues in conducting GWAS for “regional traits” defined on groups of individuals^6^, where only the AC correction avoids false positive associations. Together with observations of false negatives in GWAS using PCs, and potential problems in future with very large sample sizes, we suggest incorporation of ACs into GWAS will likely improve correction for stratification. ACs reduce overcorrecting as they are not associated with local genomic regions, which avoids the issue of deciding on the number of PCs to incorporate into an analysis. The complex patterns of ancestral mixing we observe within the UK, even within regions, with differences between e.g. cities and rural areas, suggest particular care is needed for traits such as educational attainment^24^ which show similar regional variation. A limitation of our approach is that our ACs are somewhat UK-specific – we anticipate benefits from the development of region-specific ACs for non-UK GWAS, and before such tools are available ACs might be used alongside PCs. Of course, it is possible that even our best available ACs may not completely correct for subtle population stratification.

In comparing underlying (causal) effect sizes across human populations, we took the approach of considering, for a causal mutation, its mean effect on a trait^12,56^ – for example, a mutation might increase height by 5mm in WBI ancestry individuals in UKB – and then comparing this value across groups. Although in some settings it is natural to scale genotypes by their population standard deviation and therefore scale effect sizes also, this is not the case when comparing populations. Because each causal variant operates within individuals to perturb a trait, across groups their effect sizes are *a priori* unlikely to correlate entirely with population mutation frequencies. Using genotype-scaled effect sizes might therefore downwardly bias estimates of correlation^12^, and so we avoid scaling rather than directly rescaling ancestral specific effect size in terms of allele frequency in corresponding ancestry^14^. We instead define a mutation as having identical impacts in two groups if each copy of the mutation results in the same average increase in phenotype in each group. This seems natural from a biological standpoint, and also uses a natural, interpretable scale for practical downstream applications of PGS, bypassing, for example, issues concerning varying overall PGS variances.

Our approach ANCHOR leverages local ancestry inference to allow estimation of the ratio of underlying effect sizes (defined as above) in an admixed group, to a group similar to that used for the initial GWAS and PGS construction, by directly estimating the predictive power of PGS among groups in terms of the “mean increase in trait per unit increase of the PGS”. This ratio is normally only 1 if effect sizes are identical, and otherwise estimates the correlation between underlying effect sizes in the two groups (Supplementary Note 2). While a few traits such as forced expiration volume do show evidence of differences, overall our results indicate clearly that – within the UK at least – effect sizes for the true underlying causal variants are highly similar (average correlation 0.98±0.08) between individuals of African ancestry, and European ancestry, across a range of quantitative phenotypes. This implies that (for example) the causal mutation increasing standing height by 0.5mm in European ancestry individuals also, for the large majority of such mutations, does so in African ancestry individuals.

These results appear to differ from findings^56-58^of often considerably lower underlying correlations when comparing European and African-ancestry samples, using previously developed approaches based on modelling genetic variance components and estimating local inter-population LD differences. For example one study^56^ finds different effects for BMI in UKB, another^58^ finds different effects for height in a sample including UKB, and an estimated correlation of only ∼50% across 26 traits in UKB. Because ANCHOR makes few modelling assumptions, but simply considers prediction using the PGS, it is difficult to see how estimates so close to parity among people with widely differing ancestry (e.g. 100% European vs. >80% African) might result from confounding. It is likely that the methods are targeting different quantities. ANCHOR estimates the ratio of centered and unscaled effect sizes in African segments to that of European segments after correction for genome-wide ancestry. We have shown that this quantity should be well-behaved under the null, and that a) interaction between global ancestry and effect sizes, or b) effect size differences between groups that depend on local ancestry both lead to predicted ρ<1 (Supplementary Note 2).

LD is the most likely source of differences between groups, and potentially the way it is handled creates differences between studies. We showed that while African-ancestry segments show similar predictive power in all individuals, their power is hugely reduced by approximately 3-fold relative to European ancestry segments: approximately a 3-fold reduction in the increase in trait per unit increase in PGS, remarkably consistently across traits (Figure 4e). Our (ρ=1) simulations constructing PGS as for the real data show PGS coefficient estimates resembling the real data for European segments, but a more modest relative reduction (<2-fold for traits with >100 causal markers) for African-ancestry segments. Recent work^14^ examining correlations *within* admixed UKB individuals infers identical causal effect sizes for the African-ancestry and European-ancestry variants, indicating that the 3-fold drop we see is likely the result of reduced tagging of causal variants by the PGS in African ancestry regions, and providing evidence against e.g. local interactions among clustered causal variants altering effect sizes. If, for example, selection against trait-impacting variants^59,60^ results in larger frequency differences for true causal variants between populations than the randomly selected variants used in our simulations, this could explain the observed strong drop-off between groups. ANCHOR might be applied to groups outside the UK, for example African-Americans^14^, that show similar admixture patterns. Resolution for individual traits is expected to grow greatly as sample sizes in such populations improve in coming years.

An important feature – an advantage and potential pitfall - refers to how we treat genotypes in different groups. For accurate theoretical performance, ANCHOR requires genotypes to be *centered* in a specific manner and in practice. Without this procedure, we observed strong downward bias in correlation estimates (data not shown). This centering is an implicit assumption that ancestry produces a mean effect, which may arise through lifestyle, environment or genetic variation from SNPs not highlighted in the GWAS used to construct the PGS, and for which admixed individuals receive the average. The PGS is relative to this “ancestry PGS”, with mean zero for an individual, and is local-ancestry-dependent, differing from existing approaches^11,13^. It will therefore be vital to explore whether similar centering allows for improved PGS predictive performance, in other individuals of mixed ancestry. We do, in fact, observe differences in mean values of our fitted uncentered PGS among human groups, for both PC-based and AC-based GWAS correction (Table S6). However, these differ between the two approaches for most traits due to subtle differences in their effect size estimates, likely (from our other findings) reflecting incomplete correction for population structure. Therefore, the connection(s) (if any) between these values and those for the “true” underlying PGS seem complex and further work is required. A future approach leveraging local ancestry might allow such examination, of whether mean genetic scores differ among human groups.

Our results do not imply that gene-environment, or even gene-gene, interactions are not operating across the quantitative traits we investigated here, or other traits. Interactions still likely operate to cause variation in effect size across individuals with African-ancestry. Our results though imply these must largely be shared with other groups of different ancestries, so as to not produce differences in overall (mean) effect sizes across populations. Previous work^9^ has shown that effect sizes, even within UKB individuals of British ancestry, differ among cohorts stratified by age, gender, and socioeconomic status, for example. We also observe differences between males and females in mean effect sizes (Figure S17), and by leveraging the power of ACs to capture subtle genetic differences, we find evidence of different effect sizes – albeit fairly subtle – among groups of UK individuals stratified by ACs (Figure S12). We suggest that these may reflect gene-environment interactions, given the very weak genetic differences between these groups and because patterns vary among traits. However the very strong lack of portability, in the case of African-ancestry individuals where it is strongest, does *not* seem to be meaningfully driven by interactions, apart from specific traits such as FEV1. This result motivates future efforts at, for example, joint fine-mapping^61^ within UKB or its successors, using models incorporating the same effect sizes operating across groups, aiding power. It also simplifies the task of applying genetic findings obtained from one group to another, at least within a country such as the UK, because it implies improved identification and tagging of underlying causal variants is the main issue in doing so.

The UKB resource contains diverse ancestries, but has collected reasonably homogenous data for individuals within a single country, and our population comparisons occur within this environment. This avoids trait definition differences, and perhaps reduces environmental effects, and so helps to isolate underlying biological impacts. For example, our results likely exclude differential gene-gene interactions as a major driver of non-portability, even in other countries or settings, because these would still operate strongly within the UK. However, effect size differences might be stronger between different countries - if environmental differences are greater - and are highly likely to occur where trait definitions or diagnoses differ, perhaps particularly for disease traits. Thus, we expect that for admixed groups, future application of ANCHOR or related methods will identify variable effect sizes across countries or cohorts, while other approaches^10,57,59^ enable comparisons of groups of similar ancestry.

## Methods

### Ancestry pipeline

To construct an ancestry “painting” reference panel, we merged data from the table below, leading to 9129 samples from various genotyping platforms, whose union set of SNPs contains a total of 2,011,414 distinct mutations.

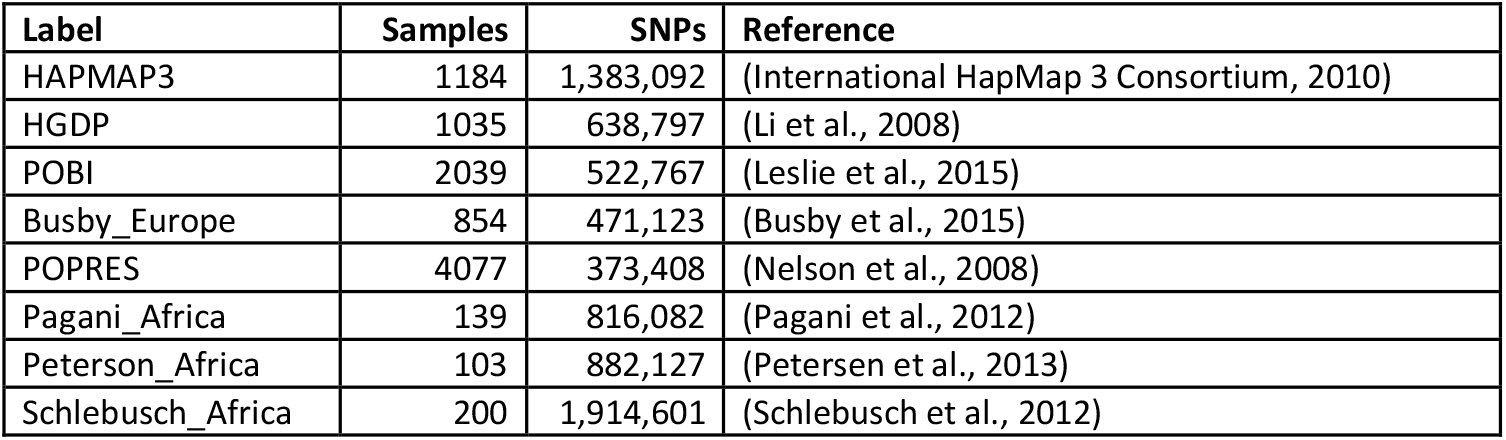

Following relatedness pruning with Plink we retained 7775 individuals. We then performed phasing (using SHAPEIT2^32^) and imputation (using IMPUTE4^25^) using the UK10K^62^ + 1000 Genomes data as a reference panel. We retained data for 851,948 sites that met a mean IMPUTE4 information score threshold of 0.9, across all the different genotyping platforms simultaneously, as well imputed sites, in the “draft painting panel”.

The remainder of the panel construction is described in Supplementary Note 1, but consists of the following stages. Firstly, we used a nested hierarchy of samples based on their representation in our painting panel samples, and so likely achievable resolution: Great Britain and Irish samples (BI), other (non-BI) European samples, African samples, and other Worldwide (World) samples. Next, we performed chromosome painting using chromopainterv2^33^ for each category of individual (i.e. 4 location groups), using only individuals within that category as donors (using an individual-level painting), for second round of quality control for relatedness based on genetic sharing. Thirdly, we formed updated “surrogate” (individuals used to construct average representative ancestry vectors, who must be unrelated to the donors) and “donor” (against who the painting is performed; because they are not directly compared, more relatedness can be tolerated) populations from sample labels and performed Non-Negative Least Squares (NNLS) admixture estimation following Hellenthal et al. 2014^22^, for each population, based on iterated reassignment. We used a semi-supervised approach to iteratively improve clusters based on their label compositions. Fourth, we performed additional quality control for the SNPs to include in the painting, based on their consistency with the chromosome painting. This resulted in 677,173 SNPs forming the “final painting panel”. Finally, we painted UK Biobank individuals who were born in regions of the world that we believed our reference individuals sampled from poorly, after excluding samples with e.g. mainly BI ancestry, and used their profile as a surrogate population reference vector (that is, these samples are not part of the painting panel per se, but their average painting properties are considered as a representation of ancestry from this region).

Annotating populations is an art, not a science and reasonable people may have made different decisions based on preferences towards “lumping or splitting” similar populations on a genetic continuum and a need to make ancestry components interpretable via a map. Our choices led to 206 surrogate populations that were genetically distinct, which we sum into the 127 interpretable ancestry labels reported in our results section. For replicability we include the SNP lists, and the final donor and surrogate population annotations (Supplementary Note 1)

### Painting Panel Processing

To obtain calibrated estimates of local ancestry, Individuals in the panel must be exchangeable with individuals we wish to compare to that panel. We therefore:

1. Construct a SNP list and sample list using the process above. Samples in the UK10K or 1000 Genomes dataset are removed.
2. Phase each sample independently, I.e *not* using the rest of the painting panel, against the UK10K + 1000Genomes dataset using SHAPEIT2.
3. Impute each sample independently, i.e *not* using the rest of the painting panel, against the UK10K + 1000Genomes dataset using IMPUTE4.
4. Using Chromopainterv2 in Expectation-Maximisation mode, we infer best-fit parameters *Ne* (which controls recombination rate) and *m* (which controls mutation rate) for 10% of samples randomly chosen for each Chromosome. We then average to obtain a single set of parameters. The painting is taken with respect to a “leave-one-out” panel, removing one individual at random from each other population, and the self from the current population, resulting in a set of populations with exactly one fewer sample than the total.
5. Using ChromoPainterv2, we paint with the inferred parameters, repeating the “leave-one-out” procedure above. Each sample receives a “donor-vector”, that is a total amount of genome shared (in centi-Morgans) with each donor population in total.
6. We obtain a surrogate panel by taking the average donor-vector within each surrogate-population.
7. Admixture estimates come from considering each sample’s donor-vector as a mixture of the surrogate-vectors in the surrogate panel using Non-Negative Least Squares (NNLS) as described in (Hellenthal et al. 2014)^22^. As part of QC, we had merged or removed any populations that were not >50% recovered this way.

### UK Biobank data included in the analysis

The UK Biobank is a prospective study of >500,000 people living in the UK. Both genotypic and phenotypic data used in this paper were analysed under application 27960 in the UK Biobank (https://www.ukbiobank.ac.uk). 487,409 UKB participants for whom autosomal haplotype and genotype imputation data (field IDs 22438, 22828) are available were used in our ancestry inference. Quality control, phasing and imputation for the UKB genetic data are previously described (http://biobank.ctsu.ox.ac.uk/). Demographic and phenotypic data we analyse here are listed in Table S7

### Running the ancestry pipeline on UK Biobank

In general, the ancestry pipeline accepts genotype data in various formats (different genotyping arrays) as input (see Supplementary Note 1 for details). For analysing the UKB data, we did not perform phasing as phased haplotypes are already available. To analyse the large-scale data of UKB, we performed the initial imputation step of the pipeline by running IMPUTE4 in batches of 1000 UKB individuals, before running individual jobs for the remaining steps:

1. Relative detection. Any close relatives are removed from the donor panel, along with 1 randomly chosen individual per donor population.
2. Paint. We paint the target using parameters Ne and m estimated from the panel to obtain a donor-vector for the target.
3. Admixture analysis. We infer global ancestry using NNLS.

### Assign UK counties with pre-defined UK regions

Within the UK and Ireland, the previous procedure identified 23 distinct populations. We assigned each UK county to one of these populations, to refine their geographic boundaries. Specifically we downloaded county-level UK map data (https://gadm.org) and mapped birthplaces of 426,879 UK-born UKB individuals to a county using the R package “sp”. After filtering to only those individuals with >50% ancestry from one of the 23 populations, we assigned a county to that population with the largest number of remaining individuals born within the county. “Irish” ancestry was assigned to the Republic of Ireland. Within Cornwall, we used the same approach, but because both “Cornwall” and “Cornwall Tip” localised to this county, we used the R package “raster” to define ancestry on finer-scale pixels, and assigned locations whose mean “Cornwall Tip” ancestry was at least 0.2 greater than their mean “Cornwall” ancestry to the “Cornwall Tip” population.

### Visualisation of ancestry based on UKB birth place

We used Gaussian kernel smoothing to generate spatial smoothed plots showing average ancestry. For each pixel in the rasterised UK map with birth place coordinates *P*=(x_p_,y_p_), we calculated the mean of a quantity of interest *O* that is defined for each individual, e.g. the ancestry coefficients estimated from our pipeline, across all the *N* = 426,879 UK-born or Irish-born UKB samples, smoothed by the Gaussian Kernel:

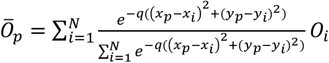

where *O*_*i*_ is the ancestral object and (x_i_,y_i_) is the birthplace coordinate for individual *i*. *Ō*_*p*_is the mean of *O* at pixel *P*=(x_p_,y_p_). We used adaptive bandwidth smoothing for *q* (Supplementary note 1).

To quantify ancestry mixing across the UK, we calculated for any individual their ancestry entropy

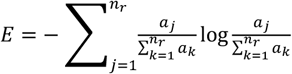

where *n*_*r*_ = 23 is the number of predefined British-Irish regions, and *a*_*j*(1≤*j*≤*nr*_) are the ancestry coefficients for that individual. *A*daptive bandwidth Gaussian kernel smoothing was used to plot the UK-wide entropy profile.

### Principal components and Genome wide association study

40 PCs (field ID 22009) were downloaded from the UK Biobank. The first 20 PCs were used in GWAS analysis. We additionally calculated the first 200 UKB PCs using the R packages “pcapred.largedata” (https://github.com/danjlawson/pcapred.largedata) and “*pcapred*” (https://github.com/danjlawson/pcapred), in which the PC loadings were constructed on near-identical SNPs to those used for calculating the UKB PCs, aside from a small number of unavailable SNPs. The correlations between the first 40 of these PCs and the 40 downloaded UKB PCs are all >99% (Supplementary Figure S18), so these PCs in practice extend the set relative to UKB.

127 ACs from the ancestry pipeline were used to regress each of 140 UKB PCs (chosen to exceed the number of ACs) on all the 487,409 UKB samples. The fitted linear model was then used to predict each PC. Similarly, we also used the 140 UKB PCs as covariates to predict the 127 ACs on the same UKB samples. *R*^*2*^ between the true PC and predicted PC was then used to evaluate the prediction. In our GWAS study, we analysed 99 continuous UKB phenotypes with sample size >10,000 in the analysis (Table S7). 343,047 unrelated white British individuals^25^ were included in our GWAS. Association testing was performed for UKB imputed SNPs with minor allele frequency larger than 0.001 and information scores larger than 0.3. Covariates included for each association test were “genotype measurement batch” (field ID 22000), “age at recruitment” (field ID 21022) and “sex” (field ID 31), as well as nonlinear terms “age^2^”, “age×sex”, “age^2^×sex”. The full model using ACs to correct stratification is:

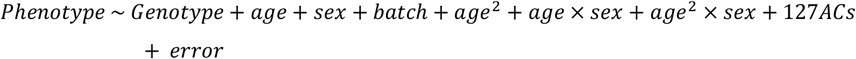

The full model using 20 PCs (and similarly for those tests using 100 PCs) to correct stratification is:

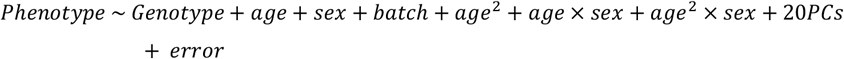

In both cases, association testing was performed by “BGENIE” version 1.2^25^. Unnormalised phenotypes were used, so that estimated effect sizes remain in units of the original phenotypes.

As well as using PCs to control for population stratification, we performed additional GWAS using Bolt-LMM^35^ with 20 PCs, to test performance of a linear mixed model for the place of birth, north coordinates (field ID 129) phenotype.

For Bolt-LMM, we QCed and pruned the UKB chip genotype SNPs using plink1.9^63^ www.coggenomics.org/plink/1.9/ (supplementary note 1) to generate 142,182 independent chip array SNPs (*r*^2^ <0.1) as input for null model building in Bolt-LMM. The participants and imputed SNPs for association testing remained as for the other GWAS. The 20 PCs included in Bolt-LMM as covariates were calculated by FlashPCA2^64^ from the same genetic relationship matrix (GRM) constructed by the 142,182 independent SNPs and 343,047 UKB white British individuals. Other covariates used in Bolt-LMM were the same as those used in the linear regression based GWAS.

To identify independent genome-wide significant signals, we used a threshold p<5×10^−8^ and for each SNP more significant than this threshold for either AC or PC, we used LD pruning with a threshold of *r*^2^ <0.01 and window size=100kb to keep only the most significant SNP among groups of variants in LD.

To identify previous GWAS hits, we queried these AC-significant or PC-significant SNPs in a GWAS database comprising over 10,000 curated, QC’d and harmonised complete GWAS summary datasets with API implemented in R package “mrcieu/ieugwasr” (https://mrcieu.github.io/ieugwasr/). For most traits we applied a genome-wide threshold of 5×10^−8^ for query output p value, though the threshold was lowered to 1×10^−6^ for traits where no SNP met this threshold. For each one of the three traits shown in Figure 3c-e, any queried results that contain the same or directly related traits were not used for downstream categorisation. Query results for input SNPs were ranked in ascending order with respect to their GWAS significant p-value across different traits such that the most significant trait was top-ranked (Table S3). For queried SNPs not possessing an “rsid”, which was used for query in “mrcieu/ieugwasr”, they were categorised as “Not queried”. SNPs with p-value of top ranked GWAS hit greater than 1×10^−6^ were categorised as “No GWAS hits”. The queried traits were assigned into the following categories: “Anthropometric”, ”Cognitive function”, “Lipids”, “Education attainment”, “Autoimmune”, “Population density”, “Allergy”, “Socioeconomics status”, “Blood pressure”, “Blood counts”, “Other medical condition” (Table S8).

There’s some exception for “Waist Circumference”. We identified in total 627 independent hits that were at least genome wide significant for one of AC or PC corrected approaches, and there were only 5 hits specifically significant in AC-corrected approach and with their value of 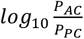 larger than 1, where *P*_*AC*_ and *P*_*PC*_ are *p*-values of AC-specific and PC-specific GWAS hits for “Waist Circumference”. We queried 4 out of those 5 AC-specific variants, which show major *p*-value difference between AC and PC (Figure 3e), using the query function in package “mrcieu/ieugwasr”. For the remaining AC-specific variant (15:84311431:TA:T), we couldn’t directly query it using the interface as it doesn’t have “rsid”, so we manually queried this SNP at opentarget^65^ (http://genetics.opentargets.org/). We found this variants strongly associated with many different “Anthropometric” related traits such as height, trunk fat mass and body fat percentage, so we labelled this variant as “Anthropometric”. For the remaining 622 variants, we didn’t query them and we categorised them as “Not queried” for the following reasons: 1). querying 622 variants in “mrcieu/ieugwasr” was computationally infeasible in the package “mrcieu/ieugwasr” and 2). most importantly, there’s no evidence of big difference between two approaches among those 622 variants as their absolute value: 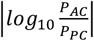, is less than 1.

As association between SNP and trait is more likely to be true if the SNP is also GWAS hit for other traits relevant to the trait of interest, we categorised the SNPs using the following criteria: for each of the three traits shown in Figure 3c-e, we assigned most relevant category to the SNP if the SNP is significantly associated with other traits which potentially “link” to the study trait within that category. For “Birth place (North/South)”, we assigned category “Allergy” to the queried SNP if it is associated with allergic phenotypes such as “hayfever”; for “Employment Score England”, we assigned category “Education attainment” to the queried SNP if it is associated with “Education attainment” or related phenotypes such as “Age completed full time education” or we assigned category “Cognitive function” to the queried SNP if it is GWAS hit for e.g. “Mood swing”; for “Waist circumference”, we assigned category “Anthropometric” to the queried SNP if this SNP is associated with e.g. “ Leg fat percentage”. If no proper category can be assigned to the SNP, we assigned category containing SNP’s top-ranked association trait to the queried SNP(Table S3).

### Heritability estimation and GWAS inflation assessment by LD Score regression

We ran LD-score regression^39^ on both AC-corrected and PC-corrected GWAS summary statistics for 99 UKB traits. We used pre-calculated LD scores using the 1000 genome phase 3 CEU panel provided alongside the LDSC software. From LD-score regression output of the intercept and *χ*^2^ values, we bounded these below at 1 to ensure non-negativity, and calculated a trait-wise LDSC attenuation ratio, defined as follows^40^:

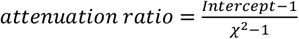

### Estimation of allele frequencies for UKB SNPs across UK/Ireland populations

Highly differentiated alleles across different regions reflect strong local genetic drift, which can for example identify evidence of natural selection. We derived an expectation-maximization (EM) algorithm to estimate maximum-likelihood regional allele frequencies based on our individual ACs (Supplementary note 1). We ran an EM algorithm on 12,977,776 imputed UKB SNPs with minimum minor allele frequency (MAF) at least 0.01% and imputation information score larger than 0.9 in the 339,304 white British samples with UK+Ireland ancestry larger than 0.95 (filtering, variant extraction and format conversion used ‘qctool2’, https://www.well.ox.ac.uk/∼gav/qctool_v2/index.html, “bgenix”^66^, and plink2.0^63^), to estimate their allele frequencies across the 23 British-Irish regions.

### Polygenic score calculations

For each of 29 traits, we constructed two polygenic scores that differed in whether they used ACs or PCs to correct for population stratification when estimating effect sizes in a GWAS of 343,047 unrelated white British individuals. PGS construction used the Hapmap3 SNPs^67^ of at least 1% minor allele frequency. For each phenotype, we performed LD-clumping by “plink1.9” ^63^ using the 1000G phase3 CEU as the reference panel with following parameters: significant threshold for index SNPs is 0.05, secondary significance threshold for clumped SNPs is 1, LD threshold for clumping is 0.1 and Physical distance threshold for clumping is 500 kb, with respect to the plink command line as follows: *“--clump-p1 0*.*05 --clump-p2 1 --clump-r1 0*.*1 --clump-kb 500*”. The PGS was then evaluated for testing purposes on the remaining 144,362 UKB individuals.

We then calculated the PGS as a linear sum over SNPs and genotypes:

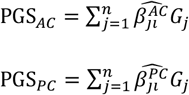

where for SNP *j, G*_*j*_ is the genotype for the target individual, 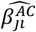 is the estimated effect size from the AC-corrected GWAS and 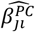 is the effect size from the PC-corrected GWAS. We note that mean-centering genotypes within each population would yield equivalent results (Supplementary Note 2), so we do not do this here.

For PGS_*AC*_, to evaluate performance across different ancestries we identified those individuals among the 144,362 UKB “test” individuals with at least 50% inferred ancestry from each of seven separate AC-labelled groups, by the ancestry pipeline: “South Central England”, “Northumberland”, “Republic of Ireland”, “Poland”, “India”, “China” and “West Africa”. By using this definition, we obtained 1,491 individuals in South Central England, 5,355 individuals in Northumberland, 13,265 individuals in Republic of Ireland, 1,503 Individuals in Poland, 4,438 individuals in India, 1,266 individuals in China and 7,108 individuals in West Africa.

For each group, and each particular phenotype, we evaluated PGS_*AC*_ for individual *i*, which we label PGS_*i*_, and then fit the model:

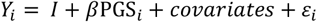

where *Y*_*i*_ is the phenotype for individual *i, ε*_*i*_ is the usual error, and the model was fit via standard linear regression, with 1000 bootstrapped sample replicates used to evaluate uncertainty. As previously the covariates included are batch, age, sex, age^2^, age×sex and age^2^×sex, as well as the 127 estimated ACs. The parameter *β* represents an estimator of the increase in the mean phenotype per unit increase in the PGS. We compare estimated values for this parameter among groups to quantify applicability of the PGS in distinct groups, and generalisations of this parameter are used also in our ANCHOR analysis.

Additionally, for each group we defined the “residual r^2^” ΔR^2^, a scale-independent measure of PGS performance, as 1 minus the ratio of the phenotypic variance remaining after fitting the above model to the (larger) variance fitting the same model with *β* = 0. This represents the fraction of variance explained by the PGS after accounting for confounding and the covariates.

We also estimated an overall value *β*_*Obs*.*Eu*_, for 19,596 individuals of BI ancestry by fitting the same model. The BI individuals were selected from three BI groups: South Central England, Northumberland and Republic of Ireland after filtering out individuals with the sum of these three ancestries less than 0.9.

### ANCHOR approach to constructing ancestry-aware polygenic scores and estimating correlation in effect sizes

To understand the behaviour of the PGS in African-ancestry individuals, we identified the 8,003 UKB European-African admixed individuals with the sum of 27 UK/European ancestries and 4 sub-Saharan African ancestries larger than 90% and the sum of 4 sub-Saharan ancestries larger than 10%. We further binned the overall 8003 individuals into 5 ancestry bins in term of their inferred African ancestry: [0.1-0.35],[0.35-0.55],[0.55,0.78],[0.78,0.95],[0.95,1]. The numbers of individuals and their mean African ancestry within each bin are: 373/0.207, 560/0.46, 475/0.707, 2556/0.89 and 4039/0.986. Individuals in bin [0.95,1] were considered as homogeneous African so we didn’t decompose the polygenic score into European and African for participants within that bin when running analysis from which the results are shown in Figure 4b-c and Figure S15.

In applying the ANCHOR approach, we generate and analyse ancestry-specific polygenic scores for a trait. We used HAPMIX^53^ to estimate local ancestry genome-wide for these individuals, using 1000G phase 3^68^ CEU and YRI groups respectively as European and African ancestry reference panels, run using default parameters. For individuals labelled *i* = 1,2, … *n*, the output of HAPMIX at each locus *j* = 1,2, … *J* provides probabilities 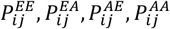 that the local ancestry is each of the four possibilities, ordering the two chromosomes arbitrarily (e.g. “EE” refers to both chromosomes possessing European ancestry). If the observed genotype at site *j* for individual *i* is *G*_*ij*_ then we first estimate the allele frequencies 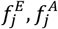 of this site on the European background and African background respectively by fitting the model

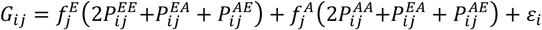

where *ε*_*i*_ is the (mean zero) noise, and the model is fit by least squares. In practice we fit the equivalent model, by noting 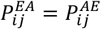 and because the ancestry probabilities sum to 1:

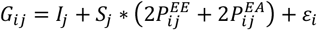

where after parameter fitting we obtain estimates 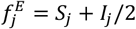 and 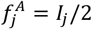.

HAPMIX also estimates expected allele counts 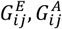 at this site on the European and African ancestry backgrounds respectively (obtained via summation; Supplementary Note 1), and we transform these to their mean-centered version (Supplementary Note 2), conditional on local ancestry probabilities:

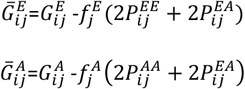

From these, the overall PGS may be decomposed into the European PGS (EPGS) and African PGS (APGS) for African-ancestry individuals *i* = 1,2, … *n* as follows:

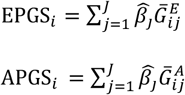

where 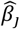 is the estimated per-copy effect size of SNP *j* on the phenotype. To investigate the role of local and non-local factors in attenuating the PGS performance in African-ancestry individuals, we fitted versions of the following model to real and simulated data (Supplementary Note 2):

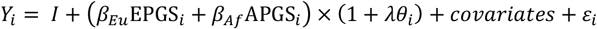

where for individuals *i* = 1,2, … *n, Y*_*i*_ is the phenotype, *ε*_*i*_ is the zero-mean noise, and the parameters to be estimated are the intercept *I*, and *β*_*Eu*_, *β*_*Af*_, λ. EPGS_*i*_ and APGS_*i*_ are the centralized EPGS and APGS as above, now after regressing out the covariates, and *θ*_*i*_ is the mean genome-wide European ancestry proportion for individual *i* = 1,2, … *n*. Covariates are identical to those described in the section “Polygenic score calculations”. We fit this model for various subsets of individuals, combinations of phenotypes, and parameter constraints (described below). In each case, we obtained parameter estimates via least squares (Supplementary Note 2), and uncertainty estimates by 1000 bootstrap resamples of individuals and model refitting. The model is linear so trivial to fit unless allowing *λ* ≠ 0; in this case, conditional on the ratio *β*_*Eu*_/(*β*_*Eu*_ + *β*_*Af*_) the model is again linear, so we use a grid search (1000 values) over this ratio from 0 to 1, fitting the linear model for each possible value and then minimizing the achieved sum of squares.

Parameters *β*_*Eu*_, *β*_*Af*_ measure the increase in the phenotype per unit increase in the appropriate polygenic score, so measure the predictive power of the score. Local factors mean that we expect *β*_*Eu*_ > *β*_*Af*_ (Supplementary Note 2; Figure S16 shows this via simulation). If there are no additional non-local factors contributing to non-portability, then *β*_*Eu*_, *β*_*Af*_ are expected to remain constant across ancestry bins, i.e. varying *θ*_*i*_, and also *β*_*Eu*_ will be shared between African-ancestry individuals and individuals of 100% European ancestry (Supplementary Note 2). If there are non-local factors, this no longer holds. To investigate this setting, it is natural to also allow that the predictive power of the two scores varies with genome-wide European/African ancestry i.e. *θ*_*i*_, and this is captured by *λ*. Because local factors still operate, the model covaries the predictive power of the *β*_*Eu*_, *β*_*Af*_ parameters together with *λ*; we found (data not shown) we in any case lack power with current sample sizes to fit separate effects, and so we simply use *λ* to capture linear effects (Supplementary Note 2).

We fit this model to analyze both real data, and simulated datasets in an identical manner. When jointly analysing phenotypes and averaging results, we first binned the African-ancestry individuals by their genome-wide ancestry *θ*_*i*_ (bin boundaries shown on Figure 4c). Within each bin ancestry varies little, so we set *λ* = 0 and fit the model independently for each bin for each phenotype. This provides estimates of *β*_*Eu*_, *β*_*Af*_ for each phenotype-bin combination, which we averaged within bins for Figure 4c, and this allows comparisons across bins, and with the effect size 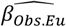 obtained for individuals of mainly European ancestry. Again holding *λ* = 0 fixed but now analyzing all individuals jointly for a phenotype produces estimates 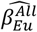 and 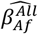 that summarize the predictive power of the PGS across the full African-ancestry sample set, shown in e.g. Figure 4d and 4e. The ratio of the means of the estimates over phenotypes, 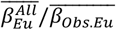, is an estimator of the correlation *ρ* in effect sizes between Africa-ancestry and European-ancestry groups, averaged over phenotypes (Supplementary Note 2); estimates of this are shown in e.g. Figure 4b,e for simulated and real data. For an individual phenotype, if the 95% bootstrapped confidence interval of the ratio 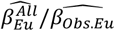 contains 1, we accept the null hypothesis of shared effect sizes. The ratio 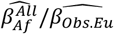 captures local (e.g. LD) effects on prediction, and so as expected is <1 for all simulated and real traits. Finally, we fit the full model allowing *λ* to vary. In individuals with 100% African [respectively European] ancestry, this fits the unit increase in the phenotype per unit increase in EPGS as 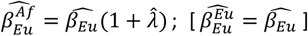, and analogous coefficients 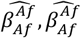 correspond to the APGS. These coefficients are shown in e.g. Figure 4d,e. Ratios of these quantities to 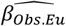, and their bootstrapped confidence intervals, are interpreted exactly as those for “All” individuals but now projected to estimate properties of individuals - for example ratios of true effect sizes – whose ancestry is 100% African or even 100% European. Where they appear in additional analyses, for example on different subsets of individuals, these quantities remain as defined here.

### Ancestry-aware simulation

To explore GWAS portability and evaluate performance of ANCHOR, we simulated phenotypes across the entire UKB cohort, and analysed as for the real data. There are 12,690,793 variants in the UKB imputed genotype data passing the following filters: minor allele frequency (MAF) ≥ 0.001, minor allele count (MAC) ≥ 25, genotype missingness ≤ 5%, HWE p-value ≥ 1 × 10^−10^ and imputation INFO score ≥ 0.8. From these, we chose a set of *J* causal variants (*J*=100, 1000 or 10,000) either at randomly selected genomic positions, or with clustering. For the clustered setting we selected *J/*10 non-overlapping 10KB regions in each of which, on average, 5 causal variants were placed. The actual number of variants placed in each region is drawn from a multinomial distribution with parameters *J/*2 (number of clustered variants) and *p*_*k*_, *k* = 1,2, … *C*/10 where *p*_1_ = *p*_2_ = ⋯ = *p*_*k*_. The other 50% of causal variants were uniformly distributed elsewhere along the genome. Having selected causal variants, their effects are drawn independently either from a N(0,1) distribution, or following a LDAK model^69^ where the effect size of variant *j* is a draw from a standard normal multiplied by a factor [2*p*_*j*_(1 − *p*_*j*_)]^−0.25^, where *p*_*j*_ is the frequency of the variant; resulting in an increased contribution from less common variants. In total this generates 3×2×2=12 scenarios; we simulated two different heritabilities 0.3, 0.6 resulting in 24 simulations in total.

Finally, to generate simulated phenotypes *Y*_*i*_ for individual *i* = 1,2, … *N*, we apply the following additive model which leverages the actual UKB genotypes and so matches properties of these data, including in particular population stratification:

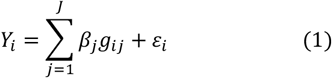

Here *β*_*j*_ and *g*_*ij*_ are the effect size and genotype of individual *i* at site *j* = 1,2, … *J*. Noise terms *ε*_*i*_ are drawn from a normal distribution with mean 0 and variance 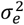. To obtain heritability *h*^2^ of 0.3 or 0.6, we set the variance of *ε*_*i*_ in (1), 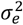 equal to 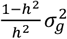, where 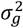 is the observed variance of the first term.

We also repeated these simulations but now allowing effect sizes to differ in the 8003 African-ancestry individuals (Supplementary Note 2). For a correlation in effect sizes of *ρ* ∈ {0.1, 0.3, 0.5, 0.7, 0.9}, we simulated from the following model:

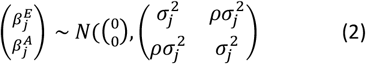

This can be done efficiently without modifying effect sizes for non-African-ancestry individuals, by noting that condition on the European effect size 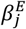 at site *j* = 1,2, … *J*, 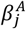 has the following distribution:

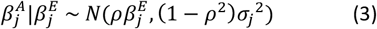

or equivalently,

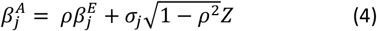

where *Z* is a standard normal random variable, and we use this to generate 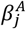 for each setting and calculate PGS for African-ancestry individuals. We adjust/do not adjust the phenotypic variance for African-ancestry individuals so as to maintain their heritability at the same value as the UKB samples as a whole. Based on the above scenarios, a simulated phenotype named as, e.g. “#causal:1K(uniform)S:0 h2:0.6” means the underlying simulation scenario for this phenotype is: a phenotype with heritability 0.6, in total 1000 causal variants, uniformly distributed along the genome, and S:0 (vs S:0.5) means there is no effect size scaling used.

## Data availability

UK and world map data can be accessed through: https://gadm.org. UK Biobank data can be downloaded by approved researchers through: https://www.ukbiobank.ac.uk. Phased haplotype data from 1000 Genome used as reference panel for HAPMIX can be accessed through: https://www.internationalgenome.org/category/data-access/. POBI was accessed using #EGAS00001000672. GWAS summary level data used in this paper can be queried using the interface implemented by “mrcieu/ieugwasr”: https://gwas.mrcieu.ac.uk and through Open Target: https://www.opentargets.org. HapMap3 site list can be accessed at: ftp://ftp.ncbi.nlm.nih.gov/hapmap/. All genetic data used in constructing the ancestry pipeline is provided by third parties and is available for use by others. Variant Frequency information for every SNP in each population is available at the University of Bristol data repository, data.bris, at https://doi.org/10.5523/bris.3g5oatl682kz82as80jakjrq91. All other resources were downloaded from their respective websites without registration requirements. The following files have been returned to UK Biobank so that they might be made available to other researchers (a) ACs on all UK Biobank participants (b) population specific allele frequency estimates for 25M variants (c) (Mean-centred) European/African genotypes and local ancestry of 8003 UKB African ancestry individuals (including the variants annotation information) (d) European/African PGS for 8003 African ancestry individuals across 29 UKB phenotypes.

## Code availability

“ANCHOR” software package is available at : https://github.com/MyersGroup/ANCHOR. Analysis scripts can be downloaded at: http://github.com/fuopen/UKB_anc. External software/packages used in this study are available as follows: “pcapred.largedata”: https://github.com/danjlawson/pcapred.largedata ; “*pcapred*” https://github.com/danjlawson/pcapred;

## Acknowledgements

This research has been conducted using the UK Biobank Resource under Application Number 27960. We thank all the participants of the UK Biobank project. POPRES was accessed using #16508: “Admixture and Selection for Fine-Scale Population Genetics of Europe”. This work was supported by the Welcome Trust grant no. 200186/Z/15/Z awarded to JM, SM and GH. L.A.F.F. is supported by the Wellcome Trust grant no. 222334/Z/21/Z. Computations used the high performance computing facilities at the Department of Statistics, University of Oxford and Oxford Biomedical Research Computing (BMRC) facility, a joint development between the Welcome Centre for Human Genetics and the Big Data Institute supported by Health Data Research UK and the NIHR Oxford Biomedical Research Centre. This work was carried out using the computational facilities of the Advanced Computing Research Centre, University of Bristol - http://www.bris.ac.uk/acrc.

## Contributions

S.R.M., D.J.L., J.M. and G.H. designed the study. S.H., G.H., J.M., D.J.L. and S.R.M. developed the methods. S.H., G.H., J.M., D.J.L. and S.R.M performed the main analyses with contributions for specific analyses from L.F. and S.S. S.H., G.H., J.M., D.J.L. and S.R.M. wrote the manuscript.

## Supplementary figures

**Figure S1.**
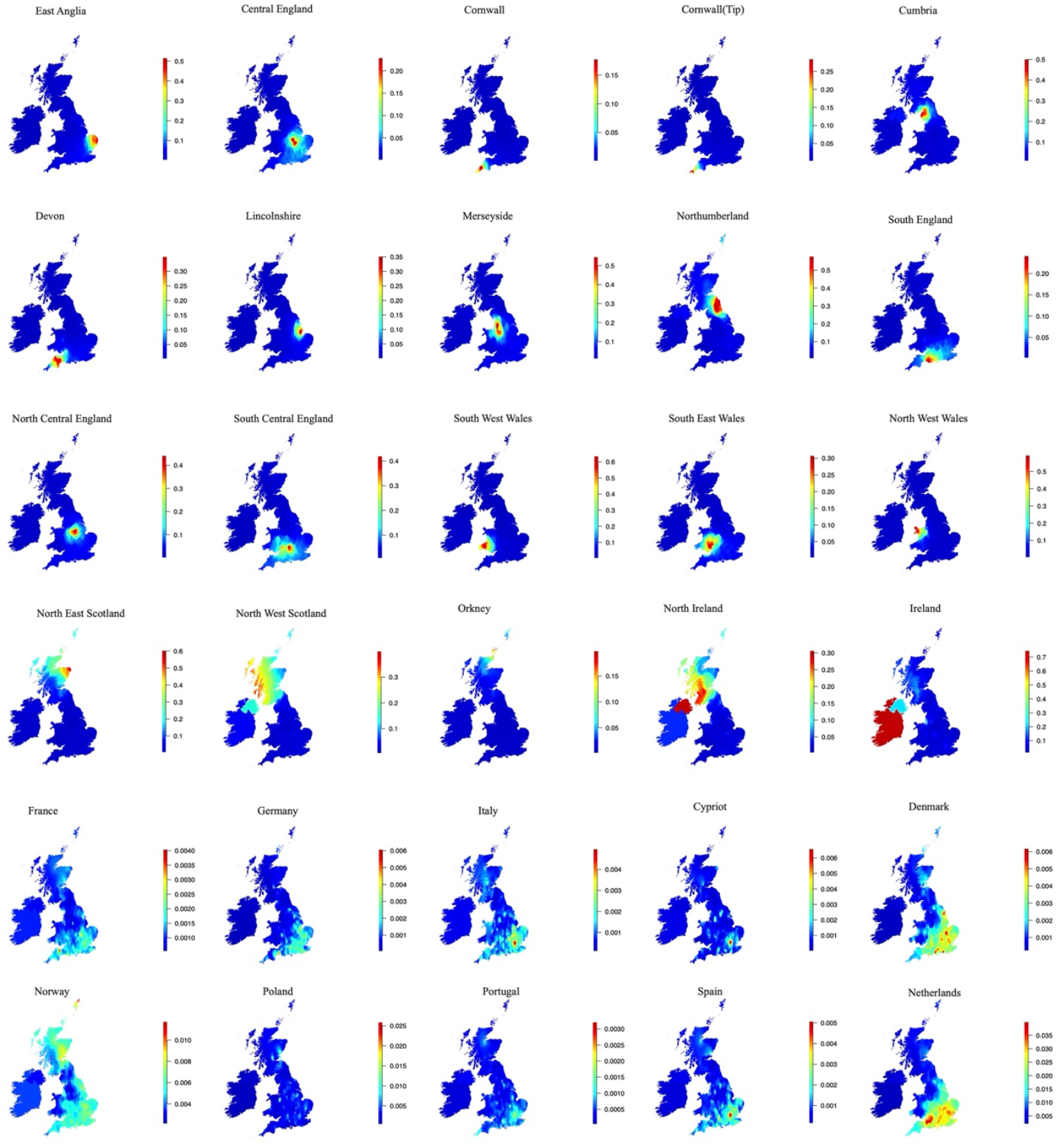
Heatmap of regional mean ancestry proportions of 30 UK+ EU ancestries, based on birthplace among UK-born and Ireland-born UKB participants. One panel per ancestry, brightest red colour on maps indicates highest ancestry, darkest blue lowest ancestry.

**Figure S2.**
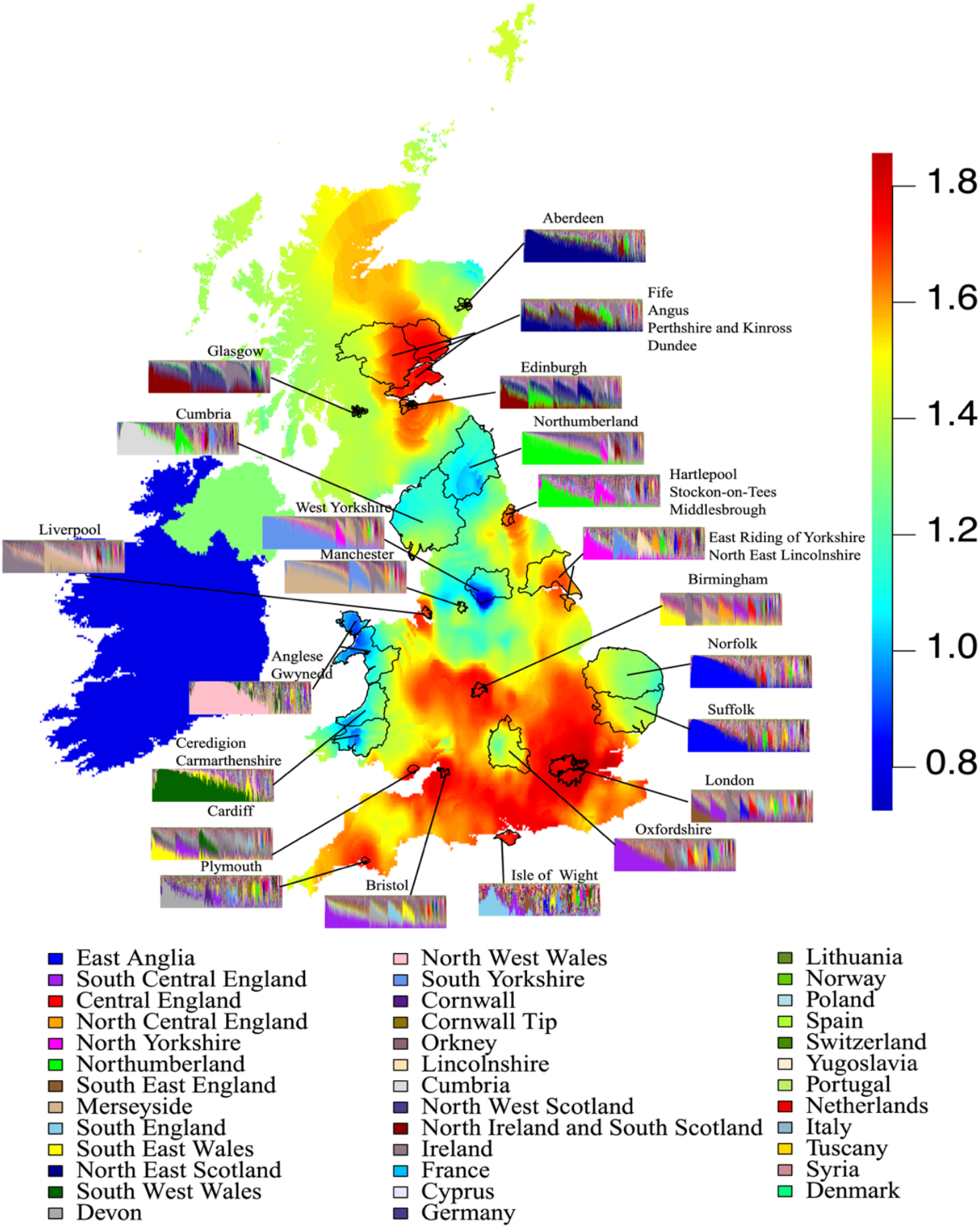
Heatmap of regional mean entropy statistic across the UK and Republic of Ireland (Methods), indicating varying degrees of admixture in the UK. Areas with high entropy are coloured by red and areas with low entropy are coloured by blue. Selected regions of high or low entropy are highlighted by the boundaries, with the barplots detailing ancestry decomposition for each such region.

**Figure S3.**
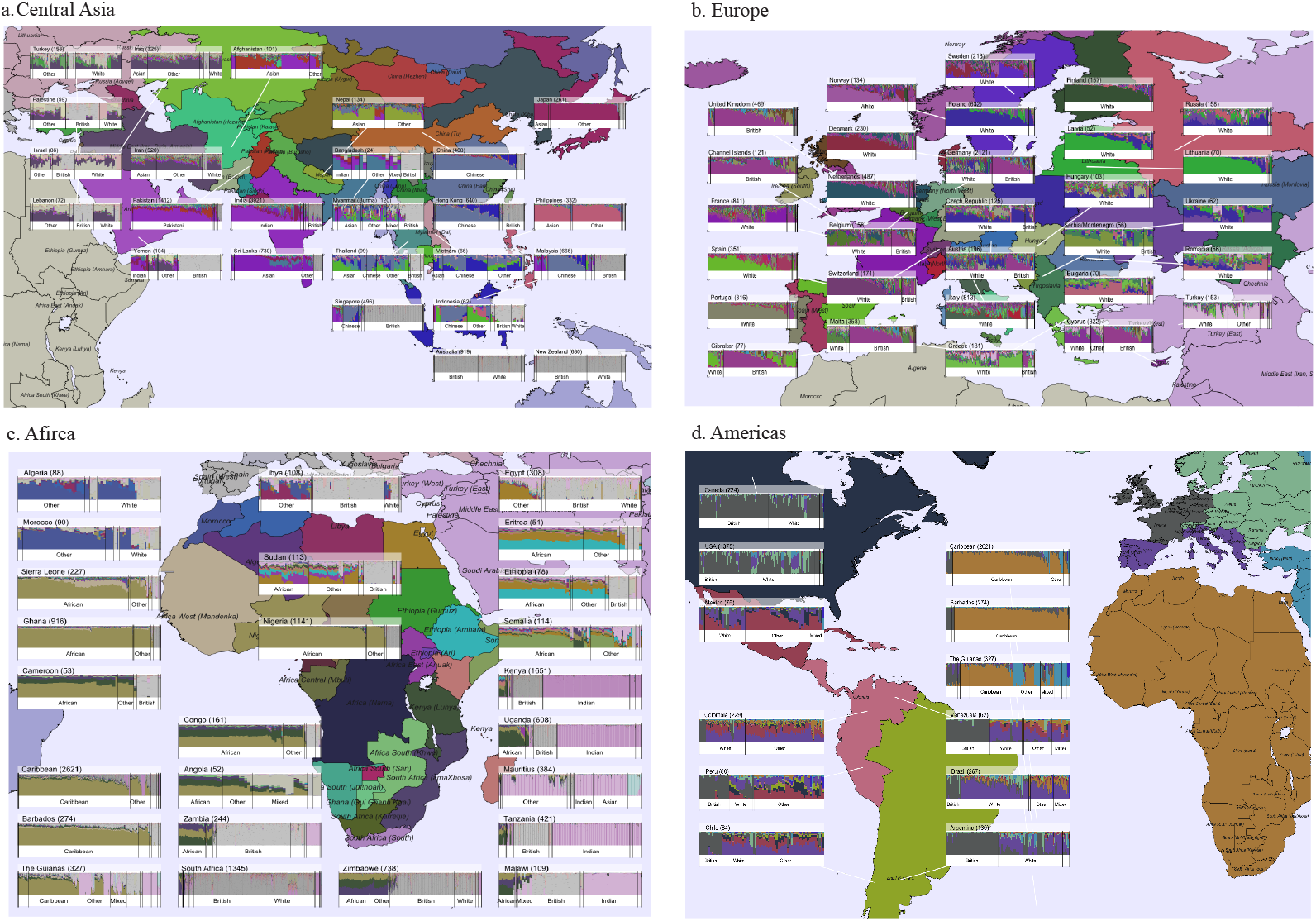
World ancestries in the UKB inferred by the ancestry pipeline. UKB participants are mapped to their self-reported country of birth. World countries are coloured by different colours if they are present in the pipeline a). Ancestries in Central Asia; b). Ancestries in Europe; c). Ancestries in Africa; d). Ancestries in the Americas.

**Figure S4.**
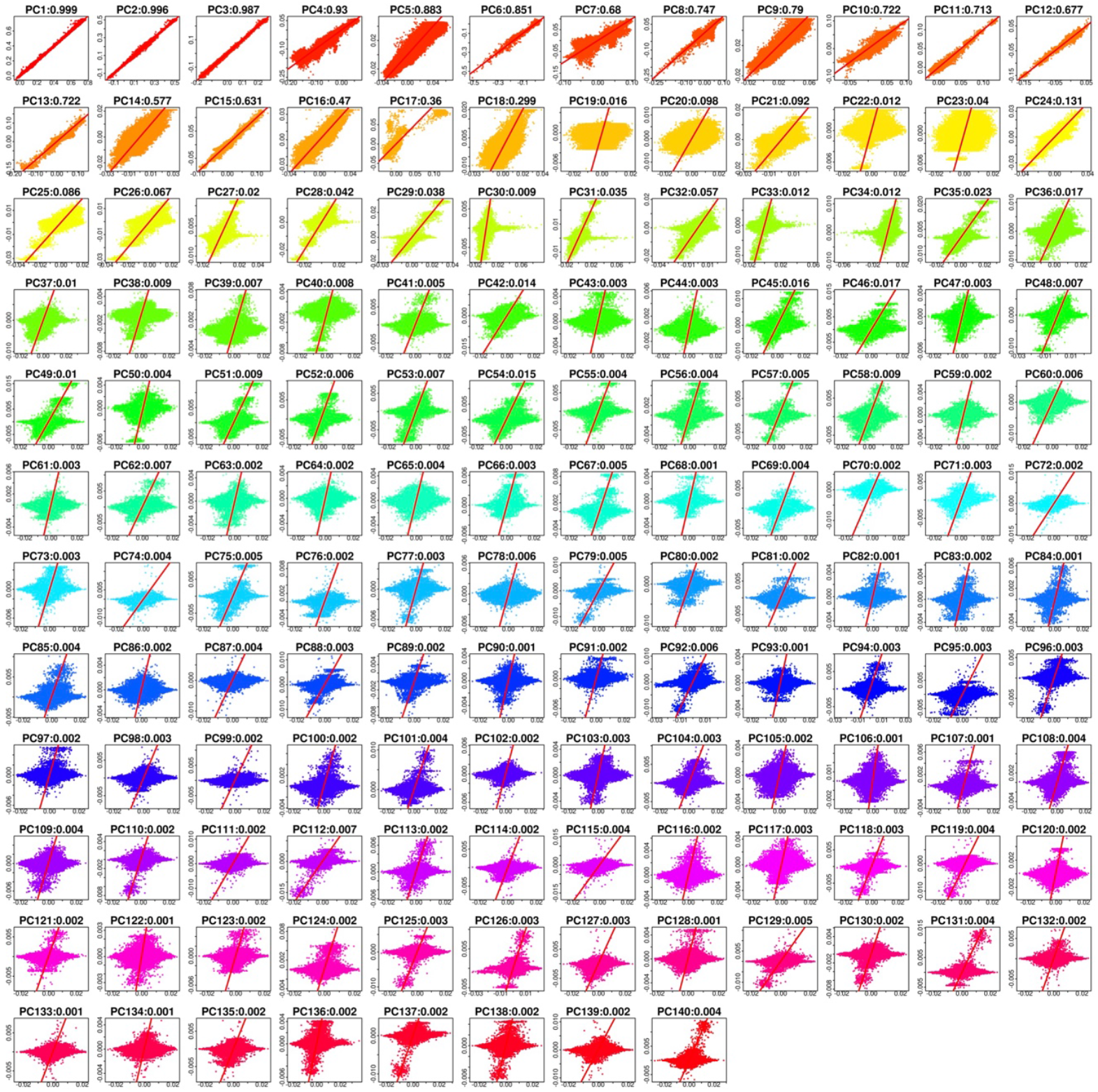
Using 127 ACs to predict 140 PCs across all UKB participants. The X-axis shows observed PCs, and the y-axis shows AC-predicted PCs.

**Figure S5.**
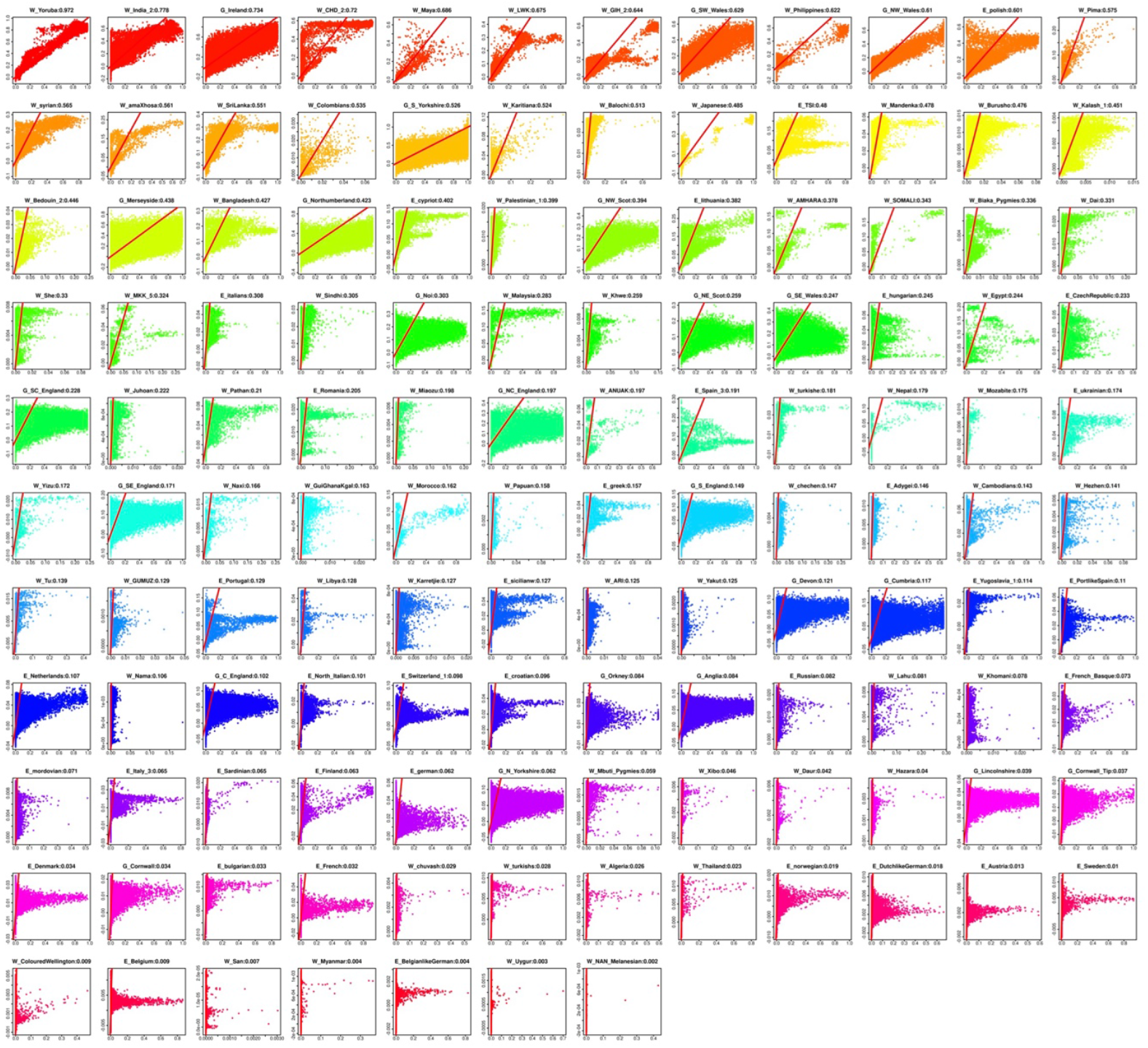
Using 140 PCs to predict 127 ACs. The X-axis shows observed ACs, and the y-axis shows PC-predicted ACs. Plots are arranged by the R^2^ between the real and predicted ACs, in descending order.

**Figure S6.**
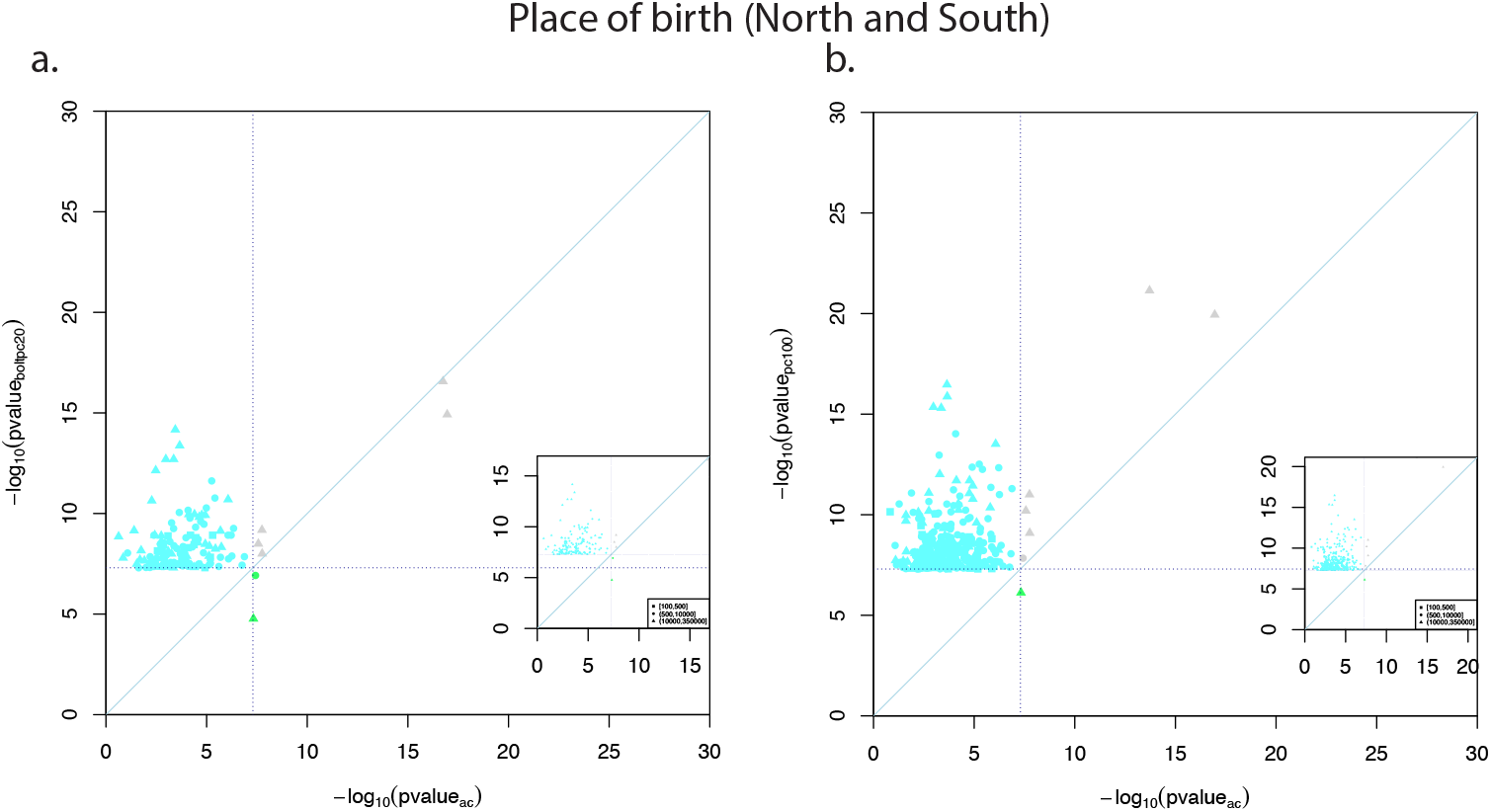
Comparison of independent GWAS hits for Place of birth (North/South) between GWAS corrected by ACs and by a) BOLT-LMM GWAS including 20 PCs as regressors; b) 100PCs. Light blue colours represent independent genome wide significant variants identified by each respective GWAS; green coloured points represent independent genome wide significant variants identified by AC-corrected GWAS. Dot shape is chosen by the minor allele count in the sample.

**Figure S7.**
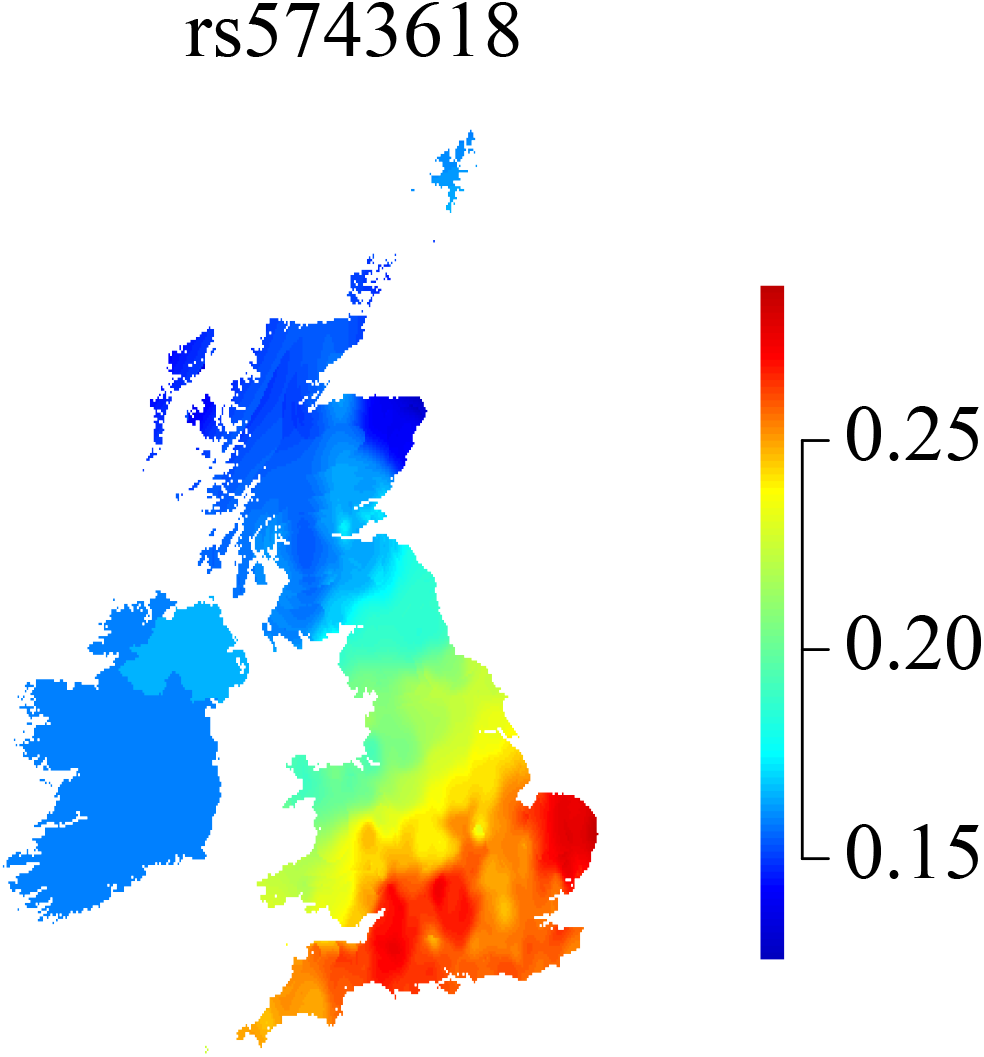
Heatmap of regional mean allele frequency of the SNP rs5743618, based on genotype and birthplace among UK-born and Ireland-born UKB participants.

**Figure S8.**
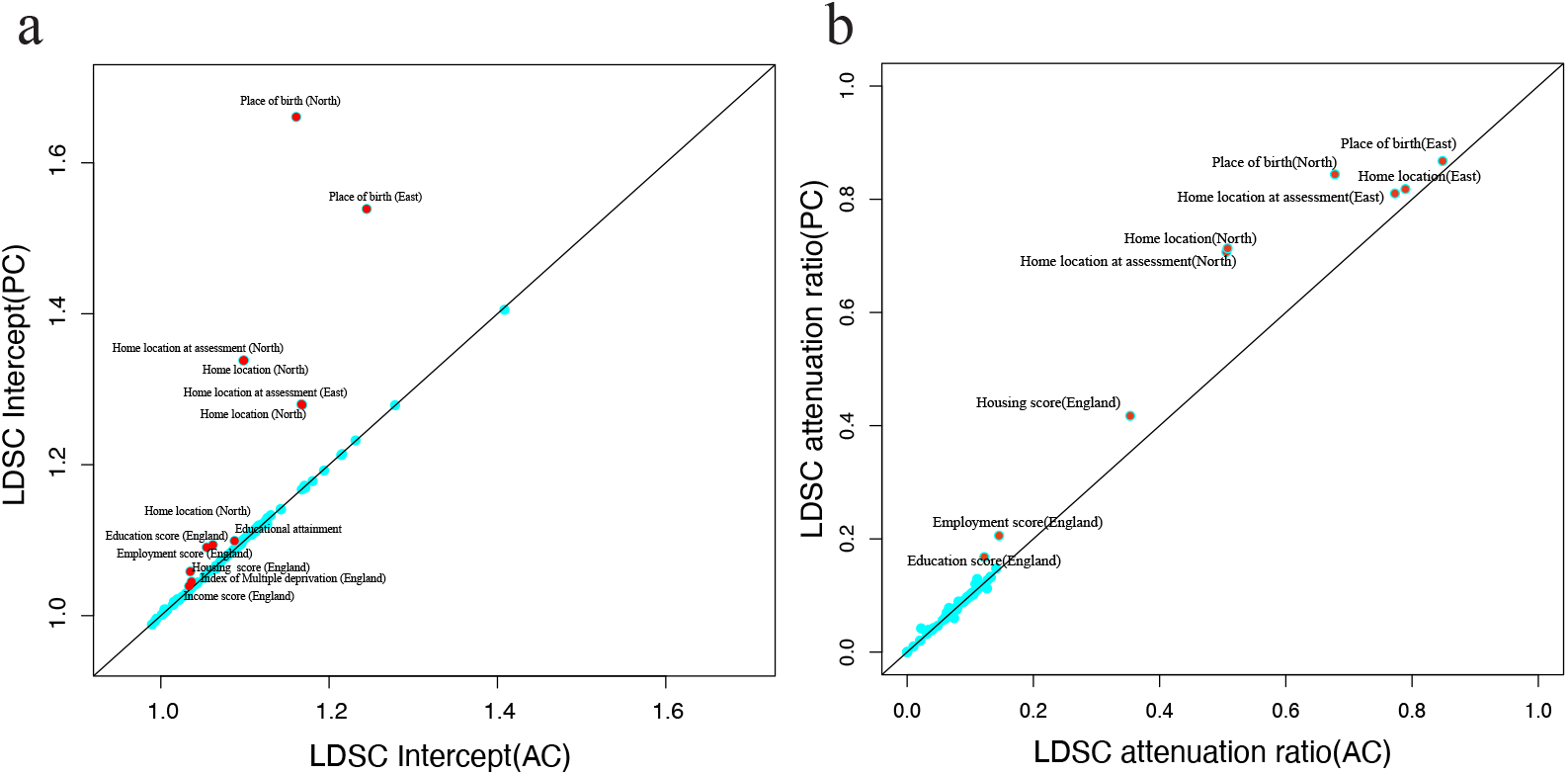
Comparison of inflation between ACs-corrected GWAS and PC-corrected GWAS across 99 quantitative traits by running LD score regression (LDSC) a). Comparison of the LDSC intercept between AC-corrected GWAS (x-axis) and PC-corrected GWAS (y-axis) where red dots indicate phenotypes with intercept difference between PC-corrected GWAS and AC-corrected GWAS >0.005; b). Comparison of the LDSC attenuation ratio between AC-corrected GWAS (x-axis) and PC-correct GWAS (y-axis), where red dots indicate phenotypes with difference of attenuation ratio between PC-corrected GWAS and AC-corrected GWAS >0.02.

**Figure S9.**
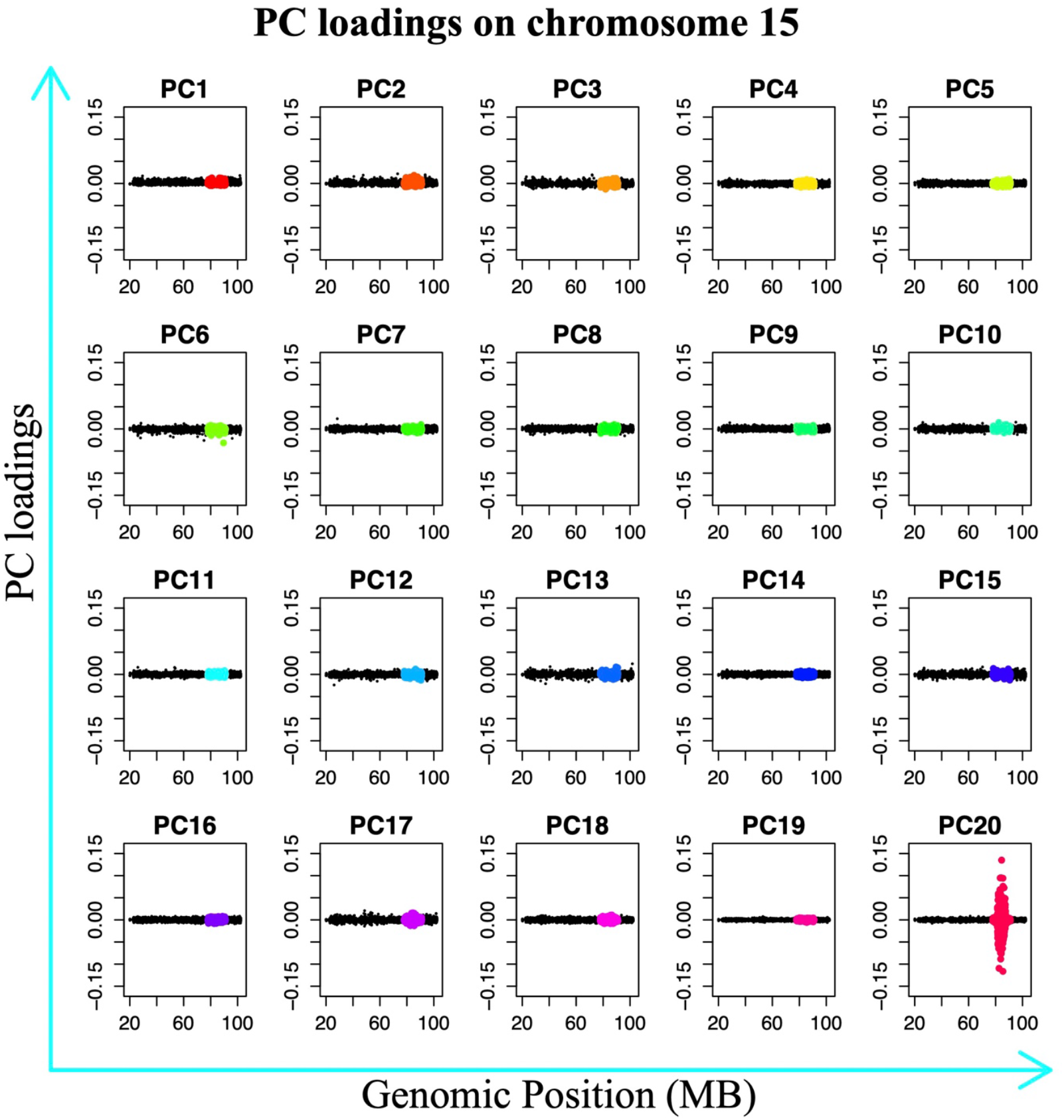
SNP loadings of the first 20 PCs, along Chromosome 15. For PC20, high loadings occur in the region 80-90 megabases.

**Figure S10.**
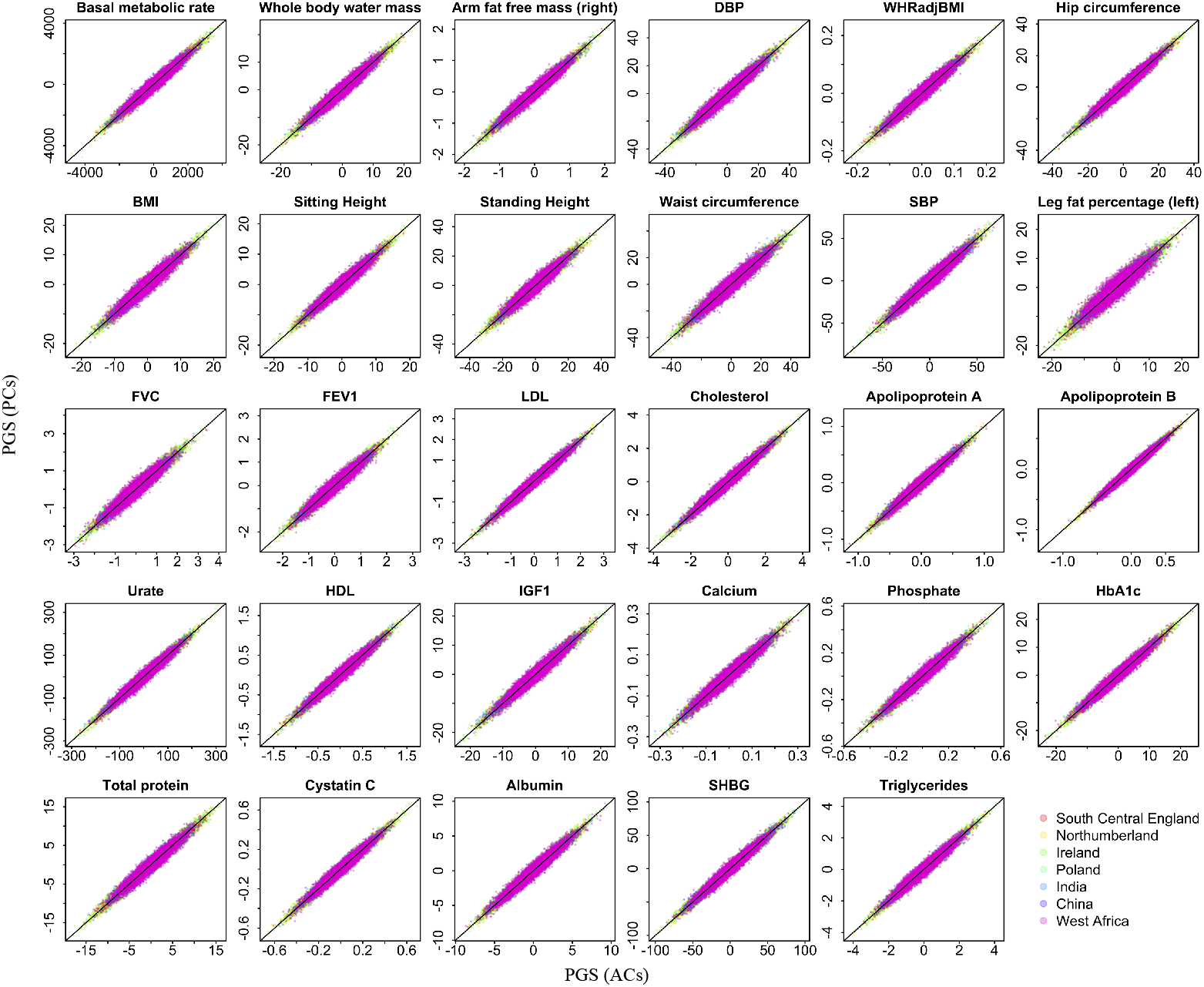
PGSs of 29 traits constructed by both AC-corrected GWAS and PC-corrected GWAS on UKB participants, in 7 ancestral groups: South Central England, Northumberland, Ireland, Poland, India, China and West Africa. X-axes represent the PGS constructed by AC-corrected GWAS and y-axes represent the PGS constructed by PC-corrected GWAS. Each PGS was centralised by regressing out age and sex prior to plotting.

**Figure S11.**
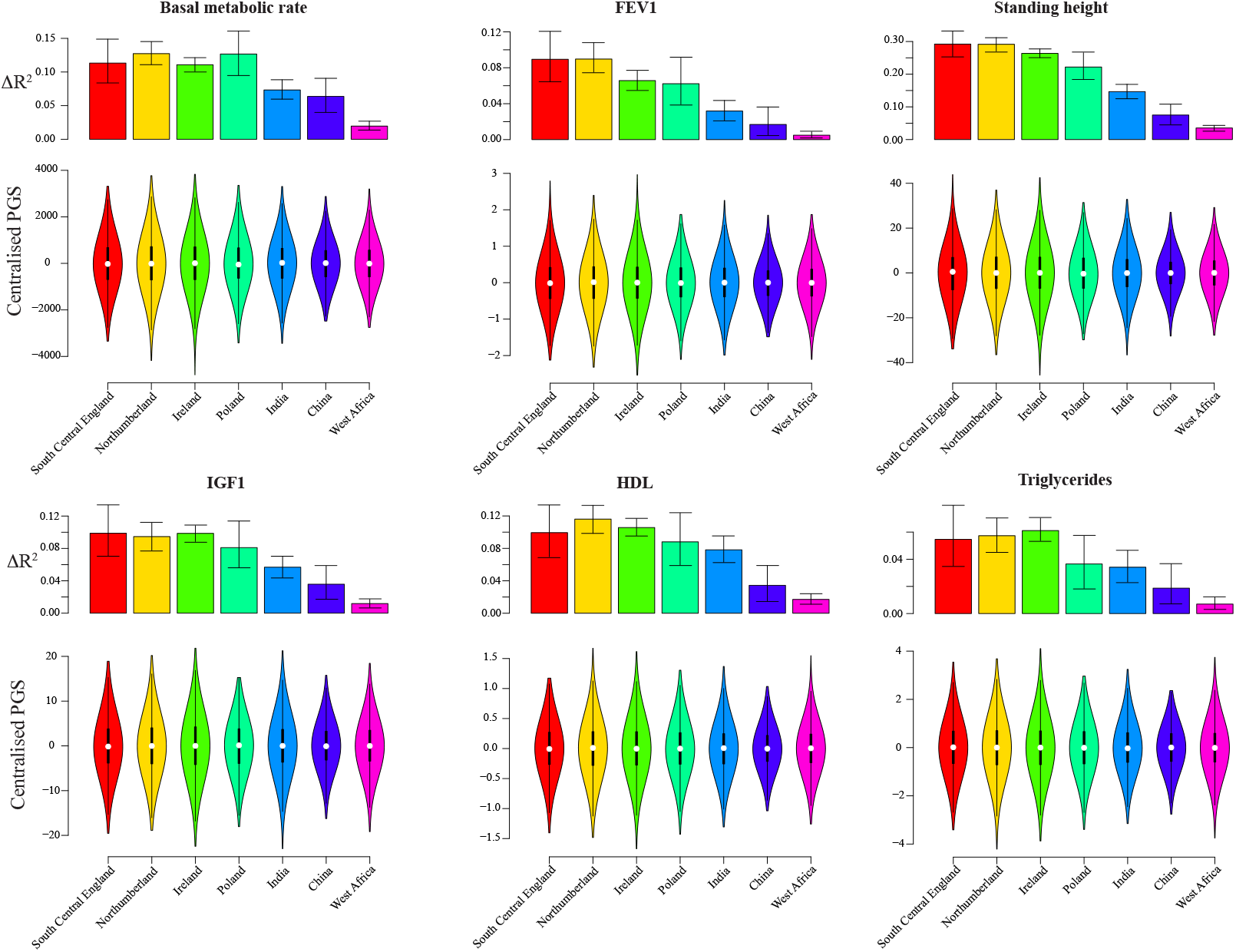
ΔR^2^ (Methods) and distribution of PGSs in 7 ancestral groups across 6 out of 29 traits, using AC-corrected GWAS. Top: ΔR^2^ of PGSs in each ancestry group; bottom: the violin plots show the distribution of PGSs, across different ancestral groups. 95% confidence intervals for each bar is calculated by bootstrap using 1000 iterations.

**Figure S12.**
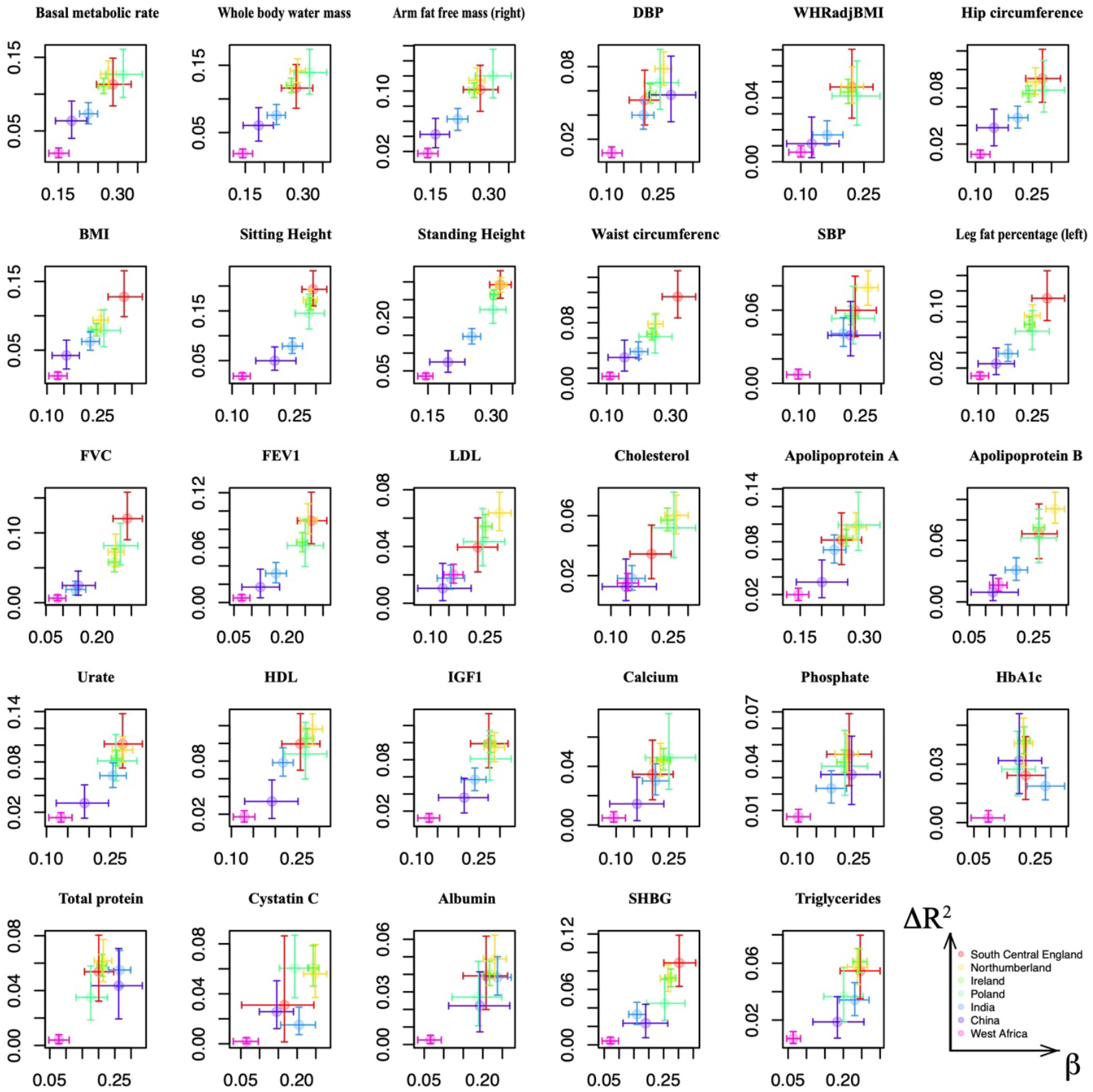
Comparison of effect size estimate of PGSs β and ΔR^2^, between 7 ancestral groups across 29 traits, using AC-corrected GWAS. In most cases, β mirrors ΔR^2^. 95% confidence intervals for each dot are calculated by bootstrap with 1000 iterations.

**Figure S13.**
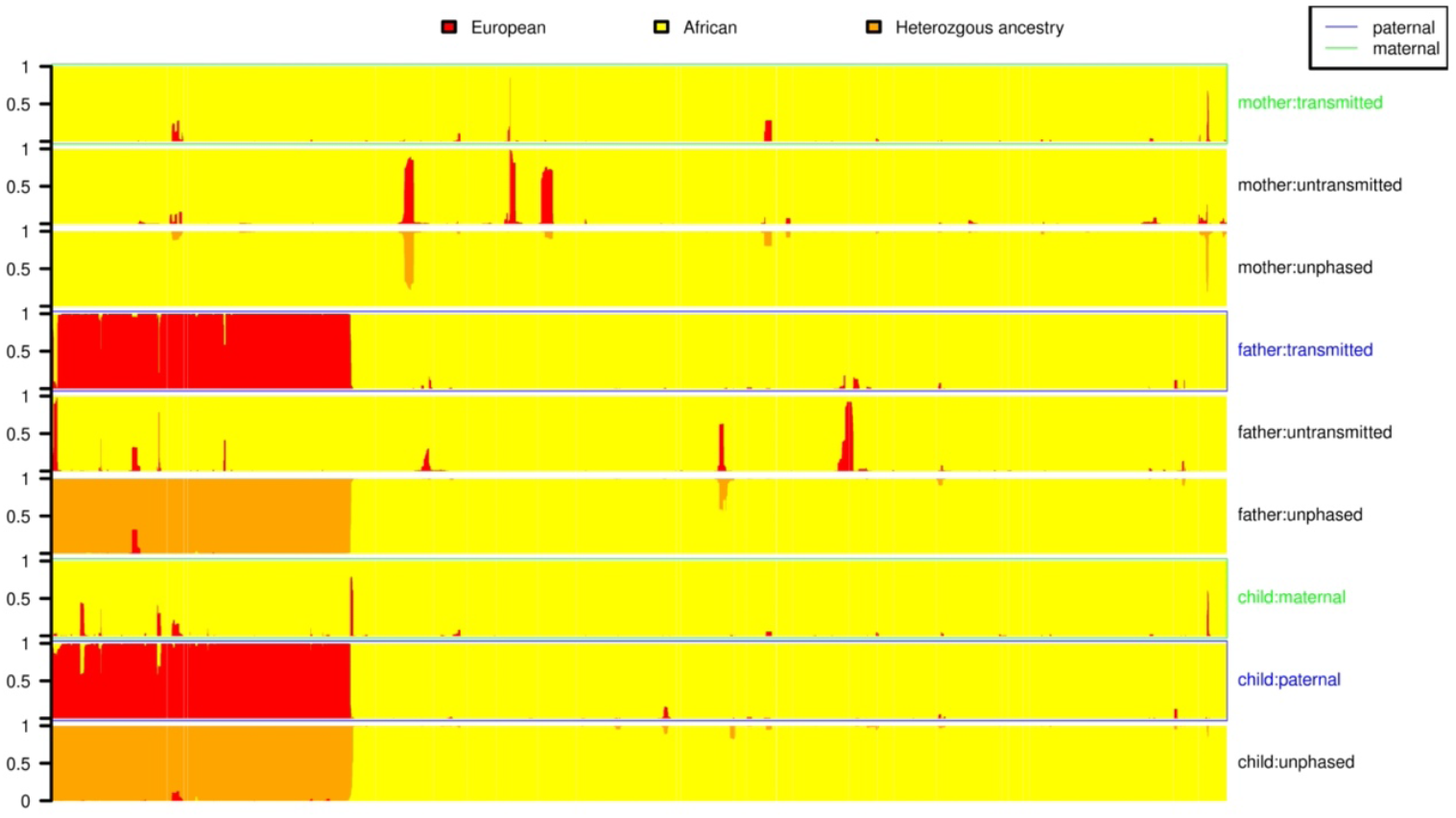
Ancestry-aware phasing of chromosome 4 in trio individuals. From bottom to top, the first three bars represent ancestry inferred using unphased genotypes, or based on the phased paternal and then maternal haplotypes for the child in the trio; the second three bars respectively represent ancestry inferred using the unphased genotypes, un-transmitted and transmitted haplotypes for the father in the trio; the last three bars are the same, but for the mother in the trio. Ancestry inference using unphased and phased data are similar, and African (coloured in yellow), or European genome segment (red) are generally large (megabases in size).

**Figure S14.**
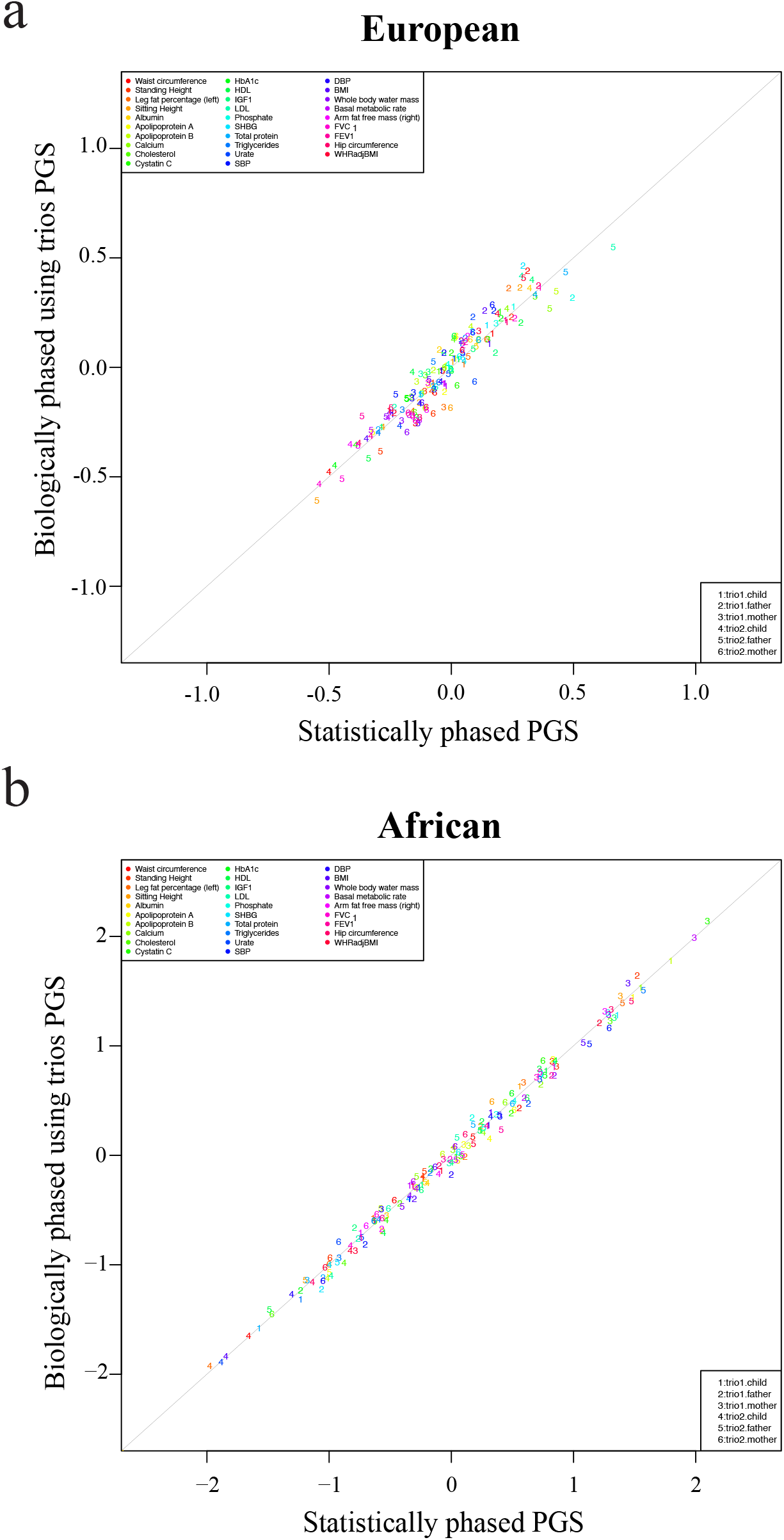
Comparison between PGS in trios calculated using either unphased genotypes (x-axis) or trio-phased haplotypes (y-axis) for 6 individuals from 2 trios, across 29 traits. For details of the normalisation performed to enable joint comparison of these PGS, see “Using trios to conduct ancestry aware phasing for PGSs”. a). comparison of unphased and phased EPGSs, calculated using only inferred European segments; b). comparison of unphased and phased APGSs, calculated using only inferred African segments. Note that across traits and individuals, the PGS are very similar.

**Figure S15.**
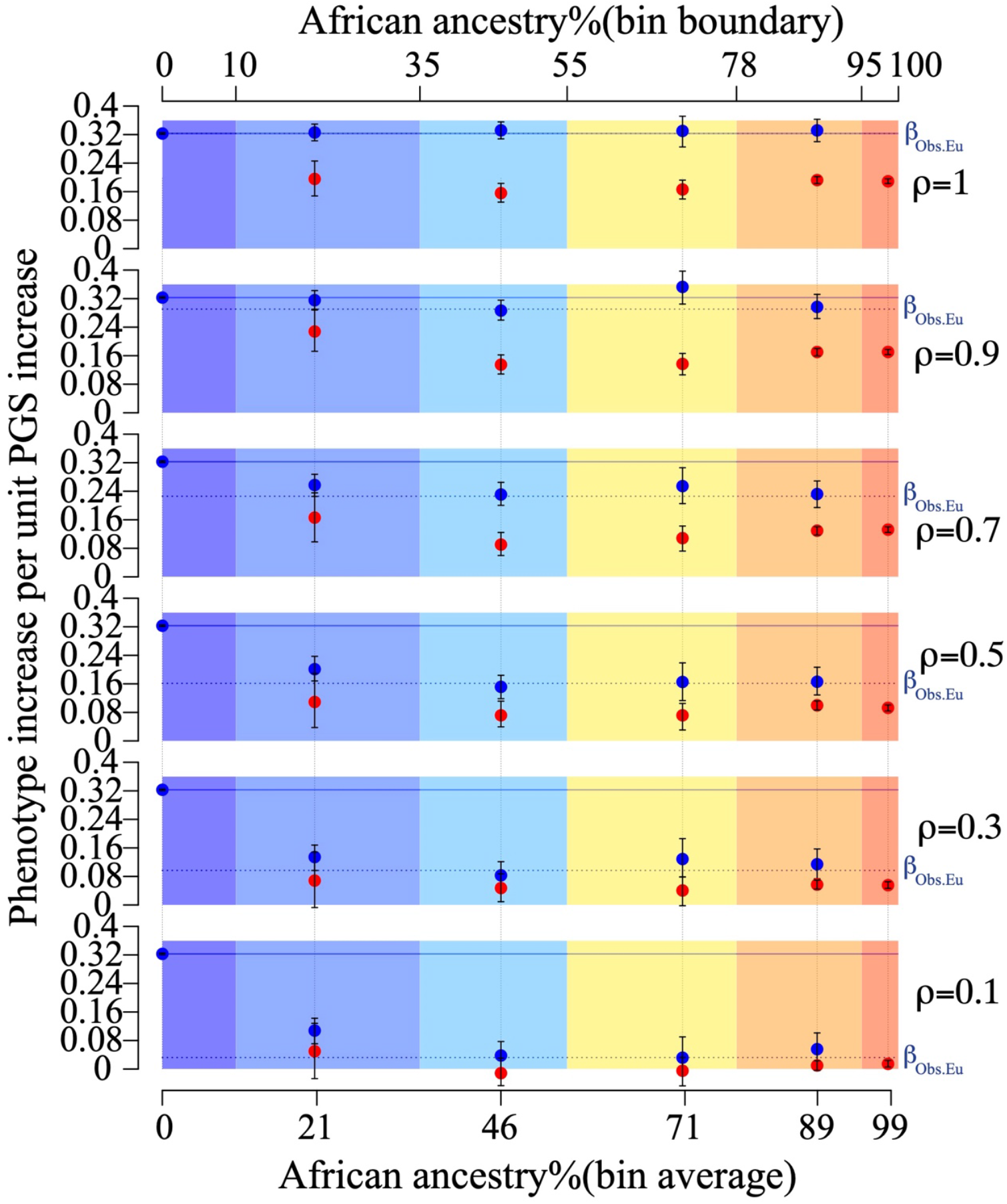
Results of application of ANCHOR for 24 UKB simulated traits, for individuals of varying African ancestry binned as in main Figure 4c (x-axis; coloured region). For each bin in each panel, estimates of coefficients β_Eu_ (blue) and β_Af_ (red) are shown with 95% bootstrapped confidence intervals, representing the increase in phenotype per unit increase in PGS within European and African genomic regions respectively (Methods). Also shown are these estimates from individuals of ∼100% European (β_Obs.Eu_; blue horizontal bar) or ∼100% African (red horizontal bar) ancestry. From top to bottom, each panel represents descending correlation ρ. The case ρ=1, where only local effects impact portability, corresponds to blue points lying along the blue line, and similarly for red points, as observed. In cases when ρ<1, the dotted horizontal line is plotted at β_Obs.Eu_ x ρ, and the blue points are predicted to lie along this line if β_Eu_/ β_Obs.Eu_ provides an accurate estimate of ρ, as predicted by theory (Methods; Supplementary Note 2).

**Figure S16.**
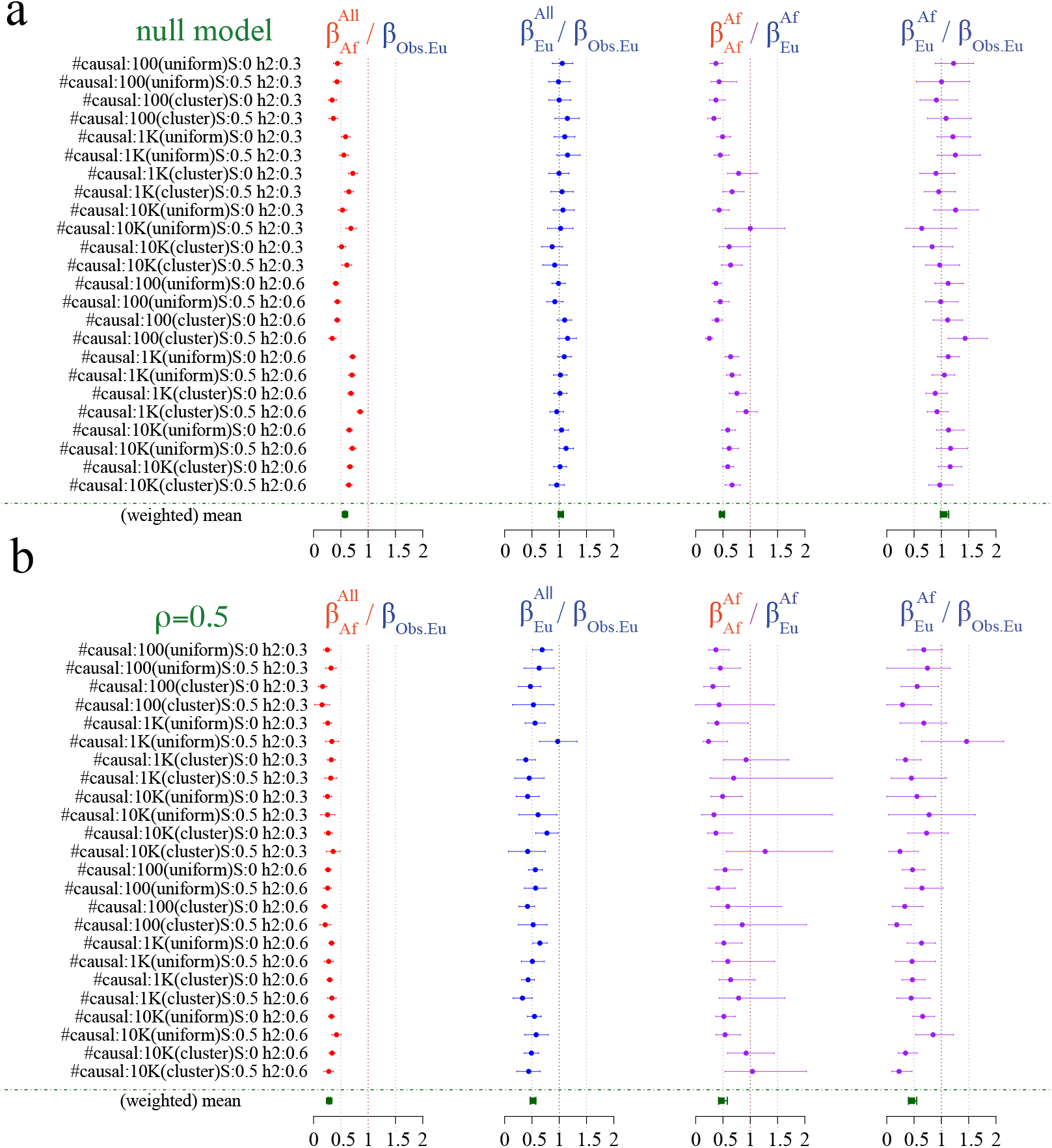
ANCHOR results for each simulated trait. Columns show ratios of estimates defined as in Main Figure 4e, and rows show simulated phenotypes. The final row shows combined estimates. The second and fourth columns estimate the underlying correlation in causal effect sizes between European-ancestry and either all 8003 African-ancestry individuals, or individuals of 100% African-ancestry, respectively, while the first and third columns additionally estimate the impact of local effects on predictive power for African-ancestry segments in the same two settings. a) Results under the null model ρ=1, and b) results under the model ρ=0.5, with incomplete correlation in underlying effects sizes between different populations.

**Figure S17.**
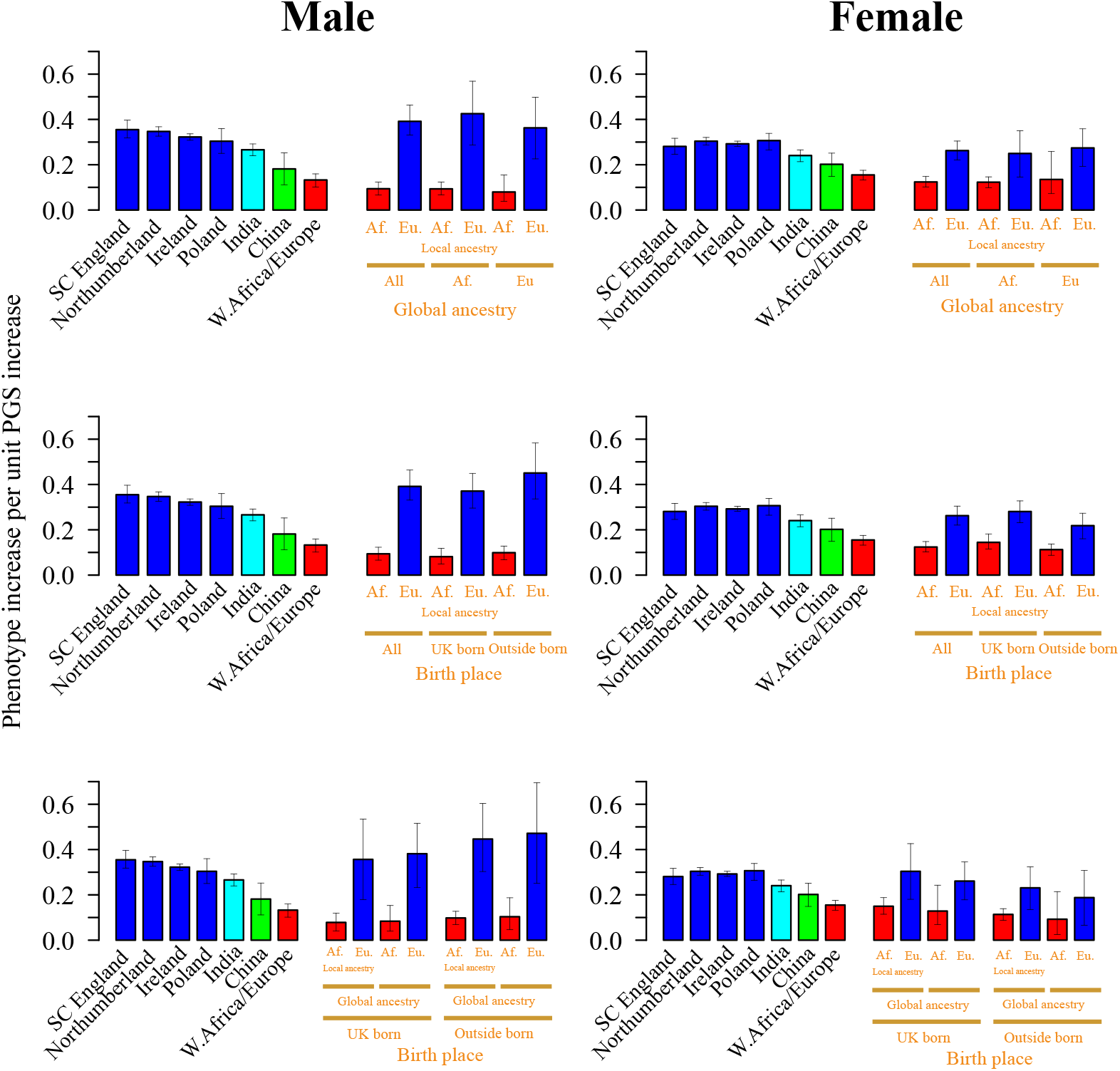
Investigation of impact of gender and UK vs non-UK birthplace on the relationship between PGS performance and ancestry, for the trait of height. For each plot, details are as described in the legend for Figure 4d unless stated otherwise. Left column: analysis results for only male African-ancestry individuals; right column, female individuals. Top row: as Figure 4d. Middle row: the last four columns in each plot estimate the estimated increase in phenotype per unit increase in the EPGS (blue) or APGS (red) polygenic score, separately for UK-born and non-UK-born individuals (note there is no significant difference in either case). Bottom row: as for the first row, but now applying the ANCHOR model to UK-born or non-UK-born individuals of the specified gender and varying ancestry (see section “Joint model fitting by accounting for other environment factors”). Once again none of the bootstrapped CIs indicate significant impacts of birthplace on results; and neither gender shows significant evidence of different β_Eu_/ β_Obs.Eu_ ratios (distinct effect sizes) for African-ancestry (right-hand blue bars) vs UK-ancestry (left hand two blue bars) individuals, indicating results are consistent with ρ=1, i.e. equal effect sizes in either gender and regardless of birthplace and ancestry. (There are obvious differences comparing males and females, corresponding to their different mean heights.)

**Figure S18.**
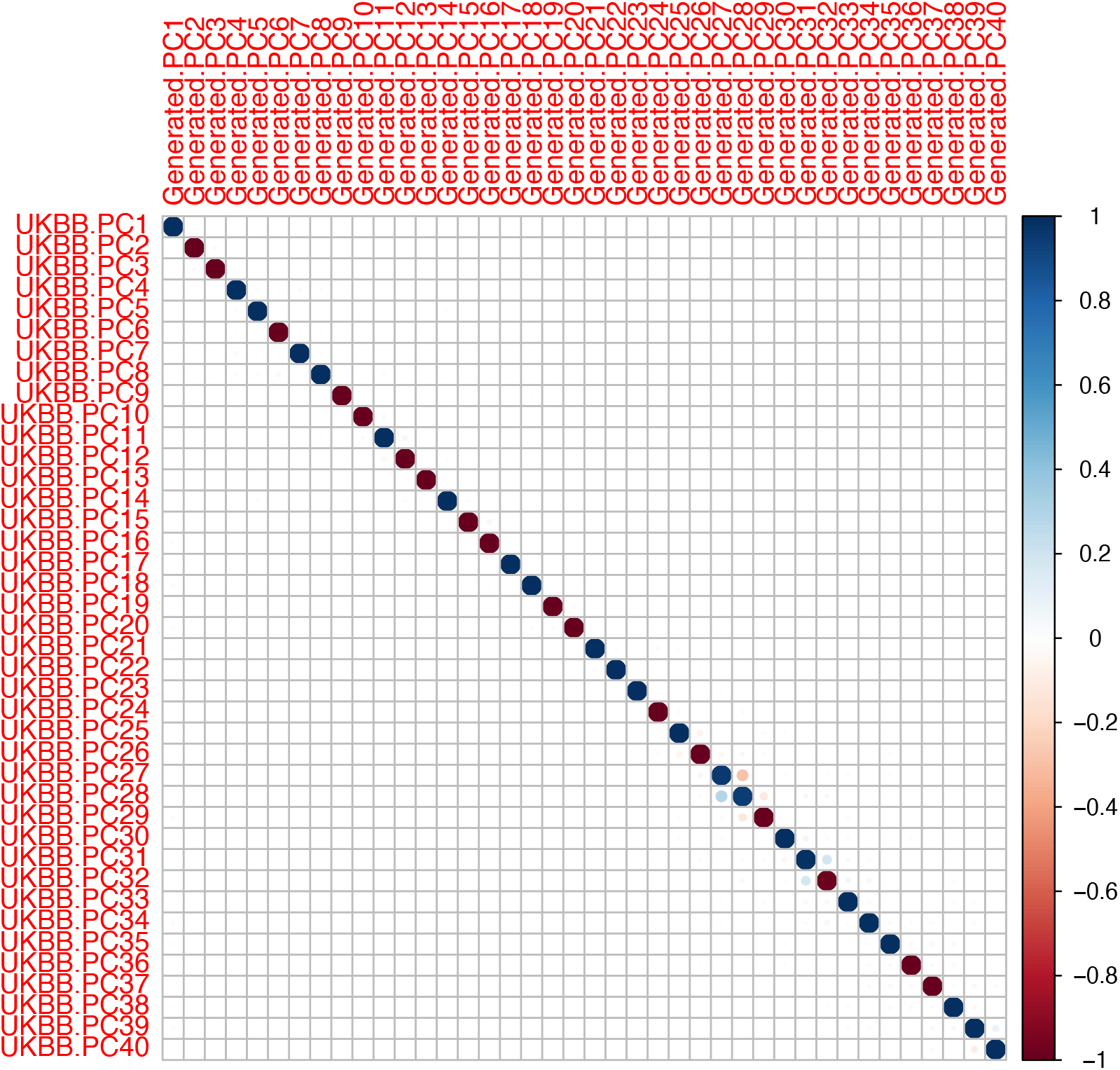
Comparison of 40PCs released by UKB, and 40PCs generated as described in Methods, to verify very high correlations (either close to 1, or close to minus 1; the sign of each PC is arbitrary).

