## Supplementary Note 1 for "Leveraging fine-scale population structure reveals conservation in genetic effect sizes between human populations across a range of human phenotypes"

#### Legends of supplementary figures and tables.

#### Supplementary figure legends

Figure S1. Heatmap of regional mean ancestry proportions of 30 UK+ EU ancestries, based on birthplace among UK-born and Ireland-born UKB participants. One panel per ancestry, brightest red colour on maps indicates highest ancestry, darkest blue lowest ancestry.

Figure S2. Heatmap of regional mean entropy statistic across the UK and Republic of Ireland (Methods), indicating varying degrees of admixture in the UK. Areas with high entropy are coloured by red and areas with low entropy are coloured by blue. Selected regions of high or low entropy are highlighted by the boundaries, with the barplots detailing ancestry decomposition for each such region.

Figure S3. World ancestries in the UKB inferred by the ancestry pipeline. UKB participants are mapped to their self-reported country of birth. World countries are coloured by different colours if they are present in the pipeline a). Ancestries in Central Asia; b). Ancestries in Europe; c). Ancestries in Africa; d). Ancestries in the Americas.

Figure S4. Using 127 ACs to predict 140 PCs across all UKB participants. The X-axis shows observed PCs, and the y-axis shows AC-predicted PCs.

Figure S5. Using 140 PCs to predict 127 ACs. The X-axis shows observed ACs, and the y-axis shows PC-predicted ACs. Plots are arranged by the $R^2$ between the real and predicted ACs, in descending order.

Figure S6. Comparison of independent GWAS hits for Place of birth (North/South) between GWAS corrected by ACs and by a) BOLT-LMM GWAS including 20 PCs as regressors; b) 100PCs. Light blue colours represent independent genome wide significant variants identified by each respective GWAS; green coloured points represent independent genome wide significant variants identified by AC-corrected GWAS. Dot shape is chosen by the minor allele count in the sample.

Figure S7. Heatmap of regional mean allele frequency of the SNP rs5743618, based on genotype and birthplace among UK-born and Ireland-born UKB participants.

Figure S8. Comparison of inflation between ACs-corrected GWAS and PC-corrected GWAS across 99 quantitative traits by running LD score regression (LDSC) a). Comparison of the LDSC intercept between AC-corrected GWAS (x-axis) and PC-corrected GWAS (y-axis) where red dots indicate phenotypes with intercept difference between PC-corrected GWAS and AC-corrected GWAS >0.005; b). Comparison of the LDSC attenuation ratio between AC-corrected GWAS (x-axis) and PC-correct GWAS (y-axis), where red dots indicate phenotypes with difference of attenuation ratio between PC-corrected GWAS and AC-corrected GWAS >0.02.

Figure S9. SNP loadings of the first 20 PCs, along Chromosome 15. For PC20, high loadings occur in the region 80-90 megabases.

Figure S10. PGSs of 29 traits constructed by both AC-corrected GWAS and PC-corrected GWAS on UKB participants, in 7 ancestral groups: South Central England, Northumberland, Ireland, Poland, India, China and West Africa. X-axes represent the PGS constructed by AC-corrected GWAS and y-axes represent the PGS constructed by PC-corrected GWAS. Each PGS was centralised by regressing out age and sex prior to plotting.

Figure S11. $\Delta R^2$ (Methods) and distribution of PGSs in 7 ancestral groups across 6 out of 29 traits, using AC-corrected GWAS. Top: $\Delta R^2$ of PGSs in each ancestry group; bottom: the violin plots show the distribution of PGSs, across different ancestral groups. 95% confidence intervals for each bar is calculated by bootstrap using 1000 iterations.

Figure S12. Comparison of effect size estimate of PGSs $\beta$ and $\Delta R^2$, between 7 ancestral groups across 29 traits, using AC-corrected GWAS. In most cases, $\beta$ mirrors $\Delta R^2$. 95% confidence intervals for each dot are calculated by bootstrap with 1000 iterations.

Figure S13. Ancestry-aware phasing of chromosome 4 in trio individuals. From bottom to top, the first three bars represent ancestry inferred using unphased genotypes, or based on the phased paternal and then maternal haplotypes for the child in the trio; the second three bars respectively represent ancestry inferred using the unphased genotypes, un-transmitted and transmitted haplotypes for the father in the trio; the last three bars are the same, but for the mother in the trio. Ancestry inference using unphased and phased data are similar, and African (coloured in yellow), or European genome segment (red) are generally large (megabases in size).

Figure S14. Comparison between PGS in trios calculated using either unphased genotypes (x-axis) or trio-phased haplotypes (y-axis) for 6 individuals from 2 trios, across 29 traits. For details of the normalisation performed to enable joint comparison of these PGS, see "Using trios to conduct ancestry aware phasing for PGSs". a). comparison of unphased and phased EPGSs, calculated using only inferred European segments; b). comparison of unphased and phased APGSs, calculated using only inferred African segments. Note that across traits and individuals, the PGS are very similar.

Figure S15. Results of application of ANCHOR for 24 UKB simulated traits, for individuals of varying African ancestry binned as in main Figure 4c (x-axis; coloured region). For each bin in each panel, estimates of coefficients $\beta_{Eu}$ (blue) and $\beta_{Af}$ (red) are shown with 95% bootstrapped confidence intervals, representing the increase in phenotype per unit increase in PGS within European and African genomic regions respectively (Methods). Also shown are these estimates from individuals of ~100% European ($\beta_{Obs.Eu}$; blue horizontal bar) or ~100% African (red horizontal bar) ancestry. From top to bottom, each panel represents descending correlation ρ. The case ρ=1, where only local effects impact portability, corresponds to blue points lying along the blue line, and similarly for red points, as observed. In cases when ρ<1, the dotted horizontal line is plotted at $\beta_{Obs.Eu} \times \rho$, and the blue points are predicted to lie along this line if $\beta_{Eu}/\ \beta_{Obs.Eu}$ provides an accurate estimate of ρ, as predicted by theory (Methods; Supplementary Note 2)..

Figure S16. ANCHOR results for each simulated trait. Columns show ratios of estimates defined as in Main Figure 4e, and rows show simulated phenotypes. The final row shows combined estimates. The second and fourth columns estimate the underlying correlation in causal effect sizes between European-ancestry and either all 8003 African-ancestry individuals, or individuals of 100% African-ancestry, respectively, while the first and third columns additionally estimate the impact of local effects on predictive power for African-ancestry segments in the same two settings. a) Results under the null model $\rho=1$, and b) results under the model $\rho=0.5$, with incomplete correlation in underlying effects sizes between different populations.

Figure S17. Investigation of impact of gender and UK vs non-UK birthplace on the relationship between PGS performance and ancestry, for the trait of height. For each plot, details are as described in the legend for Figure 4d unless stated otherwise. Left column: analysis results for only male African-ancestry individuals; right column, female individuals. Top row: as Figure 4d. Middle row: the last four columns in each plot estimate the estimated increase in phenotype per unit increase in the EPGS (blue) or APGS (red) polygenic score, separately for UK-born and non-UK-born individuals (note there is no significant difference in either case). Bottom row: as for the first row, but now applying the ANCHOR model to UK-born or non-UK-born individuals of the specified gender and varying ancestry (see section "Joint model fitting by accounting for other environment factors"). Once again none of the bootstrapped CIs indicate significant impacts of birthplace on results; and neither gender shows significant evidence of different $\beta_{Eu}$/ $\beta_{Obs.Eu}$ ratios (distinct effect sizes) for African-ancestry (right-hand blue bars) vs UK-ancestry (left hand two blue bars) individuals, indicating results are consistent with $\rho=1$, i.e. equal effect sizes in either gender and regardless of birthplace and ancestry. (There are obvious differences comparing males and females, corresponding to their different mean heights.)

Figure S18. Comparison of 40PCs released by UKB, and 40PCs generated as described in Methods, to verify very high correlations (either close to 1, or close to minus 1; the sign of each PC is arbitrary).

Figure S19. Comparison of inferred African ancestry obtained by ACs (x-axis) and by Hapmix (y-axis)

Figure S20. Distribution of African ancestry stratified by gender and whether individuals are born within the UK, or outside the UK.

#### Supplementary table legends

Table S1. UK and Ireland ancestries, and their abbreviations, as identified and used in the ancestry pipeline.

Table S2. Mean ancestries in each birth place region/countries from each of the 127 ancestry groups that genetic similarity is compared to.

Table S3. Properties of SNPs showing genome wide significance in AC-corrected GWAS (subtable labels) vs only in PC-corrected GWAS (alternate subtables) for two regionally defined traits: Employment score (England) and Birth place. Only those SNPs showing prior GWAS association evidence, for the traits catalogued in Table S6, are listed (this removes many SNPs in particular significant only in the PC-corrected GWAS). Note that the majority of SNPs significant after AC-correction for employment score show prior GWAS evidence for traits related to cognitive function, educational attainment, or socioeconomic status, while few PC-only-associated SNPs show prior GWAS evidence. The final subtable lists now those SNPS significant in only the AC-corrected GWAS, for waist circumference.

Table S4. Results of running a joint regression to investigate correlations with PCs, for those SNPs in Table S2 associated with waist circumference after AC-correction but not PC-correction. Each sheet indicates the results for regression of the genotype of the labelled SNP versus the first 20 PCs; note that PC20 shows a strong and dominant correlation in each case, corresponding to the high loadings of this PC in this region of the genome (Figure S9) and indicating this PC might result in false negatives.

Table S5. Correlations of (centralised) PGS constructed by AC-corrected and PC-corrected GWAS for 7 ancestry groups across 29 traits.

Table S6. Mean phenotypes and mean PGS constructed by AC-corrected and PC-corrected GWAS for 7 ancestry groups across 29 traits.

Table S7. Demographic and phenotypic UKB fields used in the AC-corrected and PC-correct GWAS.

Table S8. Queried traits for GWAS hits in AC and/or PC in three UKB phenotypes: “Birth place (north/south)”, “Employment score” and “Waist circumference”, and the phenotype categories into which they were assigned.

### Ancestry pipeline

#### Data Preparation

We merged the following data to generate a painting panel. From POBI[1] we extracted 2039 British samples with county-level UK annotations as described in Leslie et al. From Busby et al.[2] we extracted 511 samples with European country annotations, and a further 4077 samples from POPRES[3]. From the HGDP[4] project we extracted 1035 samples with worldwide, diverse ancestry. From Hapmap[5] we extracted 996 samples from 11 worldwide ancestry annotations. From Pagani et al.[6] we extracted 139 samples with African ancestry annotations, and a further 103 samples from Peterson et al.[7] and 189 from Schlebush et al[8]. This results in 9129 samples annotated into 4 categories: GB (shorthand for Great Britain and Ireland), Europe (Excluding GB), Africa, World (excluding World and Africa). These were available at a subset of all 2,011,414 SNPs based on their genotyping chip, to form a “union SNP list”, used for imputation. Samples in the 1000Genomes[9] or UK10K[10] data were omitted from the painting panel

| Label | Samples | SNPs | Reference |
|---|---|---|---|
| HAPMAP3 | 1184 | 1,383,092 | (International HapMap 3 Consortium, 2010) |
| HGDP | 1035 | 638,797 | (Li et al., 2008) |
| POBI | 2039 | 522,767 | (Leslie et al., 2015) |
| Busby_Europe | 854 | 471,123 | (Busby et al., 2015) |
| POPRES | 4077 | 373,408 | (Nelson et al., 2008) |
| Pagani_Africa | 139 | 816,082 | (Pagani et al., 2012) |
| Peterson_Africa | 103 | 882,127 | (Petersen et al., 2013) |
| Schlebusch_Africa | 200 | 1,914,601 | (Schlebusch et al., 2012) |

We performed basic relatedness pruning with Plink[11] leading to 7775 individuals (2160, 2959,899,1757) individuals from each category. We then performed phasing[12] (using

SHAPEIT2) and imputation[13] (using IMPUTE4) using the UK10K + 1000 Genomes data as a reference panel. 851,948 sites that met a minimum info score threshold of 0.9 in all chips and were not outliers in terms of their pairwise MAF concordance were included in the "draft painting panel".

We then performed chromosome painting with chromopainterv2[14] for each category of individual, using that category as donors (using an individual-level painting). In general, given a set of labelled donor groups chromopainter analyses a sample and produces a "copying vector" measuring the proportion of that samples' genome most similar to each labelled group, using haplotype sharing. For the World panel we also re-added European and African populations as donors. We screened these data for cryptically related individuals by removing individuals sharing too much genome as the most-recent match ("genome length" as a fraction of total in ChromoPainter), with genome fraction limits of (0.035, 0.035, 0.06, 0.05) respectively for (GB, Europe, Africa, World) to account for varying IBD patterns between populations within each region. From each pair of individuals we removed one at random, repeating this process until no individuals were highly related by this measure. This removes (12, 47, 121,57) leading to (2148, 2912,778,1700) individuals respectively.

We then formed "surrogate" and "donor" populations, beginning from sample labels and performed NNLS admixture estimation for each population[15], as described previously. Conceptually "donor" groups form a labelled reference panel, used for constructing copying vectors. Then, "surrogate" individuals from labelled populations often (but not always; see below) matching these donor groups are used to construct summary statistics comprising their chromopainter copying vectors, which are then averaged over individuals from the same surrogate group. (Surrogate individuals must not be included in the donor population to avoid excess copying among relatives; in practice, we can achieve this by "leaving out" this individual from the corresponding donor group[14]) This results in one vector of copying proportions for each surrogate population, precisely capturing their average relatedness to each donor group. Finally, to analyse a sample we run chromopainter against the same set of donors, and fit their resulting copying vector (via NNLS) as a mixture of those constructed for the surrogates, decomposing their ancestry as some set of proportions of these labelled surrogate groups.

To construct our donor and surrogate groups, we impose that all populations are larger than (10,5,5,5) for each category, respectively (10 individuals are required within GB because genetic differences are more subtle), and that all individuals assigned to such a population must have an ancestry proportion from that population, as inferred by NNLS, of at least 50%. Then, we iteratively (requiring only two iterations for convergence) used the following algorithm to identify initially identical donor/surrogate populations:

a) Attempt to split each population using Mclust [16]on their chunklengths, considering up to 5 clusters and proposing clusters that are the maximum *a posteriori* number of populations. We seek splits that maximally separate the samples, exceed the size restriction, and for every individual their recovery in an updated NNLS model is >0.5. In practice this procedure split only very few populations, notably Switzerland, Germany, Italy, and India.
b) Moved individuals whose preferred population was not their surrogate population to their preferred population (i.e. any population where their inferred ancestry was >50%), and remove individuals as surrogates who match no population (for example, because they possess mixed ancestry relative to the other samples).
c) Removed surrogate populations below the size threshold.
d) Where individuals needing to be removed due to steps b-c were in our "removed relatives" list, we checked whether their relatives could now be included instead according to the same criteria.

Finally we checked that surrogate populations remained consistent with their labels, removing populations that had fewer than half their samples from their originally labelled populations. Split populations were assigned new labels based on their genetic affinity to neighbouring populations.

There were a number of instances where we noticed that the above procedure failed to capture important genetic diversity. This occurs when the initial population contains only individuals that our algorithm assigned as relatives, specifically for the following pairs of groups: Native American (Surui and Pima), as well as Lahu and Melanesia, and San and Mbuti. In this case we took the paired populations and used one group (including potential relatives) as a donor, included in the painting panel, and the other as a surrogate population that we used only to estimate the expected “copying proportions”. In this casethe painting should capture populations similar to the surrogate/donor pair. We were also able to use some, potentially admixed individuals as donors but not as surrogates. In Europe this was true of Irish Canadians, whilst in Africa this was true of the “Coloured” label in South Africa, as well as some ASW samples.

This leads to 6632 surrogates (1743, 2648, 752, 1489 respectively) and 6683 donors (1743, 2672, 842, 1426 respectively). We then checked the per-locus painting for these individuals and removed SNPs that showed bias by SNP chip. Specifically we removed SNPs for which the inferred mutation rate (i.e. the mismatch rate to the most similar inferred haplotype, as estimated by the chromopainter miscopying parameter) conditional on the painting was >0.2%. We then checked bias by SNP chip by taking samples with sequence data that were not in our painting or imputation panels from SGDP[17] (Mallick et al., 2016) and EGDP[18] (Pagani et al., 2016). We additionally analysed all public samples (as of November 2017) from OpenSNP (https://opensnp.org/) where individuals had been genotyped on multiple SNP chips or had released sequence data. We then downsampled all sequence data to the list of SNPs for all SNP chips and put them through the phasing and imputation pipeline. Finally, we removed all SNPs that had an imputation discordance rate of >1%. The chips were HAPMAP3, HGDP, POBI, Busby_Europe, POPRES, Pagani_Africa, Peterson_Africa, and Schlebush_Africa, UK Biobank, Ancestry v1, 23andMe v3, 23andMe v4, GSA. This resulted in 677173 SNPs forming the “final painting panel”, for which imputation and phasing results were essentially identical regardless of which SNP chip was used for inference.

| **Dataset** | **# samples** | **# snps** |
|---|---|---|
| UK Biobank (Bycroft et al., 2018) | ~500k | 670,741 |
| FTDNA | 13 | 724,620 |
| 23andMe v3 | 555 | 937,812 |
| 23andMe v4 | 1,043 | 634,617 |
| Ancestry v1 | 176 | 676,010 |
| GSA (Illumina) | 686 | 588,743 |

We then repeated the chromosome painting with the new donor set, and repeat the admixture analysis with these groupings and performed a final iteration of steps b-c in order to ensure that our populations were robust to the exact choice of donor set.

We finally painted UK Biobank individuals who were both in regions of the world that our reference individuals capture poorly, and who have the appropriate ethnicity. Where these populations provided a strong cluster (>50% ancestry recovery for all samples and for a large enough sample as required above) we added these as surrogate by constructing a painting vector summarising typical properties of individuals from these regions. Note however that no UK Biobank samples are used in the running of the pipeline, i.e. as donors. Instead, the surrogate vector provides summary statistics capturing what a typical person with ancestry from Netherlands, Denmark, France, Belgium, Finland, Sweden, Austria, Hundary, Ireland, Algeria, Egypt, Morocco, Libya, Nepal, Myanmar, Bangladesh, Thailand, Vietnam, Philippines, South Korea, Senegal, Liberia, Burundi, Gambia, Cameroon, Angola, Eritrea, Ethiopia, Afghanistan, Somalia, Sudan, Congo, Zambia, Ghana, Zimbabwe and Nigeria all look like in our pipeline. For reporting, we sum ancestry from the 206 surrogate populations where no clear label could be given into the 127 interpretable ancestry labels reported in our results section.

Annotating populations is an art, not a science and reasonable people may have made different decisions based on preferences towards "lumping or splitting" similar populations on a genetic continuum and a need to make ancestry components interpretable via a map. For replicability we therefore include all annotations necessary for this process in the online supplementary materials; specifically the 3 sets of SNP lists, and the donor and surrogate population annotations.

#### Phenotypes in the UKB

We selected 99 representative quantitative phenotypes in the UK Biobank to conduct different analyses. Most of the phenotypes are directly pulled out from the UKB raw data, except the following phenotypes defined algorithmically: Education attainment score (EA), Systolic blood pressure (SBP), Diastolic blood pressure (DBP), pulse pressure (PP) and Estimated glomerular filtration rate (eGFR).

##### Definition of EA

The definition of Education attainment score is based on the UKB field 6138 "Qualification". We use the following criterion to convert categorical variable to quantitative variable:

For participants self-reported as "prefer not to answer", we assigned NA. For the rest of participants, we assigned the score based on the following International Standard Classification for Education (ISCED) definition[1,2]:

$$ES = \begin{cases} 7, if \text{ "}none\ of\ the\ above\text{"} \\ 10, if \text{ "}CSEs\ or\ equivalent\text{" } or \text{ "}O\ levels/GCSEs\ or\ equivalent\text{"} \\ 13, \text{"}A\ levels/AS\ levels\ or\ equivalent\text{"} \\ 15, Other\ professional\ qualifications\ eg: nursing, teaching \\ 19, NVQ\ or\ HND\ or\ HNC\ or\ equivalent \\ 20, College\ or\ University\ degree \end{cases}$$

##### Definition of SBP, DBP and PP

The definition of systolic blood pressure and diastolic blood pressure is based on previous literature[3,4].

The mean SBP and DBP were calculated by using automated blood pressure readings from the first assessment visit (UKBB fields 4080 and 4079) and adjusted for medication usage by adding 15 mmHg and 10 mmHg respectively. If the automated reading was not available, manual measurements (UKBB fields 93 and 94) were used. We further calibrated the blood pressure by setting the readings to missing if the value is more than four standard deviations from the mean. PP was then calculated as SBP minus DBP.

#### Definition of eGFR

eGFR was defined by using four UKBB fields: Creatine (UKBB filed 30700), age at recruiment (UKBB field 21022), sex (UKBB field 31) and Ethnic background (UKBB field 21000). Participants were defined black ethnicity if their ethnic background codes are 4001 (Caribbean), 4002 (African), 4003 (Any other black background). The eGFR was then derived by "CKDEpi.creat" function in R package "nephron" (https://cran.r-project.org/package=nephro ).

### Genetic ancestry in the UKB

#### Visualisation of the inferred genetic ancestry/entropy of UK-born UKB samples

To visualise variation in genetic ancestry, or the entropy of genetic ancestry, across the UK, we used a Gaussian kernel smoothing technique. The genetic ancestry is averaged at each pixel weighted by the Gaussian kernel:

$$\bar{O} = \sum_{i=1}^{N} \frac{e^{-q((x_p-x_i)^2+(y_p-y_i)^2)}}{\sum_{i=1}^{N} e^{-q((x_p-x_i)^2+(y_p-y_i)^2)}} O_i$$

where $\bar{O}$ is the average value (of genetic ancestry or entropy) assigned to a given pixel, $O_i$ is the value for individual *i* ($1 \le i \le N$), N=426,879 is the number of UK-born or Ireland born UKB participants, and $(x_p,y_p)$ and $(x_i,y_i)$ ($1 \le i \le N$) are coordinates of the pixel center and the birthplace of the UKB individual *i* respectively. The weight *q* is pixel-specific and determined by adaptive bandwidth smoothing as follows, so as to keep the "effective" number of individuals per pixel approximately constant at 200:

$$S = \sum_{i=1}^{N} e^{-q((x_p-x_i)^2+(y_p-y_i)^2)}$$

Repeat

q=q/2

$$S = \sum_{i=1}^{N} e^{-q((x_p-x_i)^2+(y_p-y_i)^2)}$$

Until S<200

Return *q*

#### Summary of entropy of UKB participants born in the UK/Ireland

For each participant born in the UK or Republic of Ireland, we calculated an entropy *E* (Methods) to quantify their degree of mixed ancestry (note, this is relative to our assigned population labels). We observed that overall, major cities, and their surrounds, show more mixing on average, possibly reflecting historical migrations to these places. The highest entropy in the UK is for individuals born in London, which also possesses the highest average fraction of ancestry from elsewhere in the world. West Yorkshire has the lowest value of *E,* while lower values are seen for more mountainous and/or remote parts of the UK in general, suggesting that migration into these regions has left less genetic impact. We also note that

Oxfordshire has an interesting signal: a moderate value of *E*, but much lower than the surrounding areas, and the barrier of *E* around Oxfordshire almost coincides the boundary of Oxfordshire (Figure S2).Because some of our donor samples[1] are from Oxfordshire, it is unclear whether this reflects a genuine reduction of mixture relative to surrounding areas or simply more continuous population structure, but regardless, it indicates that county-scale ancestry differences can occur even in regions were previous population structure was not apparent[1].

### Estimation of regional allele frequencies in the UK and worldwide

#### Expectation-maximization based method to estimate regional allele frequency

Highly differentiated alleles across different regions may indicate strong local genetic drift, or act as a signature of natural selection. We used an expectation-maximization (EM) algorithm to estimate regional allele frequencies based on ancestral inference for each individual and region (Methods). We ran the EM on 25,485,700 imputed UKB SNPs, and estimate their allele frequencies across the 23 British-Irish regions, and in 83 worldwide regions, using two different UKB groups (see below).

Using the ancestral inference for each individual, we developed an EM algorithm to infer the regional allele frequencies. We constructed a $2N_s \times N_m$ haplotype matrix by splitting each genotype into two single haplotypes, where $N_s$ is the number of individuals and $N_m$ is the number of SNPs of which the allele frequency will be estimated. In the $n$th EM iteration, the expectation of $h_{ijm}$, the number of copies of the $i$th haplotype derived from ancestral region $j$ at locus $m$, is calculated by (using Bayes theorem):

$$E\left(h_{ijm}\right)^{(n)} = \frac{P\left(h_{ijm} \middle| i \in I_j\right) P(i \in I_j)}{\sum_{j=1}^{Na} P\left(h_{ijm} \middle| i \in I_j\right) P(i \in I_j)} \quad (1)$$

where $N_a$ is the total number of ancestral regions, and $I_j$ is the set of individual haplotype from $j$th population. The probability $P(i \in I_j)$ that individual (haplotype) $i$ is from the $j$th population is equal to the ancestry coefficient $a_{ij}$ inferred from our ancestry pipeline: we assume that conditional on local ancestry being drawn from population $j$, $h_{ijm}$ follows a Bernoulli distribution with parameter $f_{jm}$ , where $f_{jm}$ is the allele frequency of the SNP at the $m$th locus in region $j$. It is these frequencies that we seek to infer. These assumptions lead us on to:

$$E\left(h_{ijm}\right)^{(n)} = \frac{f_{jm(n)}^{h_{ijm}} \left(1 - f_{jm(n)}\right)^{1 - h_{ijm}} a_{ij}}{\sum_{j=1}^{Na} f_{jm(n)}^{h_{ijm}} \left(1 - f_{jm(n)}\right)^{1 - h_{ijm}} a_{ij}}. \quad (2)$$

We then update the allele frequency $f_{jm}$ in the maximization step using the following step (this maximises the full data likelihood where local ancestry is fully known):

$$f_{jm(n+1)} = \frac{\sum_{i=1}^{2N_s} E\left(h_{ijm}\right)^{(n)} I(h_m{=}1)}{\sum_{i=1}^{2N_s} E\left(h_{ijm}\right)^{(n)}} \tag{3}$$

where $I(h_m{=}1)$ is the indicator variable of the mutation type "1" being present on this chromosome at this SNP. We iterate (2) and (3) until convergence is reached, as defined by the frequency reaching stability:

$$\left\| \boldsymbol{f}_{j(n+1)} - \boldsymbol{f}_{j(n)} \right\| < \delta \tag{4}$$

where $\boldsymbol{f}_{j(n)} = (f_{1(n)}, \dots, f_{N_m(n)})$ and "||.||" is the Euclidean norm. We set $\delta$ as 0.001 in our analysis.

We applied this EM to those UKB SNPs with minor allele frequency (MAF) >0.0001 and information score >0.9. We applied the EM in two different subsets of UKB samples, with one subset including all individuals with at least 95% inferred UK+Ireland ancestry, and the remaining samples forming another subset. We call the former subset the "UK+Ireland" subset and the latter the "world subset". For the UK+Ireland subset, we only rescaled the genetic ancestry of the participants such that the sum of their 23 UK+Ireland ancestries equals 1. These rescaled ancestries were then used as input for the EM algorithm to estimate the regional allele frequencies of each SNP in each of the 23 UK + Ireland regions.

For the world subset, we removed all regions with mean ancestry less than 1%, and rescaled the remaining regions such that they again sum up to 1. This resulted in 82 remaining regions whose allele frequencies were estimated in the world subset.

#### Variant rs5743618

A large number of variants exhibiting varying allele frequency across UK were identified. One such is rs5743618 discussed in the main text that has been strongly associated with hay-fever in previous GWAS of European populations. The allele frequency of this variant shows a clear decline from south to north, with the maximum allele frequency difference 24%.

### GWAS analysis

#### Running BOLT-LMM for place of birth (North/South)

BOLT-LMM[1] is a linear mixed model based method which efficiently corrects the population structure and cryptic relatedness in the samples. We applied BOLT-LMM for the trait "Place of birth in UK -North coordinate"(UKB field 129), a trait for which PC-corrected GWAS yields substantial false positives.

We downloaded the autosome genotype call data (UKB field ID 22418) from UKB, and pulled out the same 343,047 white British samples that we used for running AC-corrected and PC-corrected GWAS. We used "*plink1.9*"[2] to conduct variant quality control by filtering out variants in high LD regions, with minor allele frequency less than 0.01, genotype missingness less than 0.05 and Hardy-Weinberg equilibrium p-value less than $1\times10^{-6}$. Afterwards, we used "*plink1.9*" to run LD-pruning with option "--indep-pairwise 1000 100 0.1". Finally, we got in total 142,182 independent chip array SNPs on autosomes, as input for both the "*flashpca*2"[3] and null model building for BOLT-LMM.

20PCs output from "*flashpca*2" together with genotype batch, age, sex, $age^2$, agexsex and $age^2$xsex were used as covariates in the BOLT-LMM analysis. In BOLT-LMM, we also set parameters "bgenMinMAF=0.001" and "bgenMinINFO=03" in line with the setting we ran for "bgenie". "P-BOLT-LMM" column from the output of BOLT-LMM was used to obtain the p-value of BOLT-LMM in the downstream analysis.

#### GWAS with different number of PCs

For the 99 traits we ran AC-corrected GWAS and PC-corrected GWAS using both 20, and 40, PCs while keeping the other covariates the same. After thinning the resulting genome-wide significant variants within 200kb to keep only the most significant variant in each region, we then compared the p-value of independent variants when using 20 or 40 PCs. 48 among the 99 traits possess at least one variant with a p-value change between the 20PC and 40 PC-corrected GWAS that is greater than 100 fold, as summarized in the following table:

| Traits | #variants \|log10(P(pc40)/P(pc20)\|>2 |
|---|---|
| Calcium | 2 |
| Cystatin C | 1 |
| eGFR | 2 |
| Gamma glutamyltransferase | 1 |
| IGF1 | 3 |
| PP | 3 |
| SBP | 5 |
| SHBG | 1 |
| Triglycerides | 1 |
| Urate | 1 |
| Whradjbmi | 1 |
| Place of birth in UK north co ordinate | 17 |
| Place of birth in UK east co ordinate | 14 |
| Sitting height | 10 |
| Home location at assessment east coordinate rounded | 7 |
| Home location at assessment north coordinate rounded | 7 |
| Neuroticism score | 1 |
| Body mass index BMI | 7 |
| Weight | 2 |
| Home location east coordinate rounded | 6 |
| Home location north coordinate rounded | 9 |
| Body fat percentage | 10 |
| Whole body fat mass | 8 |
| Leg fat percentage right | 10 |
| Leg fat mass right | 9 |
| Leg fat percentage left | 14 |
| Leg fat mass left | 5 |
| Arm fat percentage right | 5 |
| Arm fat mass right | 1 |

| | |
|---|---|
| Arm fat free mass right | 1 |
| Arm fat percentage left | 7 |
| Arm fat mass left | 1 |
| Arm fat free mass left | 1 |
| Trunk fat percentage | 9 |
| Trunk fat mass | 8 |
| Trunk fat free mass | 1 |
| Employment score England | 12 |
| Education score England | 5 |
| Housing score England | 1 |
| Forced vital capacity FVC | 4 |
| Forced expiratory volume in second FEV | 6 |
| Peak expiratory flow PEF | 4 |
| Ankle spacing width | 9 |
| Heel bone mineral density BMD | 2 |
| Ankle spacing width left | 5 |
| Ankle spacing width right | 6 |
| Hip circumference | 3 |
| Standing height | 27 |

### Portability of polygenic score across different UKB ancestries

#### Local ancestry inference by "Hapmix"

In order to get the local PRS deconvolution, we selected the 8003 UKB European-African admixed individuals with the sum of (27 UK/European ancestries and 4 African ancestries) larger than 90%. We estimated the local ancestry of each Hapmap3 SNP for each individual using "hapmix"[1] with the default parameter setting. 1000G phase 3 CEU and YRI were chosen as two surrogates representing the European and African ancestries in the reference panel. Output from "hapmix" codes, for each SNP, the joint probability $h(P_1, h_1, P_2, h_2) = Prob(P_1, h_1, P_2, h_2)$ in the *k*th column, where $k = P_1 \times 2^3 + h_1 \times 2^2 + P_2 \times 2^2 + h_1 \times 2^0 + 1$. $P_1$ and $P_2$ are indicator variables that equal 0 (1) if that SNP possesses European (African) ancestry at the first ($P_1$) and second ($P_2$) chromosomal copy (haplotype), and $h_1$ and $h_2$ are also indicator variables now encoding whether the allelic type of this SNP is 1 or 0 at the first and second haplotypes. For the SNP at the *j*th locus for the *i*th individual, $G_{ij}^{EE}$ is the expected "1" allele count, occurring on a background where both haplotype ancestries are European; $G_{ij}^{EA}$ is the expected "1" allele count on the European haplotype, occurring on a background where one of the haplotypes is European and the other is African; Similarly, we defined $G_{ij}^{AA}$ and $G_{ij}^{AE}$ as the expected "1" allele counts when both haplotypes are African, or on the African haplotype when the other ancestral haplotype is European (note that $G_{ij}^{EA}$ and $G_{ij}^{AE}$ are different, reflective of ancestry-specific haplotypic phasing by hapmix). $G_{ij}^{EE}$, $G_{ij}^{EA}$, $G_{ij}^{AA}$ and $G_{ij}^{AE}$ are calculated by summing "hapmix" probabilities as follows:

$G_{ij}^{EE}$ = h(0,1,0,0)+h(0,0,0,1)+h(0,1,0,1)+h(0,1,0,1)

$G_{ij}^{EA}$ = h(0,1,1,0)+h(0,1,1,1)+h(1,0,0,1)+h(1,1,0,1)

$G_{ij}^{AA}$ = h(1,1,1,0)+h(1,0,1,1)+h(1,1,1,1)+h(1,1,1,1)

$G_{ij}^{AE}$ = h(1,1,0,0)+h(1,1,0,1)+h(0,0,1,1)+h(0,1,1,1).

We also calculated $P_{ij}^{EE}$, the probability that both haplotypes have European ancestry, $P_{ij}^{E\backslash A}$, the probability that one haplotype is European and another is African, and $P_{ij}^{AA}$ as the probability that both haplotypes are African, based on the hapmix output.

Finally, total local European ancestry $A_{ij}^{E}$ at the $j$th locus for the $i$th indvidual was calculated as follows: Let $C_{ij}^{E}$ is the number of copies from European ancestry at $j$th locus,

$$A_{ij}^{E} = P_{ij}^{E\backslash A} + 2\, P_{ij}^{EE}.$$

We observed highly consistent genome-wide (average) inference of African ancestry from both “hapmix” and the ancestry pipeline:

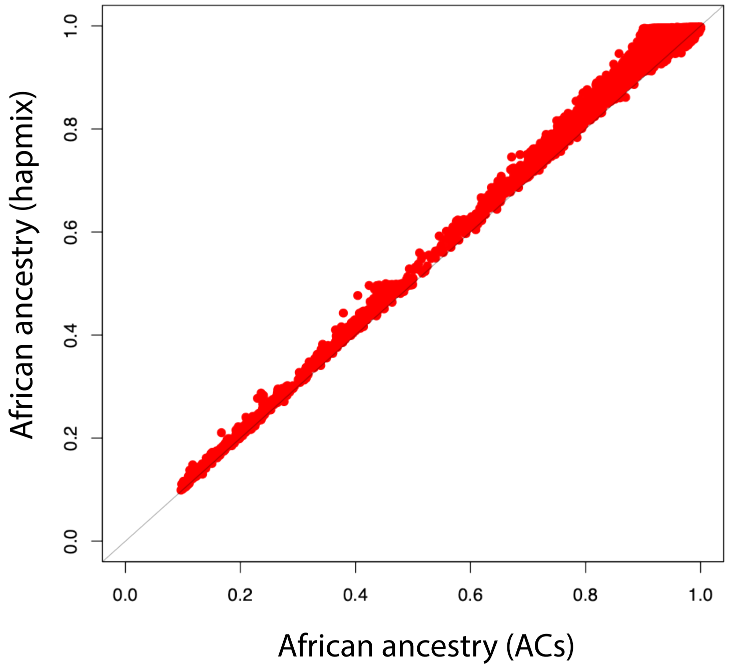

Figure S19. Comparison of inferred African ancestry obtained by ACs (x-axis) and by Hapmix (y-axis)

#### Joint model fitting by accounting for other environment factors

We further investigate if other environment factors, such as birth country or gender, have impact on the portability of polygenic scores. Based on the model in the main text (Methods), we extend the model by adding a new parameter which indicates if a sample individual was born in UK, and fit the model in male and female samples separately. The extended model is as follows:

$$Y_i = I + \left(\beta_{Eu}\mathrm{EPGS}_i + \beta_{Af}\mathrm{APGS}_i\right) \times (1 + \lambda\theta_i + \gamma\zeta_i) + covariates + \varepsilon_i \quad (1)$$

where for individuals $i = 1,2, \dots n$, $Y_i$ is the phenotype, $\varepsilon_i$ is the zero-mean noise, and the parameters to be estimated are the intercept *I*, and $\beta_{Eu}, \beta_{Af}, \lambda, \gamma$. $\mathrm{EPGS}_i$ and $\mathrm{APGS}_i$ are the centralized EPGS and APGS as above, now after regressing out the covariates, and $\theta_i$ is the mean genome-wide European ancestry proportion and $\zeta_i$ is the indicator variable that whether the sample was born in UK (1 if born in UK, 0 otherwise) for individual $i = 1,2, \dots n$.

Based on the model above, we can estimate the effect size of European PGS or African PGS with varying global ancestry for people born in UK or born outside. By stratifying the populations in terms of their gender, we fit this extended model in male and female samples separately (Figure S18) in three scenarios: 1). $\lambda \neq 0$ and $\gamma = 0$; 2). $\lambda = 0$ and $\gamma \neq 0$; 3). $\lambda \neq 0$ and $\gamma \neq 0$. Figure S20 shows the distribution of genetic ancestry stratified by gender and birth places:

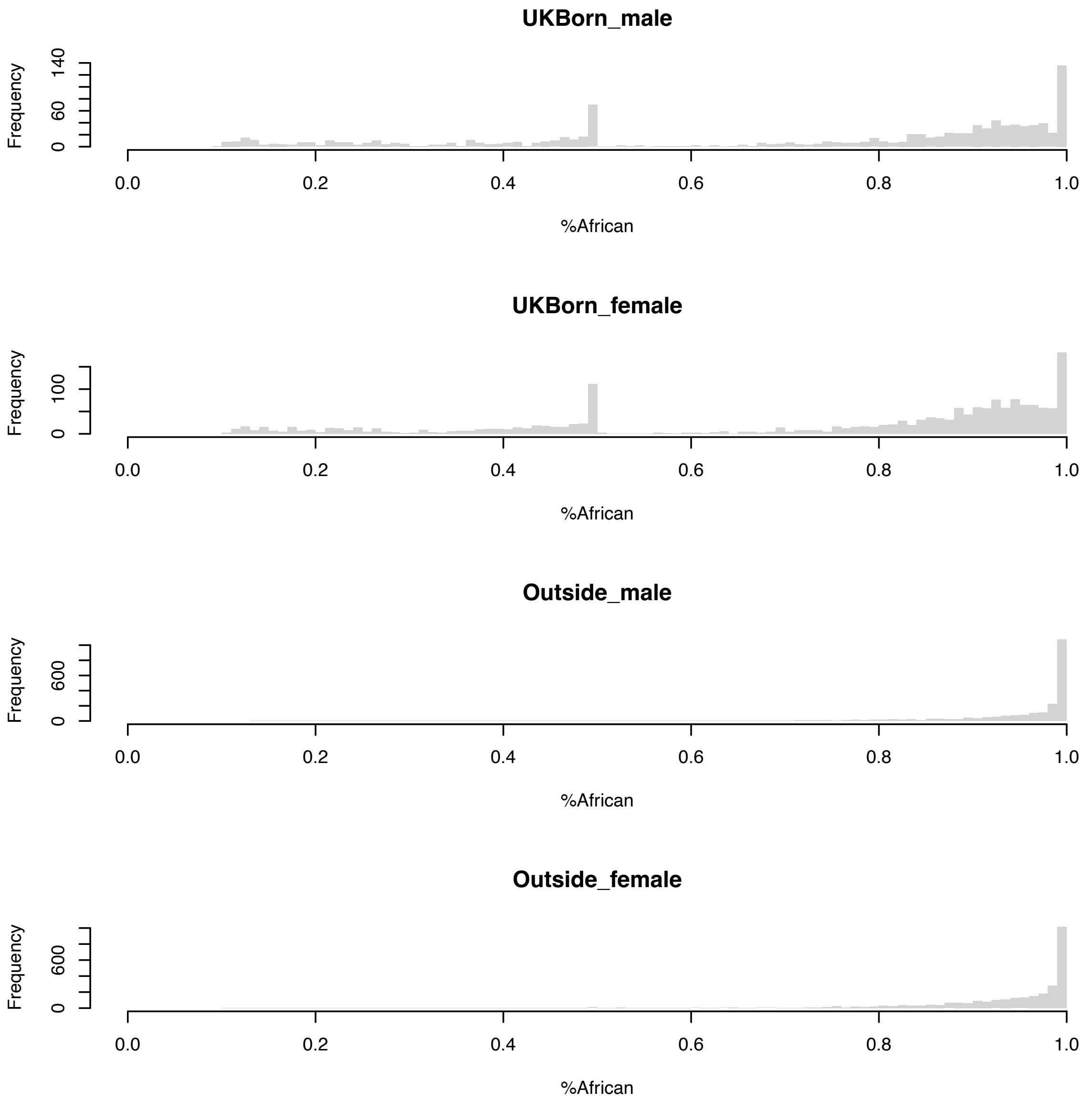

Figure S20. Distribution of African ancestry stratified by gender and whether individuals are born within the UK, or outside the UK.

#### Using trios to conduct ancestry aware phasing for PGSs

We extracted the genotypes of African-ancestry individuals from the raw chip genotype data and conducted quality control by removing the SNPs in regions of high LD, with missing rate larger than 0.05 and with Hardy-Weinberg p-value less than 0.0000001. Using "king"(version 2.2.4)[2] we then identified two trios among the 8003 African-ancestry individuals. For the identified trios, we reran Hapmix with phase specified: heterozygous sites in children are encoded as as "3" if the allele "1" is paternal and "4" if the allele "1" is maternal, and for parents un-transmitted heterozygous allele “1” cases are recoded as “3” and transmitted sites are recoded as “4”.

In trio individuals this results in direct haplotype-based “phased” estimation of the European PGS (EPGS) and African (APGS) of trio individuals across 29 phenotypes, which we compared to their unphased equivalents in the same individuals which were calculated as for the remainder of the 8003 African-ancestry people. To enable joint analysis, we normalised the resulting phased and unphased EPGS and APGS in the following way. Let $\boldsymbol{U}$ be the column vector of unphased EPGS or APGS and $\boldsymbol{P}$ the column vector of phased EPGS or APGS, and $\boldsymbol{C}$ the matrix with **1, age, sex and 31 ACs** as columns. Then we fit the model:

$$\boldsymbol{U} = \boldsymbol{C\beta} + \boldsymbol{\varepsilon}$$

where $\boldsymbol{\varepsilon}$ is the residual. If $\widehat{\boldsymbol{\beta}}$ is the least square estimator of $\boldsymbol{\beta}$, we obtain the residual for both $\boldsymbol{U}$ and $\boldsymbol{P}$:

$$\hat{\boldsymbol{\varepsilon}}_{\boldsymbol{U}} = \boldsymbol{U} - \boldsymbol{C}\widehat{\boldsymbol{\beta}}$$

$$\hat{\boldsymbol{\varepsilon}}_{\boldsymbol{P}} = \boldsymbol{P} - \boldsymbol{C}\widehat{\boldsymbol{\beta}}$$

We then get the mean and variance of residual of $\boldsymbol{U}$ weighted by individual European ancestry $\boldsymbol{A}_E$ for unphased EPGS or African ancestry $\boldsymbol{A}_F = 1 - \boldsymbol{A}_E$ for unphased APRS.

$$\bar{\varepsilon}_U = \frac{\hat{\boldsymbol{\varepsilon}}_{\boldsymbol{U}}^{\boldsymbol{T}} \boldsymbol{A}_E}{\mathbf{1}^{\boldsymbol{T}} \boldsymbol{A}_E}$$

$$var(\hat{\boldsymbol{\varepsilon}}_{\boldsymbol{U}}) = \frac{(\hat{\boldsymbol{\varepsilon}}_{\boldsymbol{U}} - \bar{\varepsilon}_{\boldsymbol{U}} \boldsymbol{A}_E)^{\boldsymbol{T}} (\hat{\boldsymbol{\varepsilon}}_{\boldsymbol{U}} - \bar{\varepsilon}_{\boldsymbol{U}} \boldsymbol{A}_E)}{\mathbf{1}^{\boldsymbol{T}} \boldsymbol{A}_E}$$

For trio individuals, the normalized $\boldsymbol{U}$ and $\boldsymbol{P}$ can be obtained as follows:

$$\widehat{\boldsymbol{U}} = \frac{\hat{\boldsymbol{\varepsilon}}_{\boldsymbol{U}} - \bar{\varepsilon}_{\boldsymbol{U}} \boldsymbol{A}_E}{\sqrt{var(\hat{\boldsymbol{\varepsilon}}_{\boldsymbol{U}})}}$$

$$\widehat{\boldsymbol{P}} = \frac{\hat{\boldsymbol{\varepsilon}}_{\boldsymbol{P}} - \bar{\varepsilon}_{\boldsymbol{U}} \boldsymbol{A}_E}{\sqrt{var(\hat{\boldsymbol{\varepsilon}}_{\boldsymbol{U}})}}.$$

We used these normalised scores to compare across different phenotypes.
