## Supplementary Note 2 for "Leveraging fine-scale population structure reveals conservation in genetic effect sizes between human populations across a range of human phenotypes"

### Estimation of polygenic score (PGS) coefficients in admixed individuals

#### Notation and assumptions

We will use the indexes $i$ to denote an individual in a sample of size $N$, and $j, k$ to index a polymorphism (SNP) within an individual, among $S$ SNPs in total which we will assume to be linearly ordered along the genome. For simplicity, we will consider two ancestries labelled as $A$ and $E$; it is not necessary that these represent "European" and "African" regional ancestries, but in our main application setting this is the case. We will assume we may regard these populations as relatively homogeneous (at least relative to the genetic distinctness of each relative to the other). We also consider the diploid genome as two contiguous haploid stretches, so that the "genotype" at SNP $j$ is either 0 or 1, and there is only one ancestry at each position. Because we are assuming a linear polygenic score in this section, this has no impact on our results, except there are two copies of each SNP coefficient (with identical effect sizes), corresponding to the two chromosomal copies. We assume Hardy-Weinberg equilibrium, so the genotypes on these two chromosomal copies are independent, and their ancestries are also independent given the appropriate genome-wide ancestry for an individual.

When there is admixture, ancestry in general varies along the genome and across individuals. Individual $i$ has genome-wide population $A$ mean ancestry $\theta_i$. At SNP $j$, define an indicator variable $I_{ij}$ which is 1 if individual $i$ possesses ancestry drawn from $A$ at SNP $j$ and 0 if their ancestry is instead $E$ at SNP $j$; we will sometimes neglect the $i$ subscript while considering a single individual.

The genotype of an individual at a particular SNP is denoted by $g_{ij}$. We will also utilise mean genotypes at SNP $j$; these are given by $\bar{g}_j^A$ and $\bar{g}_j^E$ to represent (twice) the mean frequencies in populations $E$ and $A$ respectively. It will also be helpful to consider the mean genotype of individual $i$ at SNP $j$; this is denoted by $\bar{g}_{ij} = \theta_i \bar{g}_j^A + (1 - \theta_i)\bar{g}_j^E$.

We will denote the confounders/coregressors for individual $i$ as a vector $C_i$ (forming a matrix $C$ across individuals) whose first entry $C_{i1} = \theta_i$, the ancestry. Let the observed phenotype value for individual $i$ be $Y_i$

SNPs impact the trait of interest. As is standard, we will assume linear additive effects across SNPs such that SNP $j$ has true effect $\beta_j$. We will not require that these effects have any particular distribution, other than that the trait is polygenic. We also assume that one or more genome-wide association studies (GWAS) has been conducted, resulting in an estimated effect size $\hat{\beta}_j$ for each SNP $j$. We will assume that individual $i$ (or their relatives) were not included in these GWAS, so we may regard this estimate as independent of the genetic data of this individual (this is true for our real-data applications).

We will additionally make two further simplifying assumptions about the use of PGS. First, we will assume coregressors (or confounders) corrected for in the estimation procedure (e.g. age, gender, genotyping centre, etc) are independent of genotypes featuring in the PGS, conditional on genome-wide ancestry; we will assume, as is standard in genome-wide association studies, that genome-wide ancestry is also corrected for as a confounder. Second, we will assume ancestry segments within individual are “large” in the following sense: if two SNPs are in different ancestry segments, then we may assume no linkage disequilibrium (LD) between them in either founder population, or across founder populations. Informally this assumption means that ancestry segments extend further than LD, and it is likely to be reasonably well satisfied for groups admixing within the past few hundred generations.

### Models

Here, we will derive a procedure for testing whether the $\beta_j$ terms measuring effect sizes at SNPs depend on genome-wide ancestry, indicating gene-environment or gene-gene interactions. To do this, we derive two quantities that capture the behaviour of the PGS under a null model when the $\beta_j$ coefficients *do not* depend on genome-wide (or local) ancestry, i.e. there is no variability in underlying effect sizes across populations. Estimation of these quantities in practice then allows testing for departures (if any) from this simple null model. In the final section, we will generalise to the setting where underlying effect sizes can vary between populations.

Specifically, we consider the setting where our phenotype

$$Y_i = \alpha + \sum_{j=1}^{S} \beta_j g_{ij} + \mu C_i + \epsilon_i,$$

where $\alpha$ and the vector $\mu$ are coefficients expressing the impacts of a constant term, and the confounders, on the phenotype for individual $i$. Here $\epsilon_i$ is the error, which is often assumed to be normally distributed. Under our null model, note that the terms $\beta_j$ do not depend on ancestry and so are identical across all individuals, whatever their admixture proportions. Note also that we have a term $\mu_1\theta_i$ in the above which accounts for a possible effect of genome-wide ancestry on $Y_i$.

We would like to use this model, but unfortunately the values of $\beta_j$ are not known to us, and so instead we replace these terms with their estimates, yielding the model:

$$Y_i = \alpha + \beta \sum_{j=1}^{S} \hat{\beta}_j g_{ij} + \mu C_i + \epsilon_i. \tag{1}$$

Here, note that the sum in the second term is the estimated PGS. We must also introduce a term $\beta$ in front of this term. This accounts for the fact that it is likely our PGS may, for example, systematically estimate effect sizes (winner's curse effect), include SNPs not truly associated with the trait, or include variants in LD, each of which will result in the variance of the PGS estimate not completely capturing variance of the true PGS. Accordingly, we expect $0 < \beta < 1$ in most cases. We also require that different individuals are approximately unrelated, so their genotypes can be regarded as independent, once their ancestries are taken into account. However, we are agnostic as to how the coefficients $\hat{\beta}_j$ are obtained (apart from this occurring in an independent sample). Specifically, we require no particular assumptions about what approach is used to correct for LD, or penalisation of the coefficients etc - these will impact the particular value of $\beta$ that is appropriate, but do not otherwise impact our results. The levels of LD within populations are also not important, apart from the assumption that ancestry segments extend further than within-population LD in admixed individuals. The below analysis only requires that the above linear model holds; in other words, that on average $Y_i$ increases linearly with the estimated PGS value. We believe this condition to hold reasonably well for many, or most, complex traits after appropriate rescaling.

### Overview of approach

In the standard approach to quantifying the variance explained by the PGS, the squared correlation between the phenotype and PGS is obtained by fitting the least-squares estimators $\hat{\beta}$ of $\beta$, and of the other parameters, in the model (1) and then calculating the mean squared residual variance relative to the variance when $\beta = 0$. The least squares estimator represents the mle of the parameters under the assumption that $\epsilon_i$ is normally distributed, and is unbiased and consistent provided the mean of the residuals $\epsilon_i$ are zero for each individual, and the residual variances are bounded.

More generally, for a linear model the true value of a parameter vector $\beta$ is that value which minimises the expected mean squared residuals over individuals. It is also the mean of $\hat{\beta}$. In a large iid sample, the central limit theorem means this mean squared residual is asymptotically equal to $\mathbb{E}\left(\epsilon_i^2\right)$ for a single individual $i$, now defining the expectation as taken over both data $Y_i$, and genotypes $X_i$ (which can be regarded as randomised across the sample). We may regard this as a *definition* of $\beta$, and will use this fact throughout this supplement. It is equivalent to the statement that if we have a combined matrix $X$ of predictors:

$$\mathbb{E}_{X,Y}\left(X^T Y\right) = \mathbb{E}_X(X^T X)\beta.$$

### Homogeneous populations

First, we consider a sample of individuals each drawn from population $E$. In this case, the true value of $\beta$ is $\beta_E$ say and this minimises $\mathbb{E}\left(\epsilon_i^2\right)$ under the model

$$Y_i = \alpha + \beta_E \sum_{j=1}^{S} \hat{\beta}_j g_{ij} + \gamma C_i + \epsilon_i.$$

Note that because the model fit includes a constant $\alpha$, this is identical to fitting the following model which will be more convenient:

$$Y_i = \alpha + \beta_E \sum_{j=1}^{S} \hat{\beta}_j \left( g_{ij} - \bar{g}_j^E \right) + \mu C_i + \epsilon_i.$$

By the same argument, for a sample of individuals drawn from population $A$ we have a true value $\beta_A$ say which minimises $\mathbb{E}\left(\epsilon_i^2\right)$ in the model:

$$Y_i = \alpha + \beta_A \sum_{j=1}^{S} \hat{\beta}_j \left( g_{ij} - \bar{g}_j^A \right) + \mu C_i + \epsilon_i.$$

In the setting where the coefficients $\hat{\beta}_j$ are estimated using GWAS in populations similar to $E$, we expect $0 < \beta_A < \beta_E < 1$. This is because due to regional linkage disequilibrium differences the estimated PGS is expected to on average be a better model for the true PGS in populations genetically closer to those used for the discovery. We note that using e.g. $g_{ij} - \bar{g}_j^E$ in place of $g_{ij}$ means using *mean-centred* genotypes i.e. a mean-centered PGS.

In a homogeneous population, as we have seen above the choice of whether to centre genotypes is arbitrary, and so researchers often use either form interchangeably. However, we will show that the choice has important consequences in the case of an admixed population; and so we strongly recommend the use of mean-centring, conditional on *local* ancestry, in this case.

### Full model incorporating admixed individuals

For this section, we will consider the setting where the polygenic score shows no gene-gene or gene-environment effects that are correlated with ancestry, so that the following true phenotypic model (with the true PGS) applies:

$$Y_i = \alpha^o + \sum_{j=1}^{S} \beta_j \left( g_{ij} - \bar{g}_{ij} \right) + \mu^o C_i + \epsilon_i^o. \qquad (2)$$

We consider this as the underlying generating equation for the phenotype, and examine properties of *estimated* PGS below. Here we mean centre the genotypes by subtracting $\bar{g}_{ij}$. The resulting sum over these mean genotypes depends only on the genome-wide ancestry $\theta_i$ of individual $i$, so may be absorbed into the $\alpha^o$ and $\mu_1^o$ coefficients. Therefore this model remains equivalent to that with non-centred genotypes. Note that this model is assumed to apply across individuals with differing proportions $\theta_i$ of ancestry $A$, allowing for linear effects of ancestry on the phenotype in the model. Here, $\epsilon_i^o$ is the "true" (optimal) residual if we knew the model fully; $\epsilon_i^o$ is independent of the other random variables, with mean zero and finite variance $\sigma^2$; similarly $\mu^o$ etc represent the other true underlying parameters.

Introducing the indicator of ancestry $A$ at SNP $j$, $I_j$, it is natural to consider fitting the model based on applying the separate models for $A$ and $E$ local ancestry from the previous section, for the relevant genome pieces:

$$Y_i = \alpha + \beta_A \sum_{j=1}^{S} \hat{\beta}_j I_j \left(g_{ij} - \bar{g}_j^A\right) + \beta_E \sum_{j=1}^{S} \hat{\beta}_j (1 - I_j) \left(g_{ij} - \bar{g}_j^E\right) + \mu C_i + \epsilon_i. \quad (3)$$

If we define the ancestry $A$ PGS

$$APGS = \sum_{j=1}^{S} \hat{\beta}_j I_j \left(g_{ij} - \bar{g}_j^A\right)$$

(the estimated mean-centred PGS, summing only over those parts of the genome with ancestry $A$), and similarly for $EPGS$, then we can express this linear model as

$$Y \sim EPGS + APGS + coregressors.$$

Thus, this model can be fit from real data, provided $EPGS$ and $APGS$ can be obtained. We note that equations (2) and (3) provide two ways of writing $Y_i$, which must agree for this model to be self-consistent. In particular, individuals have varying genome-wide ancestry $\theta_i$. Intuitively, one might expect that the mean coefficients of $EPGS$ and $APGS$ will equal their values $\beta_E$, $\beta_A$ from the previous section, and *not* depend on $\theta_i$. If this is the case then equation (3) is well defined, in the sense that we obtain the same coefficients across distinct samples, regardless of their ancestry proportions - for example across samples binned according to $\theta_i$. Below we show that under our no-interaction null, this is the case, *provided* we use mean-centred scores as above. Essentially, the proof extends the (assumed) fact that the true parameters $\beta_j$ are unchanging across individuals $i$ to show the *inferred* parameters also do not change with $\theta_i$. We note in passing that by our initial assumptions, conditional on genome-wide ancestry the terms $EPGS$ and $APGS$ are each uncorrelated with (in fact independent of) the coregressors (and uncorrelated with this ancestry itself), so in large samples the estimated coefficients for the coregressors are approximately independent of those for the PGS terms. Note also that the mean-centring works differently for the two ancestries $A$ and $E$, due to different frequencies of mutations in each group.

For convenience, we will consider a slightly more general model of the following form, which will allow us to consider the alternative scenario of no mean centring:

$$\begin{aligned} Y_i =& \alpha + \beta_A \sum_{j=1}^{S} \hat{\beta}_j I_j \left(g_{ij} - \bar{g}_j^A\right) + \beta_E \sum_{j=1}^{S} \hat{\beta}_j (1 - I_j) \left(g_{ij} - \bar{g}_j^E\right) \\ &+ \sum_{j=1}^{S} \left[\gamma_j I_j + \lambda_j (1 - I_j)\right] + \mu C_i + \epsilon_i. \end{aligned} \quad (4)$$

where the $\gamma_j, \lambda_j$ are non-independent coefficients such that given genome-wide mean ancestry $\theta_i$, $\theta_i \gamma_j + (1-\theta_i)\lambda_j = 0$; in other words, the mean of the terms in the third sum is zero. In our main model $\gamma_j \equiv \lambda_j \equiv 0$; allowing non-zero terms allows us to include an offset in the different parts of the score.

Next, we will consider properties of the true parameters in Equation (4); equivalent to the mean estimates of those parameters using least-squares estimation.

#### Expected squared residuals

From the previous discussion, upon fitting Equation (4), the least-squares fit results in unbiased parameter estimates, whose expectations minimise the mean sum of squares of residuals, as we average across phenotype values $Y$ and predictors $X$. It is then sufficient to consider this mean squared residual for a single individual $i$, with particular ancestry $\theta_i$.In fact we may write using equations (2) and (4):

$$\begin{aligned}
&\alpha^o + \sum_{j=1}^{S} \beta_j \left(g_{ij} - \bar{g}_{ij}\right) + \mu^o C_i + \epsilon_i^o \\
&= \alpha + \beta_A \sum_{j=1}^{S} \hat{\beta}_j I_j \left(g_{ij} - \bar{g}_j^A\right) + \beta_E \sum_{j=1}^{S} \hat{\beta}_j (1-I_j) \left(g_{ij} - \bar{g}_j^E\right) \\
&\quad + \sum_{j=1}^{S} \left[\gamma_j I_j + \lambda_j (1-I_j)\right] + \mu C_i + \epsilon_i .
\end{aligned}$$

From above the true parameters in equation (4) minimise $\mathbb{E}\left(\epsilon_i^2\right)$ (for any $i$). Solving the last equation for $\epsilon_i$, note $C_i$ is uncorrelated with the centered genotypes that appear in the other terms, which also have mean zero. Therefore the contribution from the non-$\beta$ parameters to $\mathbb{E}\left(\epsilon_i^2\right)$ is simply additive, and vanishes if and only if $\alpha = \alpha^o$ and $\mu = \mu^o$, so these parameter values are unchanged relative to equation (2), i.e. we estimate the correct underlying values for these parameters. Upon removing these vanishing terms, we obtain

$$\begin{aligned}
\sum_{j=1}^{S} \beta_j \left(g_{ij} - \bar{g}_{ij}\right) + \epsilon_i^o = \beta_A \sum_{j=1}^{S} \hat{\beta}_j I_j \left(g_{ij} - \bar{g}_j^A\right) + \beta_E \sum_{j=1}^{S} \hat{\beta}_j (1-I_j) \left(g_{ij} - \bar{g}_j^E\right) \\
+ \sum_{j=1}^{S} \left[\gamma_j I_j + \lambda_j (1-I_j)\right] + \epsilon_i .
\end{aligned}$$

Finally a rearrangement yields

$$
\begin{aligned}
\epsilon_i &= \sum_{j=1}^{S} \beta_j \left(g_{ij} - \bar{g}_{ij}\right) - \beta_A \sum_{j=1}^{S} \hat{\beta}_j I_j \left(g_{ij} - \bar{g}_j^A\right) - \beta_E \sum_{j=1}^{S} \hat{\beta}_j (1 - I_j) \left(g_{ij} - \bar{g}_j^E\right) \\
&\quad - \sum_{j=1}^{S} \left[\gamma_j I_j + \lambda_j (1 - I_j)\right] + \epsilon_i^o \\
&= \left( \sum_{j=1}^{S} I_j \left[\beta_j - \beta_A \hat{\beta}_j\right] (g_{ij} - \bar{g}_j^A) \right) && \text{(I)} \\
&\quad + \left( \sum_{j=1}^{S} (1 - I_j) \left[\beta_j - \beta_E \hat{\beta}_j\right] (g_{ij} - \bar{g}_j^E) \right) && \text{(II)} \\
&\quad + \sum_{j=1}^{S} I_j \left(\beta_j [\bar{g}_j^A - \bar{g}_{ij}] - \gamma_j\right) + (1 - I_j) \left(\beta_j [\bar{g}_j^E - \bar{g}_{ij}] - \lambda_j\right) && \text{(III)} \\
&\quad + \epsilon_i^o. && \text{(IV)}
\end{aligned}
$$

We consider the properties of the labelled specific terms on the RHS as we average over the genotype $g_{ij}$ and ancestry $I_j$ (each varying across sampled individuals). It will be convenient to write the overall genotype

$$g_{ij} = I_j g_{ij}^A + (1 - I_j) g_{ij}^E$$

where $g_{ij}^A$ represents a random draw from the genome of a single ancestry $A$ individual for each $j$, and similarly for $g_{ij}^E$. SNPs in different ancestry segments must come from different ancestors, so $g_{ij}^A$ and $g_{ij}^E$ are independent, and independent of $I_j$ also. This allows us to regard $g_{ij}^A$ as existing even if $I_j = 0$, since it does not impact the observed genotype, and similarly for $g_{ij}^E$: this will simplify taking expectations. In particular, within term (I) we may write $I_j(g_{ij} - \bar{g}_j^A) = I_j(g_{ij}^A - \bar{g}_j^A)$ which involves only independent random variables, and similarly for term (II).

Using this, term (I) and term (II) immediately have mean 0, and expected product 0, since for example $\mathbb{E}(g_{ij}^A) = \bar{g}_j^A$ and by independence of $g_{ij}^A$ and $g_{ij}^E$. Moreover the expected products of term (III) and terms (I) and (II) are zero. To see this, first take the expectation of the product of e.g. term (I) and term (III) conditional on the value of $I_j$ for each $j$. Term (III) is conditionally constant, while terms (I) and (II) have mean zero after averaging over $g_{ij}^A$ and $g_{ij}^E$. Thus the conditional expected product of term (I) and term (III) is zero, so the unconditional expectation is also. Finally, term (IV) is independent of the other terms so also has expected product with the other terms of zero. Thus, to calculate $\mathbb{E}\left(\epsilon_i^2\right)$ we must only sum the mean squares of terms (I)-(IV). The expectation of squared term (IV) is $\sigma^2$ and does not depend on the other parameters, so may be ignored in what follows. Now considering term (I):

$$
\begin{aligned}
&\mathbb{E}\left[\left(\sum_{j=1}^{S} I_j \left[\beta_j - \beta_A \hat{\beta}_j\right] (g_{ij}^A - \bar{g}_j^A)\right)^2\right] \\
&= \sum_{j=1}^{S}\sum_{k=1}^{S} \left[\beta_j - \beta_A \hat{\beta}_j\right] \left[\beta_k - \beta_A \hat{\beta}_k\right] \mathbb{E}\left(I_j I_k (g_{ij}^A - \bar{g}_j^A)(g_{ik}^A - \bar{g}_k^A)\right) \\
&= \sum_{j=1}^{S}\sum_{k=1}^{S} \left[\beta_j - \beta_A \hat{\beta}_j\right] \left[\beta_k - \beta_A \hat{\beta}_k\right] \mathbb{E}\left(I_j (g_{ij}^A - \bar{g}_j^A)(g_{ik}^A - \bar{g}_k^A)\right) \\
&= \theta_i \sum_{j=1}^{S}\sum_{k=1}^{S} \left[\beta_j - \beta_A \hat{\beta}_j\right] \left[\beta_k - \beta_A \hat{\beta}_k\right] \mathbb{E}\left((g_{ij}^A - \bar{g}_j^A)(g_{ik}^A - \bar{g}_k^A)\right) \\
&= \theta_i \mathbb{E}\left[\left(\sum_{j=1}^{S} \left[\beta_j - \beta_A \hat{\beta}_j\right] (g_{ij}^A - \bar{g}_j^A)\right)^2\right].
\end{aligned}
$$

Here the third line follows from our assumption that ancestry changes more slowly than LD, so that either $I_k = I_j$ or otherwise $\mathbb{E}\left((g_{ij}^A - \bar{g}_j^A)(g_{ik}^A - \bar{g}_k^A)\right) = 0$ because SNPs $k$ and $j$ are not in LD. The final expression depends only on $\beta_A$ among parameters to be fit. By inspection, the $\beta_A$ value maximising this expression is unchanging with $\theta_i$ provided the $\beta_j$ terms remain fixed, because the distribution of $g_{ik}^A$ is just that of genotypes of individuals from population $A$. When $\theta_i = 1$, the entire genome is from population $A$ so our model reduces to the homogeneous case. Therefore the optimal $\beta_A$ coefficient for this term is that for the non-admixed case where we simply regress the phenotype of a sample drawn from population $A$ on the centred APGS.

By the same argument replacing ancestry $A$ with ancestry $E$, term (II) has mean square which depends only on $\beta_E$ and not on other parameters to be fit. It is minimised by choosing the same $\beta_E$ coefficient for this term as for a non-admixed case, where we regress the phenotype of a sample drawn from population $E$ on the centred EPGS. Finally, note that by construction term (III) has mean zero and depends only on $I_j$, the local ancestry. Then term (III) equals:

$$
\sum_{j=1}^{S} (1-\theta_i) I_j \left(\beta_j[\bar{g}_j^A - \bar{g}_j^E] - \gamma_j\right) - \theta_i (1 - I_j)\left(\beta_j[\bar{g}_j^A - \bar{g}_j^E] - \lambda_j\right)
$$

(simplifying further does not much aid interpretability). In principle, we can make this term vanish by setting $\gamma_j = \lambda_j = \beta_j[\bar{g}_j^A - \bar{g}_j^E]$. Unfortunately, this requires complete knowledge of the polygenic score which we do not possess! Further, we cannot easily estimate this term from GWAS data by conducting only a GWAS in a population similar to population $E$, for example a GWAS of UKB White British samples, if $E$ represents European ancestry. That is because we do not know causal SNPs, preventing us from estimating their frequencies in population $A$ accurately.

#### The case $\gamma_j, \lambda_j = 0$

In our current approach, we simply set $\gamma_j = \lambda_j = 0$. This results in some value for the expected mean square of term (III) depending on $\theta_i$, but *not* on $\beta_A$ or $\beta_E$. Therefore $\mathbb{E}\left(\epsilon_i^2\right)$ depends on these parameters only through terms (I), (II), involving $\beta_A$, $\beta_B$ respectively. Thus, from above, $\mathbb{E}\left(\epsilon_i^2\right)$ is maximised by the same values of these coefficients as estimated from samples of individuals from populations $A$ only and $E$ only respectively. The "true" parameter values in the model then equal these values, which large-sample estimates converge to under our modelling assumptions (we describe variance estimation below). This is our main result: because we may estimate such coefficients in practice, testing them for equality with those estimated from non-admixed samples, or other admixed samples, represents a test of shared effect sizes (below, we extend to consider properties of these coefficients if effect sizes differ between groups).

Term (III) is stochastic (depending on local ancestry) and has mean zero, but non-zero mean square which depends on $i$ through $\theta_i$. This can be thought of as a random noise, additional to the noise term $\epsilon^o$ in the phenotype. Therefore although this does not asymptotically bias parameter estimates, it will add additional noise in their estimation. This term almost vanishes in three useful cases: for individuals where only a small amount of ancestry is from population $A$, or from population $E$, or for individuals where one parent is entirely from population $A$ and the other almost entirely from population $E$. In other cases, estimation of this term seems a useful topic for future research, and we note it contains information about genetic differences between populations $A$ and $E$.

#### Problems and pitfalls using non-centred polygenic scores

A tempting approach, rather than using the mean-centred polygenic scores $APGS$ and $EPGS$ as described above, is to simply use their non-centred versions where we do not subtract $\bar{g}_j^A$ and similar terms for SNP $j$. Note that we can immediately view this as equivalent to setting $\gamma_j = \beta_A \hat{\beta}_j \left[\bar{g}_j^A - \bar{g}_{ij}\right]$ and $\lambda_j = \beta_E \hat{\beta}_j \left[\bar{g}_j^E - \bar{g}_{ij}\right]$. From above, term (III) in our equations above then becomes

$$\sum_{j=1}^{S} (1-\theta_i) I_j \left[\beta_j - \beta_A \hat{\beta}_j\right] \left[\bar{g}_j^A - \bar{g}_j^E\right] - \theta_i (1 - I_j) \left[\beta_j - \beta_E \hat{\beta}_j\right] \left[\bar{g}_j^A - \bar{g}_j^E\right].$$

Unless our estimates of the regression coefficients $\beta_j$ are perfect, this is non-zero, and moreover this term depends on both our coefficients of interest $\beta_A$ and $\beta_E$. In general there is no reason to believe the values of these parameters optimising terms (I) and (II) respectively will also optimise this term; indeed unlike these terms, the optimal values in general depend on $\theta_i$. As a result, under our null model estimates for $\beta_A$ and $\beta_E$ using these non-centred polygenic scores must minimise the sum of terms (I-III), so need not equal those for samples drawn from the non-admixed populations $E$ and $A$. This means that only the *centred* scores are suitable for testing for effect differences between groups. In practice, when we used non-centred scores in individuals of mixed European and African ancestry, we indeed observe differences in parameter estimates. We observe reduced estimated values for $\beta_{\text{Europe}}$ in individuals with high African ancestry, and increased values for $\beta_{\text{Africa}}$ in individuals with high European ancestry.

#### Parameter estimation in real samples and simulated data

In the UK Biobank (UKB) data, we analyse 8003 individuals whose ancestry is mainly derived from a mixture of sub-Saharan African ancestry (referred to henceforth as ancestry $A$), and European (especially British and Irish) ancestry (ancestry $E$). In practice, we run Hapmix to obtain realised versions of the haplotype-specific genotypes, jointly with the inferred ancestries $I_{ij}$ at SNP $j$ for individual $i$. Previous work (Price et al. 2009)[1] has shown that this yields highly accurate ancestry estimates. Analysing data for trios suggests as expected that there is high accuracy of ancestry imputation in these data (Supplementary Figure S13) and by comparing to direct phasing, that the ancestry $A$ and ancestry $E$ polygenic scores are accurately estimated using Hapmix (Supplementary Figure S14), for a range of traits and across individuals.

To construct the $EPGS$ and $APGS$ terms, we use $PGS$ $\beta$ estimates constructed from the UKB analysis of individuals with British and Irish ancestry (Methods); note this population is closely related to group $E$. We also require the mean frequencies $\bar{g}_j^E$ and $\bar{g}_j^A$ of each SNP in the ancestral groups. We may obtain genome-wide ancestry estimates $\theta_i$ for each individual by taking the proportion of their genome inferred by Hapmix to come from population $A$ (i.e. the African-like ancestry). Then noting that we may model

$$\mathbb{E}\left(g_{ij}\right) = \theta_i \bar{g}_j^A + (1 - \theta_i)\bar{g}_j^E$$

we fit a linear regression of the observed genotypes $g_{ij}$ on the left hand side against the values $\theta_i$ and use the resulting coefficients to estimate the two required means. Given the large sample size, these estimates are expected to closely match the true SNP frequencies for the specific admixing groups (they also correlate strongly with the frequencies of the same variants in 1000G populations; data not shown).

Having obtained $EPGS$ and $APGS$, for different phenotypes and sets of individuals we first fit the model of equation (3), using our ACs, age, and gender as coregressors. We separately fitted models for each of 29 phenotypes (Main text; Figure 4) within UKB, binning individuals according to their genome-wide ancestry. This supplement demonstrates that if the PGS operates consistently (i.e. with conserved true effect sizes) across ancestries, then we estimate the same mean values for $\beta_E$ and $\beta_A$ asymptotically for each bin, and $\beta_E$ will also approximately equal the value obtained directly for the large number of individuals within UKB of British and Irish ancestry, which we estimate as $\beta_{Obs.eu}$.

To obtain confidence intervals for the underlying parameter values, we used bootstrapping (1000 bootstrapped sample replicates). To reduce noise, we either combined all 8003 samples for a single trait (Figure 4d,e: estimators $\beta^{All}_{Eu}$ and $\beta^{All}_{Af}$ for $\beta_E$, $\beta_A$ and their bootstrapped confidence intervals are plotted) or compared bins using average values across all traits (Figure 4b,c). We observe highly consistent estimates for $\beta_E$ in all bins ($EPGS$ terms) and similarly consistent estimators for $\beta_A$ ($APGS$), though these values are much lower.

To formally test whether $\beta_A$, $\beta_E$ change with ancestry, indicating varying underlying effect sizes in admixed individuals, we fit a model allowing for an interaction of their effect sizes with genome-wide ancestry. Specifically, we generalised equation (3) to fit the model

$$Y_i = (\beta_A APGS + \beta_E EPGS)(1 + \lambda\theta_i) + coregressors$$

where the coregressors are as before. If $\lambda = 0$ this reduces to the equation (3) case, but if $\lambda \neq 0$ then this model allows the effect sizes $\beta_j$ of SNPs to be altered on average as mean ancestry varies. This could capture effects due to gene-gene interactions and/or gene-environment interactions, which correlate with ancestry. We numerically obtain the least-squares estimates of the parameters of this model, and then evaluate confidence intervals via bootstraps as above. The $\lambda = 0$ case corresponds to the null of no variation in underlying effect sizes with ancestry, and so we accept the null if zero lies within the confidence interval for $\lambda$. Although we perform a 2-sided test, note that if interactions do exist, we expect to observe $\hat{\lambda} < 0$ in most cases when the GWAS producing the PGS was performed in population $E$ or a close surrogate, for the same reason we expect $\beta_A < \beta_E$ in this case: the SNPs we identify will tend to have large (mean) effects in population $E$.

Note that $\beta^{Eu}_{Eu} = \hat{\beta_E}$ measures the estimated mean increase in the phenotype per unit increase in $EPGS$ for individuals with 100% population $E$ ancestry, while $\beta^{Af}_{Eu} = \hat{\beta_E}(1 + \hat{\lambda})$ can be interpreted as the corresponding term for individuals with (near) 100% population $A$ ancestry; these terms, along with their corresponding $\hat{\beta_A}$ equivalents, are shown in Figure 4d, and ratios of terms are shown in Figure 4e. If the 95% confidence interval for $\beta^{Af}_{Eu}/\beta_{Obs.eu}$ includes 1, then there is no evidence for the underlying PGS coefficients varying between individuals of different ancestry, and this is true for almost all traits we analysed. Contrastingly, we see strong evidence for essentially all traits that $\beta^{Af}_{Af}/\beta^{Af}_{Eu} < 1$, so that differences in local LD and mutation frequencies impact all PGS examined, with an overall average estimate of 35% for this ratio indicating a huge disparity between groups.

Finally, although Hapmix appears accurate in our testing, we also tested robustness of our estimates by masking uncertain or short ancestry segments (likely to be more prone to errors) from our estimation procedure and re-estimating coefficients; the results were virtually unchanged (Figure S21). Errors in ancestry estimation or assignment of mutations to particular backgrounds would result in ancestry $A$ mutations being assigned to $E$, and vice-versa. Therefore, we would expect such errors to result in more similar estimated coefficients for $\beta_E$ and $\beta_A$ than is truly the case, by mixing their contributions. Since we estimate coefficients for $\beta_E$ that closely resemble those for non-admixed individuals from a population similar to $E$, rather than moving towards $\beta_A$, our results seem unlikely to be strongly impacted by such errors.

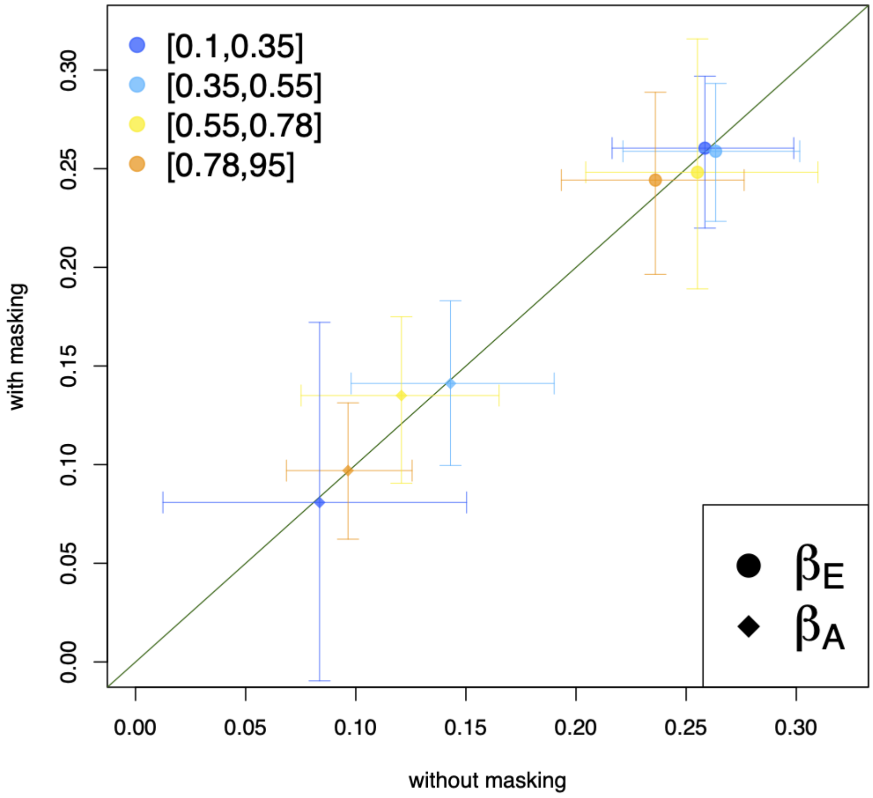


Figure S21: Comparison of $\beta_E$ and $\beta_A$ between without (x-axis) and with masking (y-axis) the short ancestry segments. Colours represent different African ancestry bins (same as Figure 3b)

We analysed data for simulated phenotypes in exactly the same way as described above, i.e. matching the real data analyses.

#### Parameter estimation in real samples: variable effect sizes between populations

Consider the setting where population $J$ is now an admixed population of interest, while we conduct an initial GWAS, and construct a PGS, in a population genetically identical to a second population $I$. Suppose these populations now have different effect sizes at SNPs, for example due to gene-gene or gene-environment interactions. For a particular causal SNP $j$ with effect sizes $\beta_j^I$ and $\beta_j^J$ in each group, initially we assume these are normally distributed with mean zero and covariance matrix

$$\text{Var}\begin{pmatrix} \beta_j^I \\ \beta_j^J \end{pmatrix} = \begin{pmatrix} \sigma^2 & \sigma^2\rho \\ \sigma^2\rho & \sigma^2 \end{pmatrix}$$

We previously considered the setting $\rho = 1$ (without imposing any normality assumption). We now consider properties of the estimators $\hat{\beta}_E$ and $\hat{\beta}_A$, if the two ancestries for population $J$ are $A$ and $E$, for any $\rho$. Conditional on the value of $\beta_I$ we have by properties of the Normal distribution that $\mathbb{E}\left(\beta_j^J \mid \beta_j^I\right) = \rho\beta_j^I$. This is always reduced, because $\rho \leq 1$. Considering the estimators, we may still consider equations 2 and 4 and the subsequent derivation and properties of terms (I-IV) remain unchanged, if the true underlying PGS coefficient $\beta_j = \beta_j^J$ now represents the causal effect size of SNP $j$ in population $J$. By normality we can represent $\beta_j^J = \rho\beta_j^I + \sqrt{1-\rho^2}\sigma Z_j$ where for each $j$, $\beta_j^I$, the causal effect size in group $I$, and $Z_j \sim N(0,1)$, representing the perturbation of causal effects, are independent. From our earlier derivation, the large-sample estimator of $\beta_E$ minimises term (II) and hence the following:

$$\mathbb{E}\left[\left(\sum_{j=1}^{S}\left[\beta_j^J - \beta_E\hat{\beta}_j\right](g_{ij}^E - \bar{g}_j^E)\right)^2\right]$$

$$= \mathbb{E}\left[\left(\sum_{j=1}^{S}\left[\rho\beta_j^I + \sqrt{1-\rho^2}\sigma Z_j - \beta_E\hat{\beta}_j\right](g_{ij}^E - \bar{g}_j^E)\right)^2\right]$$

$$= \rho^2\mathbb{E}\left[\left(\sum_{j=1}^{S}\left[\beta_j^I - \frac{\beta_E}{\rho}\hat{\beta}_j\right](g_{ij}^E - \bar{g}_j^E)\right)^2\right] + \sigma^2\left(1-\rho^2\right)\mathbb{E}\left[\left(\sum_{j=1}^{S} Z_j(g_{ij}^E - \bar{g}_j^E)\right)^2\right].$$

By inspection, the second term is independent of $\beta_E$ while the first is identical after reparameterizing to the corresponding term for population $I$. If population $I$ represents European ancestry, this is optimised by the parameter $\beta_{Obs.eu}$. Hence the optimal value $\beta_E$ is $\beta_E = \rho\beta_{Obs.eu}$. Therefore, we can interpret the estimated relative coefficient $\hat{\beta}_E/\hat{\beta}_{Obs.eu}$ as an estimate of the correlation in effect sizes, at causal SNPs, between European-ancestry and mixed African/European ancestry individuals (within any frequency bin).

We can relax the assumption of normality in this derivation, provided that the expectation of the true PGS in population $J$ conditional on the true PGS in population $I$ *and* the estimated PGS in population $I$, is equal to $\rho$ times the true PGS in population $I$. (This immediately holds under normality.) That is, $\rho$ is the increase in mean trait value in the admixed group for a genetic perturbation causing an increase in mean trait value of 1 in population $I$, and the estimated PGS does not give information about population $J$ not available from the *true* PGS in population $I$. We expect this second condition to normally hold in practice provided population $I$ closely approximates the GWAS population used to produce the estimated PGS. We therefore define $\rho$ in general to equal the limiting value of $\hat{\beta}_E/\hat{\beta}_{Obs.eu}$. $\rho$ equals the correlation in true genetic risks, if the variances in effect sizes are equal in $I$ and $J$. In applications, even if effect size variances differ between two populations, the ratio $\hat{\beta}_E/\hat{\beta}_{Obs.eu}$ estimates the mean increase in mean trait value in the admixed group for a genetic perturbation causing an increase in mean trait value of 1 in population $E$, which still has a useful interpretation. In particular if the admixed group is more responsive to genetic perturbations overall it is possible to obtain $\rho > 1$, even in the presence of incomplete correlation in effect sizes between groups; note that this means the PGS is more predictive of changes in the trait in the admixed group.

In our real analysis we never observe significant evidence $\rho > 1$ for any trait (main text), and overall $\rho \approx 1$ explains our data well, and is close to the average value across traits. This is expected under a null model of largely shared (causal) effect sizes between groups and across traits. Under a model of differing causal effect sizes, observing $\rho \approx 1$ would require African-ancestry individuals to be systematically more responsive to genetic perturbations (higher variance), which we are not aware of evidence for from the literature, in just such a way as to cancel out the incomplete correlations; this seems very non-parsimonious across a diverse group of traits, so we regard the data as supporting shared causal effect sizes, and this in turn allows $\rho$ to be interpreted as a correlation.

In our simulations incorporating either complete or incomplete correlation of effect sizes, we used the above covariance model $\beta_j^J = \rho\beta_j^I + \sqrt{1-\rho^2}\sigma Z_j$, simulated equal underlying variances and estimated $\rho$ using $\hat{\beta}_E/\hat{\beta}_{Obs.eu}$ to obtain values shown (Figure 4b, Supplementary Figures S15 and S16).
